# Telomerase-independent survival leads to a mosaic of complex subtelomere rearrangements in *Chlamydomonas reinhardtii*

**DOI:** 10.1101/2023.01.23.525193

**Authors:** Frédéric Chaux, Nicolas Agier, Clotilde Garrido, Gilles Fischer, Stephan Eberhard, Zhou Xu

## Abstract

Telomeres and subtelomeres, the genomic regions located at chromosome extremities, are essential for genome stability in eukaryotes. In the absence of the canonical maintenance mechanism provided by telomerase, telomere shortening induces genome instability. The landscape of the ensuing genome rearrangements is not accessible by short-read sequencing. Here, we leverage Oxford Nanopore Technologies long-read sequencing to survey the extensive repertoire of genome rearrangements in telomerase mutants of the model green microalga *Chlamydomonas reinhardtii*. In telomerase mutant strains grown for ∼700 generations, most chromosome extremities were capped by short telomere sequences that were either recruited *de novo* from other loci or maintained in a telomerase-independent manner. Other extremities did not end with telomeres but only with repeated subtelomeric sequences. The subtelomeric elements, including rDNA, were massively rearranged and involved in breakage-fusion-bridge cycles, translocations, recombinations and chromosome circularization. These events were established progressively over time and displayed heterogeneity at the subpopulation level. New telomere-capped extremities composed of sequences originating from more internal genomic regions were associated with high DNA methylation, suggesting that *de novo* heterochromatin formation contributes to restore chromosome end stability in *C. reinhardtii*. The diversity of alternative strategies to maintain chromosome integrity and the variety of rearrangements found in telomerase mutants are remarkable and illustrate genome plasticity at short timescales.

## Introduction

Protection of chromosome extremities is essential for genome integrity. For most eukaryotes, it is achieved by repeated DNA sequences called telomeres and by telomere-bound factors, which collectively prevent chromosome ends from being processed as DNA damage^1,2^. Telomeres shorten with each round of replication due to the end replication problem and are in general maintained by telomerase, a dedicated reverse transcriptase able to elongate telomeres *de novo*. In its absence, some telomeres eventually reach a critical length that triggers replicative senescence, an arrested state induced by the DNA damage checkpoint. Replicative senescence was shown in some species to increase genome instability due to repair attempts and bypass of the checkpoint arrest through the adaptation to DNA damage process^3-11^. In senescent cells that eventually escape cell cycle arrest, such as some precursor cancer cells, telomeres become dysfunctional and induce further genomic instabilities, a phenomenon termed telomere crisis^12,13^.

The absence of telomerase therefore generates genome instabilities that stem from telomeres and take many shapes: point mutations, deletions/insertions, translocations, aneuploidy, duplications and even more dramatic rearrangements, such as chromothripsis during telomere crisis^7^. The precise molecular mechanisms underlying these alterations are not all well understood but often involve classical and alternative non-homologous end-joining (c- and a-NHEJ), homology-directed repair (HDR), including homologous recombination and break-induced replication (BIR), together with missegregation of chromosomes, breakage-fusion-bridge (BFB) cycles and other dynamic phenomena that act in cascades over multiple cell divisions^3-5,7,13-17^.

Subtelomeres are the genomic regions adjacent to telomeres and often contain families of paralogous genes or pseudo-genes, ribosomal DNA (rDNA) arrays, transposable elements and other repeated sequences^18-24^. Subtelomeres are often involved in telomere-associated rearrangements due to their repetitive nature promoting HDR, replication fork stalling and template switching (FoSTeS), and BIR^19,25-32^. Consistently, subtelomeres evolve rapidly even in closely related species and within species^18,27,33-35^. In some species, in the absence of telomerase, telomeres can be stabilized using the alternative lengthening of telomeres (ALT) pathway, which depends on homologous recombination and uses repeated sequences found in telomeres and subtelomeres as substrates^36-39^.

Genome alterations, especially structural variations (SVs), initiated by telomere shortening and dysfunction, despite been widely studied in different models including cancer cells, have been difficult to map exhaustively due to the complex nature of the rearrangements and the frequent involvement of repeated sequences such as the ones found in subtelomeres^13,40^. Only recently were long-read sequencing technologies used to enable the resolution of complex rearrangements at chromosome extremities, in response to telomere shortening and dysfunction in *Caenorhabditis elegans* and *Saccharomyces cerevisiae* ^41-43^.

We recently provided a comprehensive map of all 34 subtelomeres of the unicellular green alga *Chlamydomonas reinhardtii* (17 chromosomes as haploid, 111 Mb)^24^. All contain arrays of repeated elements, the most common being the Sultan element, present in 31 out of the 34 chromosome extremities in a haploid strain, arranged in tandem repeats of up to 46 elements. The basic Sultan element has a length of ∼850 bp and forms class A subtelomeres. The Sultan element of class B subtelomeres contains additional insertions. Next to most Sultan arrays (29 out of 31), a sequence of ∼500 bp called Spacer is unique to each subtelomere and may serve as promoter for downstream non-coding RNA genes. The 3 remaining subtelomeres are entirely composed of rDNA, for a total of approximately 350 copies corresponding to ∼3 Mb. Two other repeated elements, called Suber and Subtile, were found next to Sultan elements at 3 subtelomeres, called class C. The Suber element, initially named pTANC^44^, contains the most abundant interstitial telomere sequence (ITS) of the genome. We previously found experimental evidence of telomere-associated genome rearrangements potentially involving subtelomeres in telomerase mutants of *C. reinhardtii*, correlated with long-term survival^45^. Indeed, although some telomerase-negative mutant subclones underwent senescence-induced cell death, many managed to survive telomerase absence and must have therefore found a solution to maintain and protect telomeres. In this work, using long-read Oxford Nanopore Technologies (“Nanopore”) sequencing able to traverse large repeated regions, we investigated genome instability in telomerase mutants in *C. reinhardtii* and provide an exhaustive view of the landscape of genome rearrangements. Our data show that subtelomeres are massively altered by rearrangements within and between subtelomeres, involving amplification and deletion of repeated elements (or even entire subtelomeres), fusions, BFB cycles, and other events of high complexity. Rearrangements occur progressively over time and create a mosaic population of cells which undergo competition and selection. Telomeres are maintained at a majority of extremities in a telomerase-independent manner and possibly promote methylation of adjacent DNA to help stabilize chromosome extremities.

## Results

### Long-read Nanopore sequencing of prolonged cultures of telomerase mutants

To investigate the genome rearrangements induced by the long-term absence of telomerase, we cultivated two different telomerase mutant strains, *tel-m1* and *tel-m2* (Fig. 1A). The two mutant strains were obtained from a library of random insertion mutants^46^ and the insertion of the paromomycin resistance gene in the RNA-binding domain (*tel-m1*) and catalytic domain (*tel-m2*) of *CrTERT*, the gene encoding the catalytic subunit of telomerase, was confirmed previously^45^. We estimated that the mutants underwent ∼700 generations between creation of the library of mutants in the Jonikas lab and collection of the samples called *tel-m1-1* and *tel-m2-1*, based on the procedure of transformation, propagation and freezing described in^46^ and our subsequent passages. Additionally, for *tel-m1*, we collected an earlier sample cultivated for ∼250 generations named *tel-m1-0*, which corresponded to the earliest timepoint we could reasonably obtain upon reception of the strain in our lab. The wild-type strain used to make the library of mutants, CC-4355, was also cultivated upon reception alongside the mutants, for at least 450 generations.

**Figure 1.**
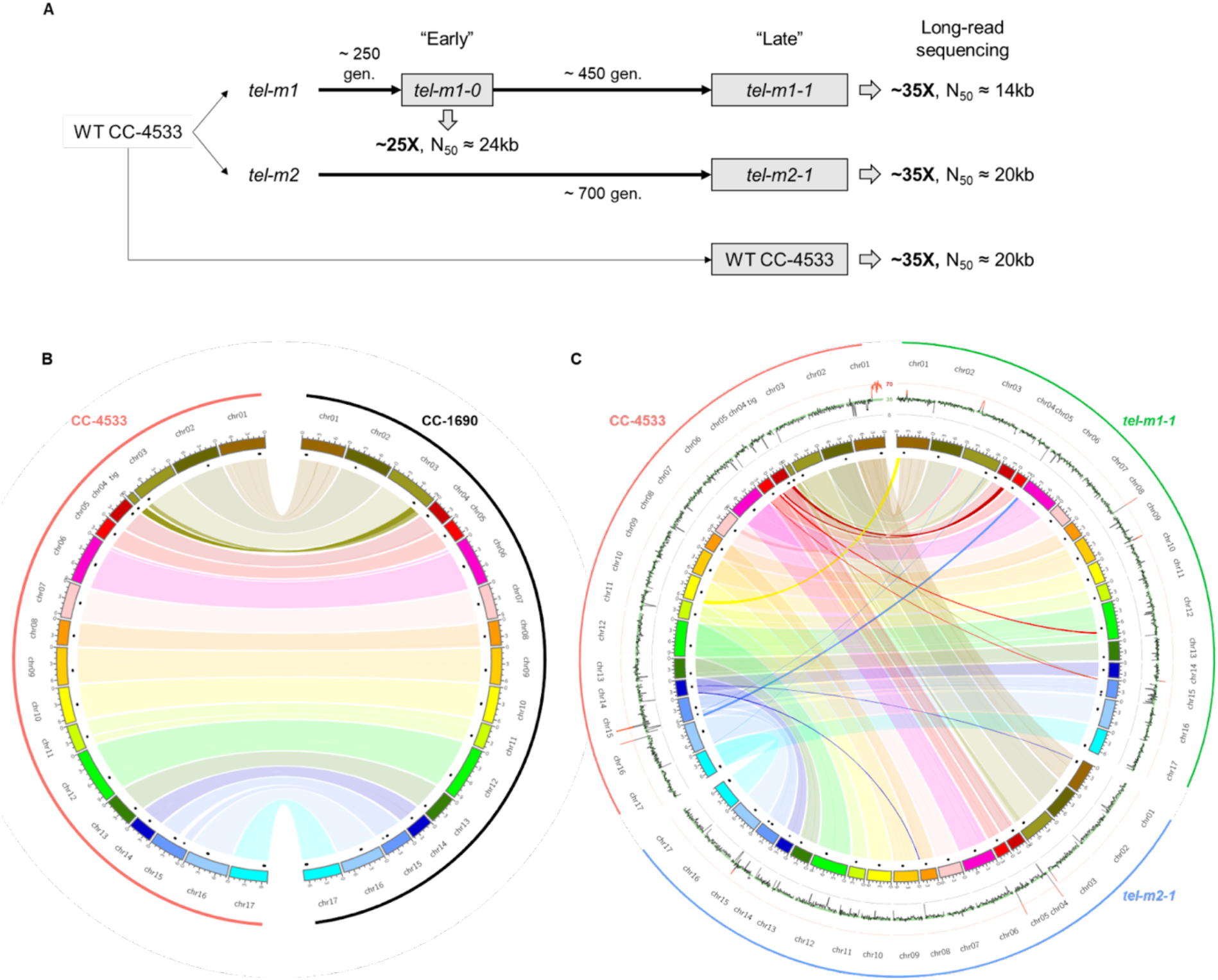
Independent genome assemblies of CC-4533 and telomerase mutants *tel-m1-1* and *tel-m2-1* reveal few re-arrangements at the genome assembly level. (A) Experimental setup and main sequencing output. (B-C) Circos plot^48^ of (B) chromosome scaffolds from the long-term culture of CC-4533 as compared to CC-1690 and to *tel-m1-1* and *tel-m2-1*, shown as colored boxes, with black marks for centromere clusters (composed of the *Zepp-L1* repeat). Links connect homologous blocks, colored according to CC-4533 chromosomes. Outer plot displays sequencing depth, averaged over 100 kb regions, with 0X, 35X and 70X grid lines shown in black, green and red, respectively. Depth over 50X is highlighted in red and genomic position is indicated in megabases.

We optimized a CTAB/phenol/chloroform purification protocol followed by a size selection step to extract and obtain long DNA molecules^47^. We then sequenced the genomic DNA of these 4 samples by long-read Nanopore sequencing and obtained 4 sets of high-quality reads of N50 > 14 kb and depth > 25X (Supp. Table ST1). We previously proved that long reads allowed the accurate reconstruction of the highly repeated subtelomere structures in *C. reinhardtii*^24^, which is instrumental for the analysis of rearrangements potentially implicating telomeres, subtelomeres and other repeated elements.

The long-read datasets were processed by 3 assemblers, allowing us to generate *de novo* genome assemblies of CC-4533 and each telomerase mutant (Fig. 1B and C). We reached large chromosome-scale scaffolds similar to the most contiguous reference genome of CC-1690^49^, to which we compared our new assembly of CC-4533. Sequencing depth (Fig. 1C, outer circles) indicated a partial duplication of chromosome 1 in CC-4533 not present in previous sequencing^50^, suggesting that a duplicated mini-chromosome 1 emerged during the long-term culture in our laboratory, a phenomenon that is quite common since similar duplications were observed in other laboratory strains^51^. The copy number variation (CNV) on chromosome 15 and 16 were due to frequently misassembled regions known to contain repeated sequences^52^. No other chromosome-scale difference was found between CC-4533 and CC-1690 (Fig. 1B). When compared to CC-4533, *tel-m1-1* displayed a number of large-scale SVs and CNVs, corresponding to translocations and duplications of chromosome extremities (Fig. 1C). The other mutant, *tel-m2-1*, showed fewer large-scale rearrangements in the assembly, but CNVs were detected at several locations, suggesting genome alterations. We then used the SV caller MUM&Co^53^ to detect insertions, deletions, duplications, inversions and translocations compared to CC-1690 and we found similar numbers for CC-4533 and the telomerase mutants assemblies, with about half of SVs shared by all 3 strains (n = 370 out of a total of 746) (Supp. Fig. S1A and B). The assemblies of the telomerase mutants displayed a moderate number of unique SVs (n = 52 and 49 for *tel-m1-1* and *tel-m2-1*, respectively), involving sequences of median size < 600 bp.

However, we found that the quality of read mapping to subtelomeres was lower than those corresponding to the core genome both in CC-4533 and in the telomerase mutants (Supp. Fig. S1C), presumably because the repeated elements found at subtelomeres present a challenge for genome assembly even using long-read sequencing data^54^. Consistently, the assembled genomes reached only 25, 2 and 19 telomeres in CC-4533, *tel-m1-1* and *tel-m2-1*, respectively. We thus suspected that the assembly-level analysis would miss genome rearrangements affecting chromosome extremities. Additionally, assemblers were not designed to handle genome heterogeneity at the subpopulation level, which is likely to be the case in telomerase mutants. We therefore directly looked for genome alterations at the level of the sequencing reads instead of the assemblies in the rest of this work.

### Telomere shortening and loss in telomerase mutants

We were first interested in investigating changes related to telomeres. To detect all telomere sequences at the level of individual reads, we developed a new method called TeloReader that scores 8-mers with respect to their level of identity to any of the canonical 8-mers (TTTTAGGG/CCCTAAAA and circular permutations) in sliding windows and uses thresholds for the average score to find the boundaries (see Methods section). TeloReader allowed us to detect all telomere sequences of at least 16 bp, whether they were at a chromosome extremity (terminal) or not (interstitial telomere sequence, ITS), in the read datasets.

We measured the length of all terminal telomere sequences in the wild-type and the mutants reads and found shorter telomeres in the mutants (CC-4533: 293 ± 126 bp, *tel-m1-0*: 167 ± 109 bp, *tel-m1-1*: 162 ± 120 bp, *tel-m2-1*: 181 ± 88 bp; Fig. 2A). Telomeres were already short in the *tel-m1-0* sample and their length did not further decrease in the *tel-m1-1* sample, suggesting the stabilization of telomere length by telomerase-independent mechanisms or *de novo* telomere formation after complete loss. Because we suspected that the aggregate length distribution measurement would mask telomere length dynamics at each extremity, we assigned the reads containing a terminal telomere to a specific chromosome arm based on the presence of specific and unique Sultan elements in the reads and measured telomere length distribution for each Sultan-associated extremity (Supp. Fig. S2). Out of the 27 studied class A and B extremities, 8 showed telomeres further shortening between *tel-m1-0* and *tel-m1-1*, with large variations in the rate of shortening. Interestingly, they were stabilized or even slightly increased in length between *tel-m1-0* and *tel-m1-1* in 16 cases, indicating a telomerase-independent mechanism of telomere maintenance or *de novo* formation. In the remaining 3 extremities, the very low or even absence of reads in *tel-m1-0* and/or *tel-m1-1* prevented comparison and suggested the loss of the entire telomere or subtelomere sequence. Telomere length distribution at each extremity also displayed variations in *tel-m2-1*.

**Figure 2.**
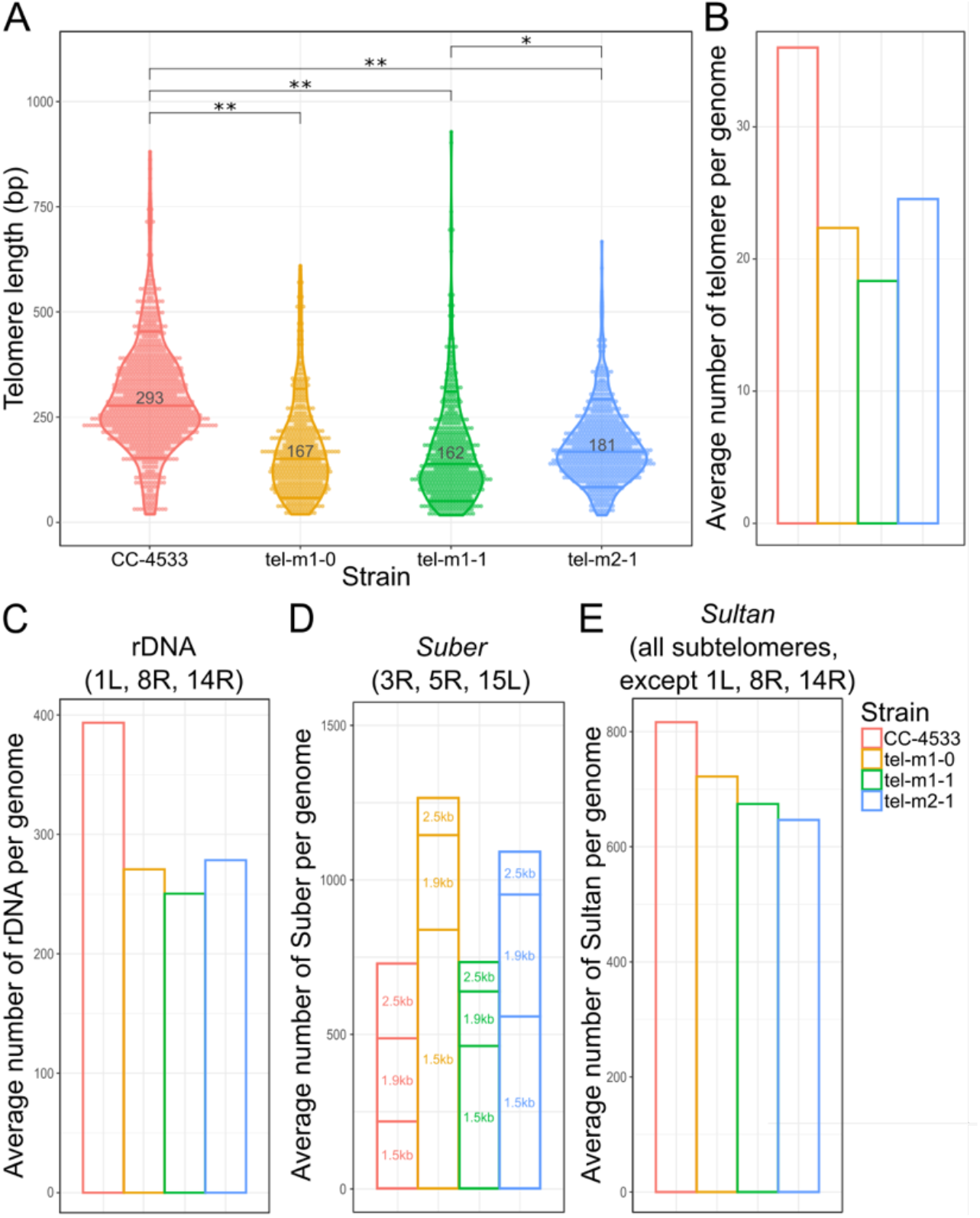
Telomere length distribution and CNVs of subtelomeric elements. (A) Telomere length distribution of terminal telomere sequences detected by TeloReader from the reads dataset in CC-4533 and 3 telomerase mutant samples. * and ** indicate statistically significant difference with p-value = 6.8 × 10^−9^ and p-value < 2.2 × 10^−16^, respectively, using a non-parametric Kruskal-Wallis test. (B) Number of telomeres in reads normalized by the sequencing depth. (C-E) CNVs for repeated elements normally found at subtelomeres including (C) the rDNA, the Suber elements subdivided into 3 types according to their length (“1.5kb”, “1.9kb” and “2.5kb”), and (E) the Sultan elements.

To test if a subset of telomeres were completely lost, we computed the total number of telomere-containing reads normalized to the sequencing depth of the nuclear genome and found that the average number of telomeres per cell was smaller in the mutants: 22 in *tel-m1-0*, 18 in *tel-m1-1* and 25 in *tel-m2-1* vs 36 in CC-4533 (which has an extra mini-chromosome 1 in addition to the 17 chromosomes) (Fig. 2B).

Overall, telomeres shortened as expected in telomerase mutants but our data revealed the heterogeneity of shortening pattern at each specific telomere and suggested that telomeres were maintained or newly formed in *tel-m1-1* as compared to *tel-m1-0* in a telomerase-independent manner. In the next sections, we investigated changes and chromosomal rearrangements occurring at subtelomeric regions.

### Copy number variations of subtelomeric elements in telomerase mutants

As a first insight into the stability of repeated subtelomeric elements (rDNA, Suber and Sultan), we computed their CNVs in telomerase mutants compared to CC-4533.

We used the rDNA sequence as a query to find all reads containing rDNA. We observed an overall decrease of 31%, 36% and 29% in rDNA copy number in *tel-m1-0, tel-m1-1* and *tel-m2-1*, respectively, compared to CC-4533 (Fig. 2C). We also found that, while in the wild-type, it was exclusively found at 3 subtelomeres as expected^24^, rDNA could be mapped at several additional regions of the genome in telomerase mutants, implicating rDNA in genome rearrangements (see below). The two major rDNA clusters at subtelomeres 8R and 14R likely lost a large fraction of their rDNA copies.

The same analysis was performed with the Suber element, which is found as 3 main subtypes of sizes ∼1.5 kb, ∼1.9 kb and ∼2.5 kb^24,44^, and showed changes specific to each subtype (Fig. 2D). The copy number of the 1.5 kb element was greatly increased in the telomerase mutants compared to wild-type, especially in *tel-m1-0*, in contrast to the 2.5 kb element which decreased in copy number in all mutants. The number of 1.9-kb elements only decreased in *tel-m1-1*. Given the organization of Subers in arrays containing only one subtype^24^, the different changes in copy number depending on the subtype likely reflected duplicated, deleted and potentially rearranged arrays.

Finally, the Sultan element, the most abundant and widespread repeated element in subtelomeres, was decreased in copy number in both telomerase mutants (Fig. 2E). In the next sections, we investigate in more detail what this decrease entails at the level of each chromosome extremity. Overall, all the main subtelomeric repeated elements showed CNVs in the telomerase mutants, indicating that subtelomeres are involved in telomere-induced chromosome rearrangements.

### Contraction, erosion and expansion of Sultan arrays

To analyze subtelomere structure, we looked for the presence of the Spacer element in all reads because this element is present as a single copy in 29 out of 34 subtelomeres in the wild-type strain, with sufficient sequence differences to uniquely identify the corresponding subtelomere^24^. Reads containing a Spacer element were thus grouped according to their Spacer (Supp. Dataset 1).

While for all Spacers, reads from CC-4533 were consistent with a subtelomeric structure identical or near-identical to the one we inferred from the genome assembly of CC-1690^24^ (Fig. 3, Supp. Dataset 1), reads from the telomerase mutants revealed many differences compared to CC-4533, including a striking variety of rearrangements (Table 1 and Supp. Dataset 1).

**Figure 3.**
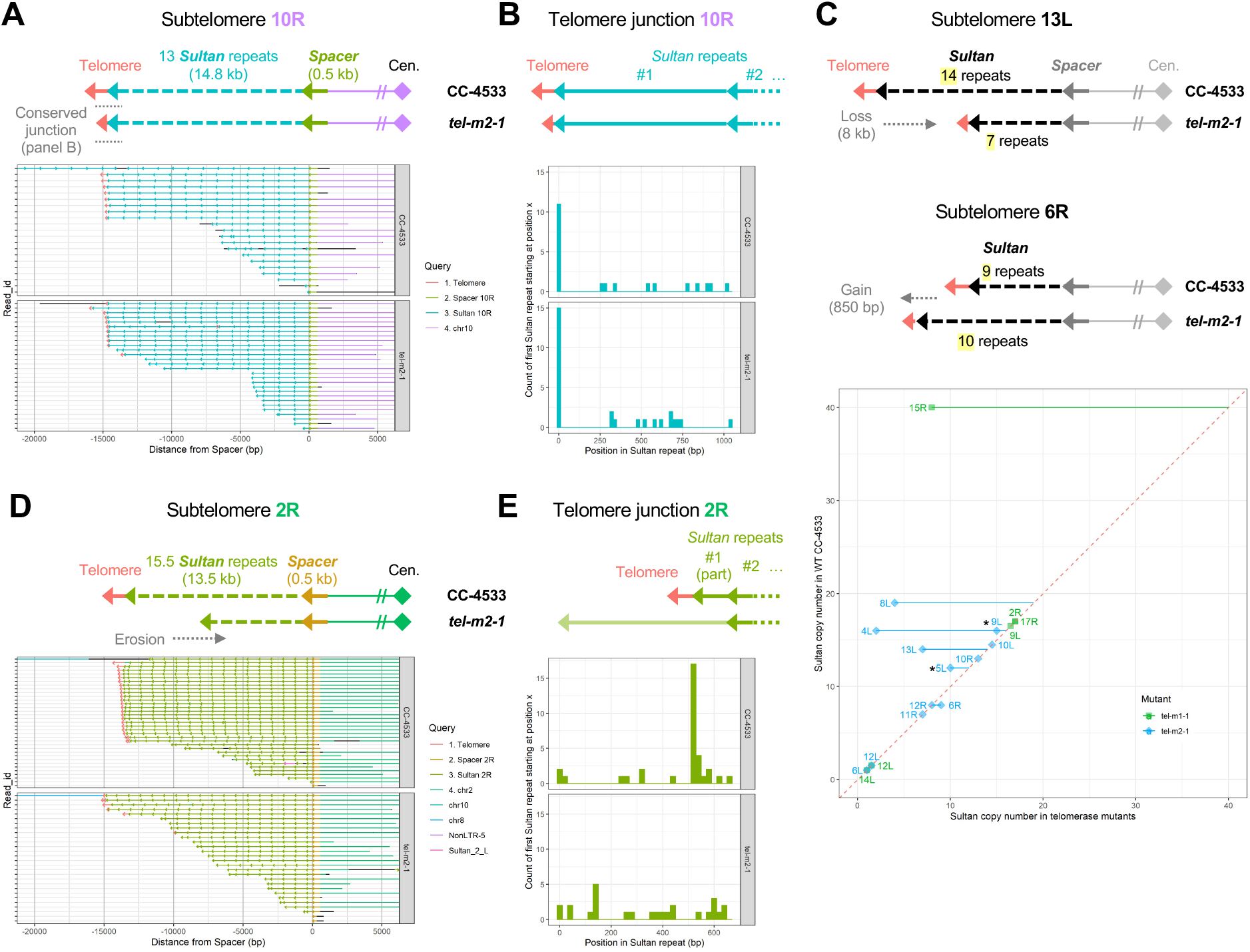
Contraction, erosion and expansion of Sultan arrays. (A) The stabilized 10R Sultan array of *tel-m2-1* capped by a telomere sequence is depicted in comparison to CC-4533. Raw Nanopore reads, colored based on the element (as indicated in the legend; black segments correspond to sequences that did not align to any of the selected queries) and anchored at the Spacer, are shown. (B) For each read, the starting position of the Sultan element adjoining the telomere sequence is recorded and the overall distribution of counts over the length of a Sultan element is represented. (C) Schematic representation of contraction and expansion of stabilized Sultan arrays (upper part) and overall changes in Sultan number in stabilized Sultan arrays for *tel-m1-1* and *tel-m2-1* (lower part). (D) Same as (A) with the unstable 2R Sultan array of *tel-m2-1* mostly uncapped by telomeres. (E) Same as (B) but for the unstable 2R Sultan array of *tel-m2-1*.

**Table 1.** Summary of rearrangements classified by types, found at Spacer-containing subtelomeres. The number of the described rearrangements/events found at Spacer-containing subtelomeres, compared to CC-4533, is indicated. For subtelomeres with reads that represented multiple subpopulations (“Heterogeneous subpopulations”), the rearrangements/events of each subpopulation were counted. Only rearrangements supported by at least two reads were counted.

|  | *tel-m1-0* | *tel-m1-1* | *tel-m2-1* |
| --- | --- | --- | --- |
| Similar to WT | 9 | 7 | 9 |
| Absence of Spacer | 1 | 1 | 3 |
| Contraction of stabilized Sultan array | 3 | 1 | 5 |
| Unstable Sultan array | 3 | 3 | 4 |
| Expansion of stabilized Sultan array | 1 | 0 | 1 |
| Fusion between Sultan arrays (total): | 13 | 17 | 3 |
| - Same orientation | 3 | 5 | 1 |
| - Inverted orientation | 10 | 12 | 2 |
| Recruitment of rDNA | 4 | 5 | 1 |
| Recruitment of other sequences | 3 | 4 | 2 |
| BFB | 4 | 5 | 0 |
| Chromosome circularization | 1 | 1 | 0 |
| Heterogeneous subpopulations | 7 | 9 | 4 |

We first focused on subtelomeres composed, in CC-4533, of an array of Sultan elements as the only repeated element, which are the vast majority^24^ (class A and B, 27 subtelomeres out of 34). In CC-4533, at each class A/B Sultan subtelomere, the reads were consistent with a single population of subtelomeres with a fixed number of Sultans and capped by telomeres, as supported by a majority of reads (Fig. 3A and D). Reads that did not reach the extremity of the chromosome were due to DNA molecules physically broken during the extraction and preparation procedure and would be expected to terminate at random position in the Sultan array and within a Sultan element, which we confirmed (Fig. 3B and E). Additionally, in CC-4533, all reads containing a specific Sultan element and telomere sequence could be retrieved and their number was often close to the number of Spacer-containing reads (Supp. Fig. S3A), consistent the fact that Spacer-containing reads did not always reach a telomere sequence because of the physical breakage of the extracted DNA molecule. Interestingly, both telomerase mutants frequently showed reads that also supported a fixed number of Sultans (Fig. 3A and C). We call such subtelomeres “stabilized Sultan arrays”. However, the number of Sultan elements could differ from CC-4533 (Fig. 3C): in *tel-m1-1* and *tel-m2-1* combined, we observed a decreased number of Sultans in 6 cases by 2-32 repeats, and in 1 case on the contrary, the number of Sultan element was slightly increased by 1. For the rest of the cases (n = 11), no variation of the number of Sultans was observed. We found that stabilized Sultan arrays in telomerase mutants were capped by short telomere sequences (Fig. 3A and Supp. Fig. S3A), suggesting that these telomeres were maintained in a telomerase-independent manner.

We then asked whether at the extremities of stabilized Sultan arrays, the telomeres could be established *de novo* after complete loss or whether Sultan elements were excised or amplified internally independently of the initially present telomere. To address this point, we identified the junction between the telomere sequence and the last Sultan element and compared it between CC-4533 and the mutants (Fig. 3B and Supp. Fig. S3B). In 4 out of 18 stabilized Sultan arrays in the two mutants combined, we found that telomeres transitioned into the Sultan element at a different position within the Sultan compared to CC-4533, indicating that the telomeres were added *de novo* on truncated Sultans (Supp. Fig. S3A and B). In the rest (14 out of 18), telomeres were found at the same position of the last Sultan in CC-4533 and the mutants, suggesting that changes in the number of Sultan elements of the array might also be due to excision of internal Sultan elements, without affecting the last Sultan.

At some subtelomeres (3 for *tel-m1-1* and 6 for *tel-m2-1*), the reads were not consistent with a stabilized number of Sultan elements, and the vast majority had different counts, suggesting a heterogeneous population of cells in which that specific extremity shortened progressively. These Sultan arrays were qualified as “unstable”. This situation is exemplified by subtelomere 2R in *tel-m2-1* (Fig. 3D). Consistent with a progressive erosion of the extremity, the last Sultan element ended at different positions of the Sultan element in different reads and was often not capped by telomeres (Fig. 3E and Supp. Fig. S3A and B). To rule out that these reads containing a variable number of Sultan elements were not simple due to physically broken DNA molecules, we searched for all reads containing the same Sultan elements and telomeres and compared their number with the number of reads containing the corresponding Spacer (Supp. Fig. S3A). In contrast to CC-4533, unstable Sultan arrays in the mutants showed fewer telomere-containing reads than Spacer-containing ones, indicating telomere loss and supporting the idea that these extremities truly ended with Sultan elements. The progressive loss of telomere repeats and Sultan element was consistent with the overall decrease in the number of telomeres and Sultan copy number (Fig. 2B and E). Erosion of Sultan arrays would eventually lead to their complete loss, a situation that we could clearly evidence for subtelomere 7L (Supp. Fig. S3C): in *tel-m1-0*, the subtelomere contained only 3 Sultans, compared to 8 in CC-4533, but in *tel-m1-1* the extremity was eroded beyond the Spacer and a new telomere was formed with the additional translocation, in a subpopulation of reads, of a sequence from chromosome 11 which had a microhomology to the telomere sequence (Supp. Fig. S3D and E). Interestingly, an analogous comparison between CC-4533, *tel-m1-0* and *tel-m1-1* for the class C subtelomere 15L tended to suggest that Suber arrays could also be progressively eroded until completely lost (Supp. Fig. S3F).

### Complex rearrangements within and between subtelomeres

In telomerase mutants, we observed a striking fraction of telomere-containing reads with unusual combinations of subtelomeric elements (Table 1 and Supp. Dataset 1). To better characterize the structure of these rearranged subtelomeres, we again used the Spacer-based analysis to help anchor the reads to a given chromosome extremity.

#### Fusion between subtelomeric elements

Fusions of elements from different subtelomeres were the most frequent genome rearrangement occurring at subtelomeres (Table 1). The simplest type corresponded to the juxtaposition of two or more arrays of different Sultan elements, either in the same or in reverse orientations, such as subtelomere 6R in *tel-m1-0* (Fig. 4A) or subtelomere 15R in *tel-m2-1* (Supp. Fig. S4A). Multiple subtelomeric elements of different types, such as rDNA and Sultan in subtelomere 6L of *tel-m1-0* and *tel-m1-1*, were also observed (Fig. 4B and Table 1). Because rDNA and Sultan do not share sequence homology, we speculate their fusion stemmed from NHEJ-dependent translocation events or end-to-end fusion.

**Figure 4.**
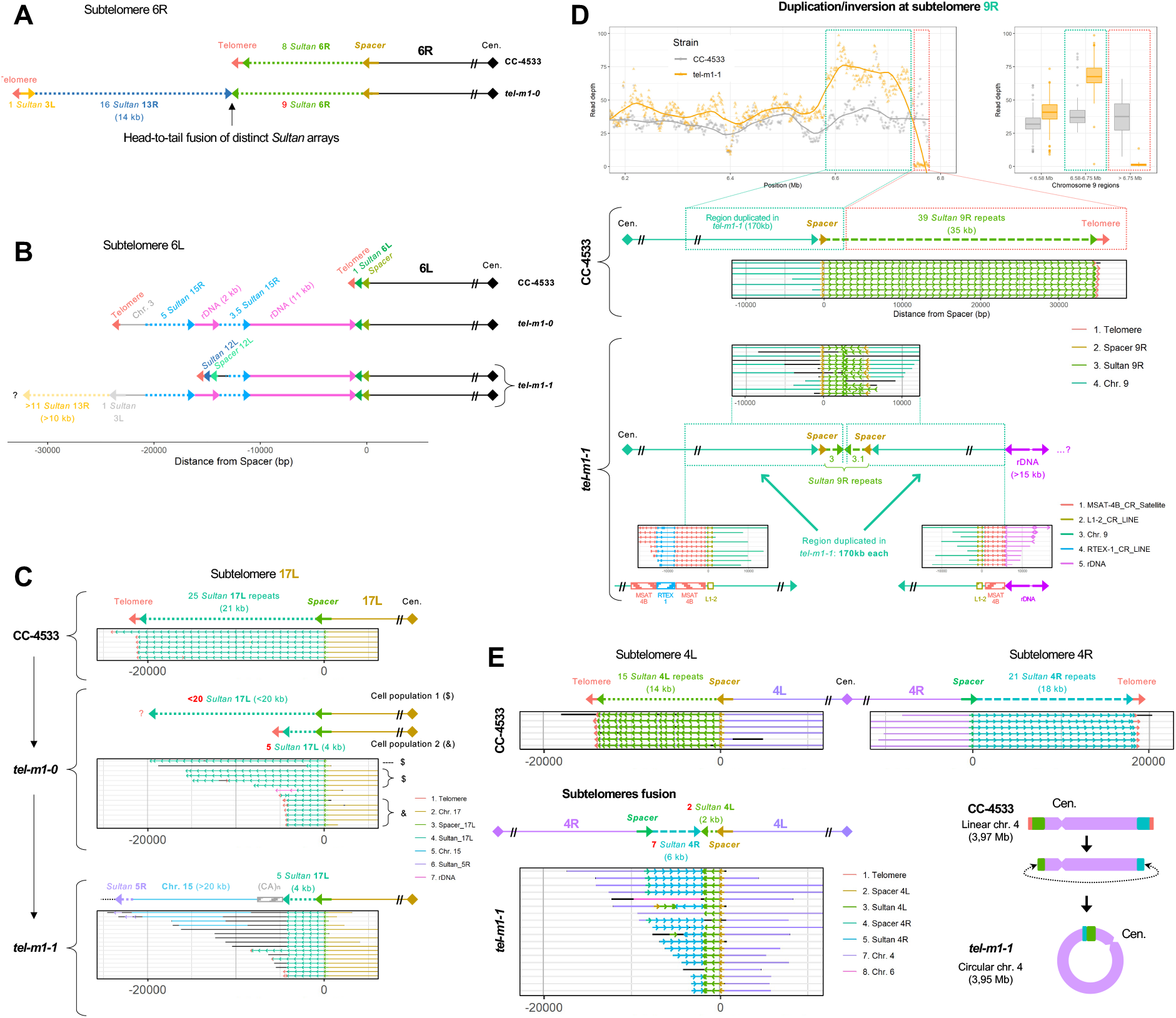
Complex genome rearrangements at subtelomeres. (A) Schematic representation of subtelomere 6R where, in *tel-m1-0*, 2 additional Sultan arrays were fused to the initial Sultan array in inverted orientation. (B) Schematic representation of complex rearrangements at subtelomere 6L in *tel-m1-0* and *tel-m1-1* involving Sultan elements from different subtelomeres and rDNA sequences. In *tel-m1-1*, 2 subpopulations of reads reveal 2 distinct structures stemming from the initial rearrangement found in *tel-m1-0*. (C) Schematic representation of the dynamics of rearrangements at subtelomere 17L between CC-4533, *tel-m1-0* and *tel-m1-1*. The structure of the subtelomere in *tel-m1-1* appeared to have evolved from the structure in cell population 2 of *tel-m1-0*. (D) Signature of a BFB event at subtelomere 9R in *tel-m1-1*. (*Upper part*) Duplication of the last 170 kb of the chromosome end. (*Middle part*) Individual reads supporting the loss of telomeres and 36 Sultan elements, and the end-to-end fusion of the 9R sister chromatids. (*Lower part*) Reads showing the recruitment of > 15 kb of rDNA sequences at the new 9R extremity, disrupting an array of MSTA-4B satellite sequences. Black segments correspond to sequences that are not aligned to any of the selected queries. (E) Representation of reads supporting the fusion of subtelomeres 4L and 4R in *tel-m1-1*, after complete loss of telomeres and a total of 27 Sultan elements. (Lower right) Scheme of the inferred circularization of chromosome 4, also supported by *de novo* genome assembly (Supp. Fig. S4D-E). Black segments correspond to sequences that are not aligned to any of the selected queries.

The complexity of the observed genome rearrangements suggested that they formed through a multistep process. To test this hypothesis, we compared the rearrangements found in *tel-m1-0* to those in *tel-m1-1*, as these two samples were collected with a ∼450-generation delay. We could find intermediate rearrangement states in the earlier *tel-m1-0* sample, suggesting that at least some complex rearrangements were formed in multiple steps over time (Fig. 4B and C; Supp. Fig. S3C and S4B). For example, at subtelomere 6L, the already complex structure found in *tel-m1-0* further changed in *tel-m1-1* and diverged into two structurally distinct subpopulations (Fig. 4B). As another example, subtelomere 17L displayed a reduced number of Sultan elements in *tel-m1-0* compared to wild-type with a subpopulation of reads capped by telomeres, representing a short stabilized Sultan array, but contained additional sequences from a (CA)n microsatellite, from chromosome 15 and from 5R Sultan elements, fused to the remaining 17L Sultans in *tel-m1-1* (Fig. 4C).

#### Signature of BFB events

The read coverage mapped to the assembly of *tel-m1-1* genome identified a duplicated region at the extremity of chromosome 9R (Fig. 4D, upper part). Closer inspection of the reads anchored by their Spacer element revealed a pattern of 2 inverted Sultan arrays fused to each other in a head-to-head manner (Fig. 4D, middle part). All Sultan elements could be attributed to subtelomere 9R unambiguously. Each array was adjacent to its own 9R Spacer element as well as downstream 9R-specific sequences. Overall, this configuration was indicative of a fusion between sister chromatids followed by a breakage in anaphase, an event that initiated a canonical breakage-fusion-bridge cycle until the extremities were stabilized. We then looked for the new 9R extremity in *tel-m1-1* and identified the reads that mapped close to the 1X/2X coverage boundary. Strikingly, reads that mapped only to the 2X side continued with at least two rDNA units over >15 kb (Fig. 4D, lower part). The rDNA sequence disrupted a RTEX-1 long interspersed nuclear element (LINE) retrotransposon framed by two arrays of MSAT-4B satellites, as shown in Fig. 4D. What lay beyond the rDNA array remained unknown, in particular whether the extremity was capped by additional telomere sequences. In another example, the structure of the 11R extremity of *tel-m1-0* was consistent with at least two cycles of BFB, based on the multiple inverted repeats (Supp. Fig. S4C). We found a total of 5 events consistent with BFB in *tel-m1-1* (Table 1), indicating that BFB cycles were a common mechanism at play once telomeres became too short or were lost.

#### Circularization of chromosome 4

We report the head-to-head fusion of subtelomeres 4L and 4R in *tel-m1-1*, after complete loss of the telomere sequences and partial loss of Sultan elements (from 15 to 2 for 4L and from 21 to 7 for 4R) (Fig. 4E). The *de novo* assembly of chromosome 4 in *tel-m1-1* without relying on scaffolding with a reference sequence indicated that chromosome 4 was made of a single contig with no extremity and had therefore a circular structure (Supp. Fig. S4D-E). Circularization of the chromosome bypasses the need for functional telomeres to stabilize the extremities of linear chromosomes.

### New telomeres outside canonical subtelomere regions lead to drastic karyotype alterations

We next investigated terminal-telomere-containing reads not associated with the canonical subtelomeric elements, with the aim of revealing the formation of new extremities outside their normal subtelomeric contexts. Using TeloReader, we gathered all reads containing telomere sequences that were not associated to known subtelomeric elements (*Sultan, Spacer, Suber, Subtile* or rDNA). We mapped these reads on the genome of CC-4533 and found such subtelomere-less telomeres at 3 and 2 locations in *tel-m1-1* and *tel-m2-1*, respectively. Consistently with the genome assembly, we also detected the new telomere extremity of the duplicated mini-chromosome 1 from CC-4533, which transitioned directly into the chromosome 1 sequence through a microhomology and without subtelomeric element nor other exogenous sequence (Supp. Fig. S5A).

At subtelomeres 14L in *tel-m2-1* and 7L in *tel-m1-1* (Fig. 5A, Supp. Fig. S3C-E), telomeres connected to the chromosome arm several kilobases away from the expected telomere location in CC-4533, and the corresponding arrays of *Sultan* repeats were lost, but without loss of annotated genes. At these novel telomere junctions, the chromosome arm showed no sequence homology with telomeres, suggesting that the telomere was recruited at these sites by non-homologous end joining rather than homology-mediated mechanisms. At subtelomere 7L in *tel-m1-1*, a minor population of reads showed an additional fragment from chromosome 11 next to the new telomere, with a 4-bp microhomology at the junction (Supp. Fig. S3D). The resulting interstitial telomeric sequence was shorter than the telomeres observed in the main population of reads (Supp. Fig. S3E).

**Figure 5.**
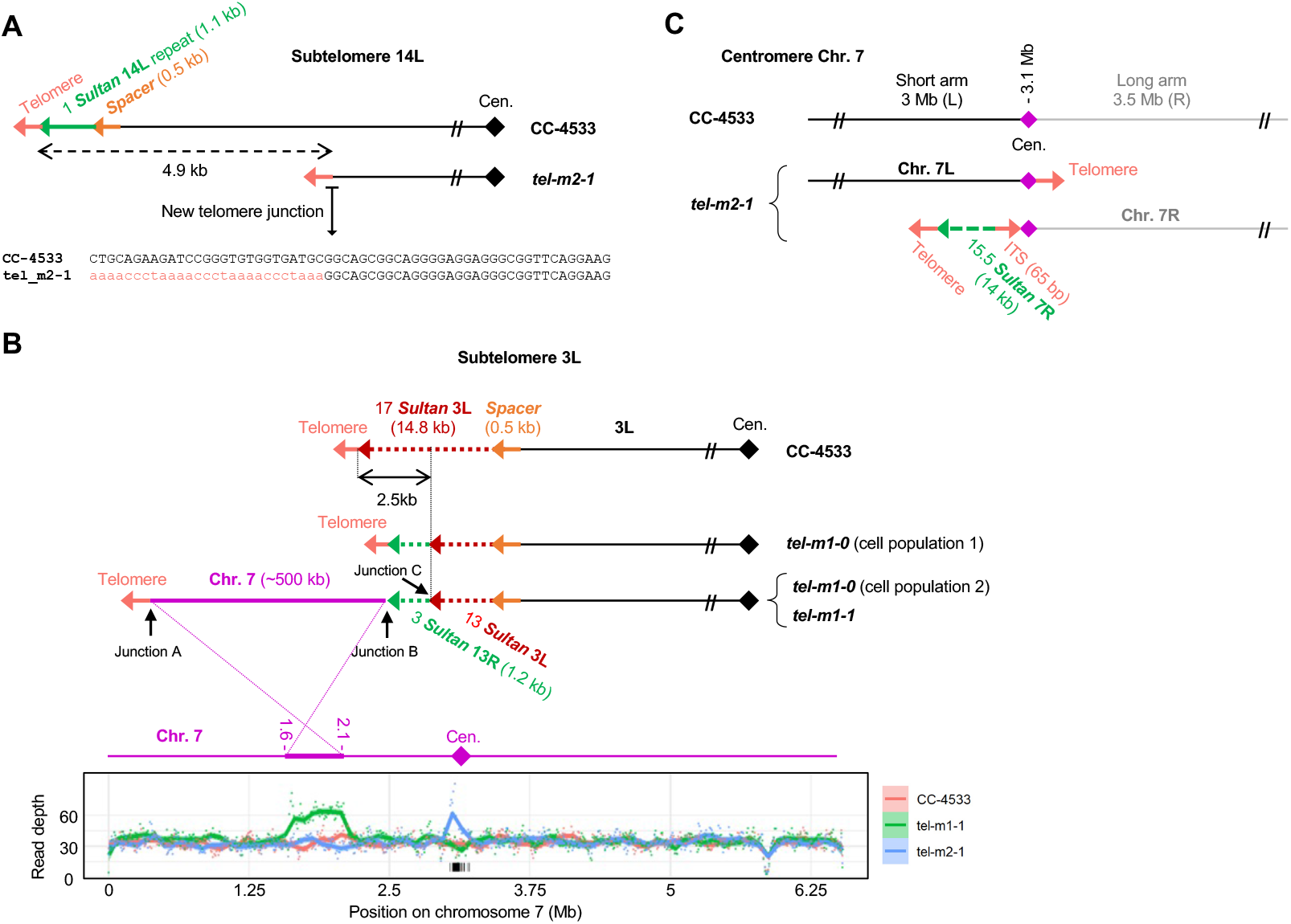
New extremities without canonical subtelomeric elements. (A) Scheme of the loss of 4.9 kb of sequence and formation of a new telomere at subtelomere 14L of *tel-m2-1*. The new junction sequence is shown. (B) Scheme of the complex new 3L extremity in *tel-m1-0* and *tel-m1-1*. The sequencing depth for chromosome 7 is shown, revealing the duplicated 500-kb region in *tel-m1-1*. The sequences of Junction A, B and C are shown in Supp. Fig. S5B-D. (C) Scheme of the Robertsonian fission that occurred on chromosome 7 of *tel-m2-1*. The new centromere-proximal extremities were stabilized by telomeres, with in addition a Sultan array and an ITS for the 7R long arm.

The other new telomeres were involved in more complex rearrangements. The subtelomere 3L of *tel-m1-0* had 13 Sultan elements remaining after loss of the telomere sequence and 4 Sultan elements, followed by a short array of 13R Sultan elements and telomere sequence (Fig. 5B). In a subpopulation of *tel-m1-0* reads and in *tel-m1-1*, this structure was further altered by the loss of the telomere and the fusion of 500 kb of duplicated sequence originally located at 2.1 Mb from the end of the left arm of chromosome 7, and capped by a new telomere (Fig. 5B). Junctions to telomeres (junction A) and to the 13R Sultan array (junction B) involved micro-homologies of 2-4 bp (Supp. Fig. S5B and C), while the Sultans from 13R and 3L were connected with the insertion of a short sequence of unknown origin (junction C) (Supp. Fig. S5D), but without homology nor truncation of the neighboring Sultan elements.

On chromosome 12 of *tel-m1-1*, the 2-Mb distal part of the right arm was translocated to the subtelomere 5L and the new right end of chromosome 12 (at 7.7 Mb from its initial extremity) carried telomeres preceded by a tandem duplication consisting of a fragment from chromosome 12, 9 telomeric repeats forming an interstitial array and a short Sultan array from subtelomere 7L (Supp. Fig. S5E).

In *tel-m2-1*, new telomeres were found at two close locations around the centromere of chromosome 7, at the extremities of a 90-kb region displaying a 2X sequencing depth, indicating that the centromere was duplicated and the two arms of chromosome 7 split into two telocentric chromosomes (Fig. 5C), thus constituting a Robertsonian fission, the reciprocal event of a Robertsonian fusion whereby two telocentric chromosomes are combined through their long arm with the loss of the very short arms and one centromere^55,56^. On the left end of the right arm, the new telomere capped an array of Sultan 7R lacking Spacer and a 65-bp ITS. On the right end of the left arm, the new telomere directly transitioned into the centromere.

Overall, duplications, non-reciprocal translocations, deletions and more complex rearrangements created new extremities outside canonical regions and altered the karyotype of the telomerase mutant strains. For each new extremity, stability seemed to be ensured by newly recruited telomere sequences.

### DNA methylation is maintained at chromosome extremities and at displaced Sultan arrays

We previously showed that Sultan arrays and to a lesser degree, Suber arrays, were hypermethylated, whereas the two major rDNA clusters were hypomethylated except for a few telomere-proximal repeats^24^. Sultan subtelomeres are also associated with the heterochromatin mark H3K9me1^57^. Since subtelomeres in telomerase mutants were heavily rearranged, we asked whether DNA methylation remained associated with the Sultan elements that were no longer close to chromosome extremities and whether new extremities would acquire hypermethylation.

We thus basecalled 5-methylcytosines (5mC) at CpG sites using Nanopolish^58^ and focused on rearranged loci. We first investigated the complex rearrangement found at extremity 3L of *tel-m1-1* where a duplicated 500 kb of chromosome 7 capped with telomere formed the new extremity and the original 3L Sultan elements were thus located much more internally (> 500 kb) in the chromosome (Fig. 5B). To analyze 5mC content of the new extremity, we selected reads that aligned to the 500 kb of chromosome 7 and at the same time contained the terminal telomere sequence, and in a second control group, reads that aligned to the 500 kb region but also continued beyond on chromosome 7 (Fig. 6A and Supp. Fig. S6A). Strikingly, the new 3L extremity was highly methylated over a length of ∼12 kb, whereas the same sequence on its original locus on chromosome 7 was unmethylated, and so was the same locus on the wild-type strain (Supp. Fig. S6A). The 3L Sultan elements as well as the neighboring 13R Sultans in *tel-m1-1* maintained their high methylation status despite being now > 500 kb away from the extremity.

**Figure 6.**
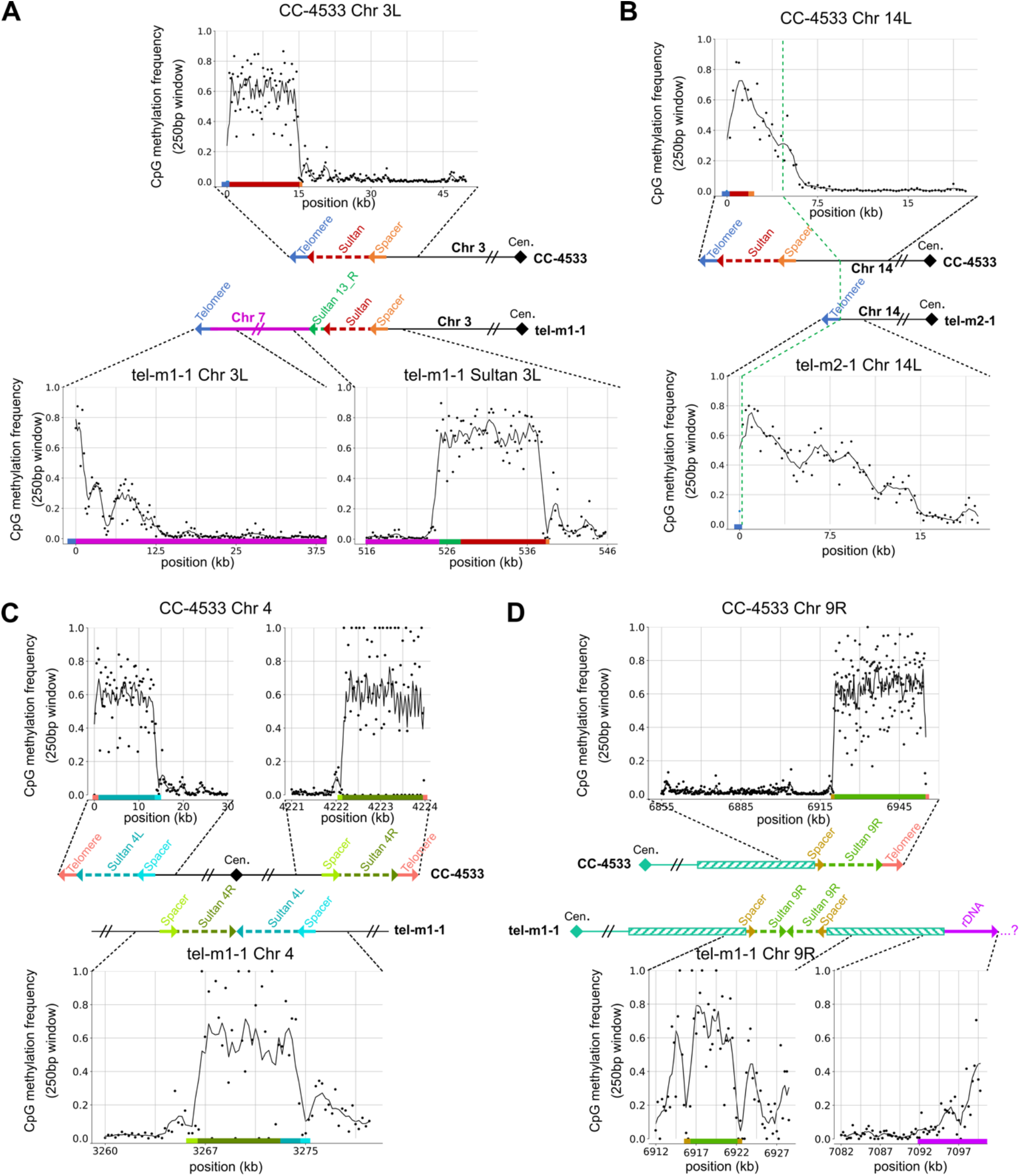
Methylation frequency of new extremities and displaced Sultan elements. (A-D) Analysis of 5mC frequency at CpG sites using reads that unambiguously spanned the indicated regions, in different illustrative cases. (A) At the rearranged 3L chromosome arm of *tel-m1-1*. Methylation frequency was plotted over different regions of subtelomere 3L in *tel-m1-1* and in CC-4533 as depicted. See also Supp. Fig. S6A for the methylation frequency at chromosome 7 in CC-4533 and *tel-m1-1*. (B) At the broken 14L subtelomere of *tel-m2-1*, where internal genomic sequence was then capped by a telomere, compared to CC-4533. (C) At the fused 4L and 4R Sultan arrays of *tel-m1-1*, compared to the two arrays in CC-4533. (D) At the head-to-head fused 9R Sultan arrays and the transition to rDNA in *tel-m1-1*, compared to the 9R extremity of CC-4533.

That new extremities were hypermethylated even though they did not contain canonical subtelomeric sequences was confirmed at the truncated 14L extremity of *tel-m2-1* (Fig. 5A and 6B) and 7L of *tel-m1-1* (Supp. Fig. S6B). Other examples of Sultan elements no longer located at subtelomeres but still methylated included the 4L and 4R Sultan elements of the circularized chromosome 4 in *tel-m1-1* (Fig. 4E and 6C) and the head-to-head fused 9R Sultan elements, which resulted from the BFB event we described in a previous section (Fig. 4D and 6D). In the latter example, the Sultan elements were located at least 185 kb away from the new extremity. The reads mapping to the rDNA sequence at the 9R subtelomere showed that the rDNA was methylated (Fig. 6D), which in CC-1690 was only true for the rDNA of subtelomere 1L and a few telomere-proximal rDNA sequences at 8R and 14R subtelomeres, suggesting that the rDNA might constitute the new subtelomere of 9R in *tel-m1-1*. Subtelomere 16R in *tel-m2-1* provided another example of translocated rDNA, now directly between a terminal telomere sequence and Sultan elements, which was also hypermethylated (Supp. Fig. S6C).

Overall, the Sultan elements that were no longer near chromosome extremities maintained their hypermethylation level. This observation was verified in all cases in *tel-m1-1* and *tel-m2-1*, for Sultan arrays separated from the extremity by up to > 500 kb of other non-methylated sequences. Conversely, sequences that became capped by telomeres and formed new extremities acquired a hypermethylation pattern.

## Discussion

Telomeres and subtelomeres are extensively implicated in genome instability induced by telomere shortening or dysfunction, as exemplified in post-telomere-crisis tumors^13^. Many types of telomere-related rearrangements have been studied in experimental systems of different model organisms and in tumor samples from patients. However, a direct assessment and comprehensive picture of the repertoire of genome rearrangements that functional telomeres protect from have rarely been achieved. Here, we took advantage of long-read Nanopore sequencing to access telomere-induced SVs that would have escaped detection from short-read sequencing or at least would have been difficult to identify, particularly in repeated regions. We chose to sequence heterogeneous populations of telomerase-negative cells containing a complex mixture of subclonal genome rearrangements and to perform our analyses mainly at the level of reads, revealing mosaic rearrangement patterns that would have been missed in assembled clonal genomes.

### Alternative telomere maintenance

We previously demonstrated that telomeres in *C. reinhardtii* are maintained by telomerase and that mutants of *CrTERT*, the gene encoding its catalytic subunit, displayed an “ever shorter telomere” phenotype^45^. We evidenced cell death consistent with replicative senescence in some telomerase mutant cultures but also that other long-term cultures failed to show any growth defect, suggesting that cells with alternative telomere maintenance mechanisms were selected. In this work, analysis of chromosome extremities in the long-term cultures of telomerase mutants showed that overall ∼60% were still capped by short telomeres, reminiscent of type I post-senescent survivors in *S. cerevisiae* ^37,59^. Interestingly, for at least a subset of them, the telomere sequence was likely recruited *de novo* on a truncated subtelomere by non-reciprocal translocation, as shown before for cancer cell lines^60^. In all cases, telomeres were maintained, albeit at a short length distribution, which implied that an alternative maintenance mechanism was used, relying for example on homologous recombination or BIR.

A number of chromosome extremities completely lacked telomere sequences and their lost telomeres might have been translocated to the new telomere sites we described. Individual read analysis of such telomere-less extremities showed that the Sultan array composing the subtelomere displayed a length distribution at the population level consistent with progressive loss of sequence. This observation leads to the intriguing possibility that the array of Sultan elements might directly function in end protection instead of telomeres. Consistently, in end-to-end fusion events involving Sultan arrays (e.g. Fig. 4D and E), fewer than 10 Sultan elements were left when fusion occurred, which might suggest that larger Sultan arrays could confer some partial telomere protection. The Sultan element must then ensure telomere protection by for example binding a yet unknown factor, which might be achieved through the short telomere-like sequence at the beginning of the element^24^ or through another intrinsic binding site. Since Sultan elements are associated with heterochromatin, indirect binding of factors on modified histones might also provide sufficient protection, as proposed for some telomerase-negative *S. pombe* survivors^61^. Consistently, even when the chromosome extremities are not composed of Sultan elements, but of rDNA repeats or even other sequences, the DNA sequence is highly methylated (see discussion below).

Alternatively, *C. reinhardtii* might transiently tolerate unprotected or partially unprotected telomeres, as a result of the lack of detectable DNA damage checkpoint kinases Chk1 and Chk2 orthologs or at least differences in DNA damage response machineries compared to other Viridiplantae and to Opisthokonts^62,63^.

Another strategy to maintain chromosome integrity without telomerase consists in chromosome circularization, which we observed for chromosome 4 in *tel-m1-1*. Although we encountered only one such circularization event, the fact that we did not detect another subpopulation with another structure for chromosome 4 suggests that this circular chromosome conferred some selective advantage (e.g. telomerase-independent genome integrity) or at least was not counter-selected. A similar circularization strategy in response to telomerase absence or disruption of telomere protection was also described in other eukaryotes^38,64-67^ and therefore appears to be relatively well-tolerated across evolution.

Overall, although they came at the cost of wide-spread genome rearrangements, the diversity of alternative strategies to maintain chromosome integrity in telomerase mutants found in a single organism is remarkable and illustrates genome plasticity at short timescales.

### Mechanisms of telomere-shortening-induced genome rearrangements

Because the approach used in this work consists in a direct visualization of chromosome sequences on a large scale with as little source of bias as possible, we were able to document a striking variety of rearrangements. The fact that most rearrangements involved telomeres and subtelomeres suggested that critically short or dysfunctional telomeres preferentially induced local instabilities, which can propagate through various mechanisms such as BFB cycles involving dicentrics, as reported in yeast and human tumors^4,5,13,68-70^.

Short or dysfunctional telomeres can lead to end-to-end fusions as observed across evolution^14,71-75^, although telomere sequences themselves may be completely absent at fusion sites in telomerase-negative cells^3,4,38,64^. Despite many instances of end-to-end fusions, we found no remaining telomere sequences at fusion sites in our read datasets, suggesting that in *C. reinhardtii*, even very short telomeres are sufficient to inhibit fusions. Instead, we detected many subtelomere-subtelomere fusions, the most frequent involving Sultan arrays. They can occur between different subtelomeres, as in the case of chromosome 4 circularization, or between sister chromatid subtelomeres. Sister chromatid fusions were followed by cycles of BFB, propagating genome instability over multiple divisions and over more internal genomic regions, as for extremity 9R in *tel-m1-1*. Of note, we did not find any signature of chromothripsis, resulting from the aberrant repair of the fragmentation of a chromosomal region, despite chromothripsis being associated with BFB cycles and telomere crisis in cancer^76,77^.

Interestingly, most rearrangements implicated the repeated elements that are found at the subtelomeres, which could promote homology-mediated mechanisms for amplification, deletion and translocation. When different types of elements were found juxtaposed, the transition sequence often involved microhomologies of a few base pairs (e.g. Fig. 5B and Supp. Fig. S5B-C), suggesting that mechanisms such as microhomology-mediated break-induced replication (MMBIR)^78,79^ or microhomology-mediated end-joining (MMEJ)^80^ might be involved. Even when no homology or microhomology was detected at junctions, canonical subtelomeric elements were still frequently found in rearrangements. We speculate that these subtelomeric elements were frequently excised from their native locus due to terminal instability and were therefore available in subsequent fusion events. Nevertheless, other internal genomic regions were sometimes duplicated and translocated to chromosome ends, supporting the idea of a genome-wide increase in genome instability in telomerase-negative cells^3,4,6,8,9^.

### New chromosome extremities establish heterochromatin

We found that new telomere-capped chromosome extremities formed by sequences originating from internal genomic regions showed high levels of 5mC. We therefore speculate that telomeres themselves are sufficient to establish new DNA methylation domains in *C. reinhardtii*, likely to be associated with heterochromatin, stabilizing the chromosome end and protecting it from degradation and repair. While the heterochromatic nature of telomeres and subtelomeres is conserved in eukaryotes (except for *A. thaliana* where it is less well established^81-88^), whether and how heterochromatin forms at a new telomere is less well known across organisms. The detection of new extremities in our work thus provides an additional piece of evidence for heterochromatin formation spreading from the telomere sequence being an evolutionarily conserved property of chromosome extremities. Since many chromosome extremities were not capped by telomere sequences but directly by the hypermethylated Sultan arrays, one can speculate that heterochromatin might be the primary functional feature of a protected extremity, even more important than telomere sequences themselves.

Interestingly, Sultan elements that are displaced from their normal subtelomeric location retain a high level of 5mC, even when found at > 500 kb of the extremity of the chromosome. We previously showed that Sultan elements are exclusively found at subtelomeres in wild-type strains. We thus propose that their sequence coevolved with their function at subtelomeres, so that they might recruit methyltransferases to actively contribute to heterochromatin maintenance and spread. This property would allow then to remain hypermethylated even when displaced from subtelomeres. Interestingly, in most cases, the Spacer sequence still acted as the boundary of the hypermethylated domain even when not located at subtelomeres.

To conclude, our results show that *C. reinhardtii* telomerase-negative cells use a variety of strategies to protect chromosome extremities, including the establishment of DNA methylation, and undergo diverse and complex rearrangements, highlighting a remarkable plasticity of the genome. This work demonstrates the potential of long-read sequencing to provide a comprehensive view of complex genome rearrangements even in mosaic populations of cells at the level of subclones, an analysis that could be applied to tumor genome heterogeneity.

## Material and Methods

### Strains and growth conditions

Wild-type CC-4533 and *tel-m1* and *tel-m2* mutant strains were obtained from the Jonikas lab^46^. They were maintained on plates in TAP medium^89^ at 25°C under low light (5 µE.m^-2^.s^-1^) and restreaked in bulk without subcloning. Before DNA extraction, cells were grown in 200 mL liquid TAP medium until they reached ∼2×10^7^ cell/mL and collected by centrifugation at 5000 g for 5 min.

### DNA extraction and size selection

To extract genomic DNA while preserving high-molecular-weight fragments, we developed a modified version of a Joint Genome Institute protocol (https://www.pacb.com/wp-content/uploads/2015/09/DNA-extraction-chlamy-CTAB-JGI.pdf), as described here^47^. The extraction is followed by a clean-up using magnetic beads and size selection based on solid-phase reversible immobilization^90^ using the SRE kit (Circulomics).

### Library preparation and sequencing

Sequencing libraries were prepared from SRE-treated high-molecular-weight DNA (except for *tel-m1-1* which was sequenced in two runs, one without SRE treatment and one with), following Oxford Nanopore Technologies (ONT) protocols for genomic DNA without pre-amplification. Kits LSK109 and LSK110 were obtained from ONT and Companion Module NebNext from New England Biolabs (NEB). As a minor modification, we started the preparation with 3 µg of DNA. The second *tel-m1-1* library was barcoded using barcode NB07 from kit NBD104 (ONT).

DNA libraries were sequenced on R9.4 or R10.4 Nanopore flow cells in a MinION Mk1C sequencer with default parameters on the MinKNOW operating software, except for the basecalling with Guppy which was set to real-time high accuracy and for the MUX scan which was decreased to 1 hr. For each run, 500 ng of DNA were loaded and sequenced for 8-12 hrs. The flow cell was then washed using the Wash kit (ONT) and reloaded with the same sample, up to 4 times.

### Genome assembly

Reads in FASTQ files and with quality >7 were processed using Porechop with default parameters (https://github.com/rrwick/Porechop), for removing adapters/barcodes and splitting artefactual chimeras. Reads were assembled using Canu^91^, SMARTdenovo^92^ and NextDenovo (https://github.com/Nextomics/NextDenovo) using default parameters, and all chromosomes were successfully represented by one main contig. These draft assemblies were polished using Racon^93^ (https://github.com/lbcb-sci/racon) and Medaka (ONT; https://github.com/nanoporetech/medaka) with the same reads, then scaffolded on the CC-1690 reference genome using RagTag^94^ (https://github.com/malonge/RagTag). These assemblies were compared to each other and to references (CC-1690, CC-4532) using D-genies^95^ (https://github.com/genotoul-bioinfo/dgenies) to assess contiguity. To generate a genome model for each strain, we chose for each chromosome the assembly giving the most colinear chromosome with CC-1690.

To test if chromosome 4 of *tel-m1-1* was circular at the assembly level without relying on a reference genome for scaffolding, we used the assembler Flye^96^ (https://github.com/fenderglass/Flye), as it generated contigs long enough for this purpose. Visualization of the circular chromosome 4 was done using Bandage^97^ (https://rrwick.github.io/Bandage/).

### Telomere sequence detection

To detect telomere sequences in individual Nanopore reads, we developed TeloReader, a Python (v3.9.12) script with only 3 dependencies (pandas (v1.4.3), numpy (v1.19.5) and matplotlib (v3.5.1)). TeloReader scans each read in both directions, from 5’ to 3’ and from 3’ to 5’, to search for the C-rich telomere motif and the G-rich one, respectively. In the first step of TeloReader, the DNA sequence is transformed into a series of scores corresponding to the best alignment score (using pairwise2 from the package biopython (v1.79)) of each 8-mer against all 8 circular permutations of the telomere motif (CCCTAAAA for the C-rich and TTTTAGGG for the G-rich). This score is between 0 and 8, but all scores lower than 4 are replaced by 4, to reduce the impact of sequencing errors in the following step. The second step defines the sequence in a sliding window of size 15 bp (size_window) as telomeric if the average score is greater or equal to 7 (min_mean_window). In a third step, consecutive overlapping telomeric sliding windows are merged to form a single telomere sequence. We added other constraints and rules to ensure the specificity and sensitivity of TeloReader: the minimal length of a telomere sequence is set at 16 bp (min_len); a telomere sequence must contain at least 8 scores of 8; a configuration where a non-telomeric sequence is found between 2 telomere sequences (as defined after the third step) can be considered as a telomere sequence is the non-telomeric sequence is shorter than 20 bp (max_size_gap) and if the average score of the whole sequence encompassing the 2 telomere sequences and the non-telomeric sequence in-between is greater than 6.5 (min_mean_telo). A telomere sequence is considered as terminal if it is found at < 50 bp from the extremities of the input sequence (3’ extremity for a C-rich telomere and 5’ extremity for a G-rich telomere). The code for TeloReader is available at: https://github.com/Telomere-Genome-Stability/Telomere_2023/tree/main/TELOREADER.

### 5-Methylcytosine detection

To detect 5mC in a CpG context, we used nanopolish^58^ (https://github.com/jts/nanopolish) on the FAST5 files and their associated FASTQ files. Reads were aligned to their corresponding genome assembly using Minimap2^98^ (https://github.com/lh3/minimap2). Alternatively, for detection of 5mC in subpopulations, reads were first selected using fast5_subset (https://github.com/nanoporetech/ont_fast5_api) and aligned to a consensus sequence obtained through a multiple alignment using Clustal W^99^. The frequency of methylation for each base was obtained using nanopolish call-methylation and the script calculate_methylation_frequency.py.

### Bioinformatic analysis

For genome-to-genome comparison, genome models were aligned against references genomes (CC-1690 and CC-4533) using Minimap2^98^ and visualized with Circos plots^48^. SV caller MUM&Co^53^ (https://github.com/SAMtoBAM/MUMandCo) was used for structural variant detection and classification. For read analysis, reads were mapped to genomes using Minimap2 with parameters: -a -x map-ont -K 5M -t 3. We used Integrative Genome Viewer (IGV)^100^ (https://igv.org/) and Tablet^101^ (https://ics.hutton.ac.uk/tablet/) to visualize read mapping. To calculate sequencing depth, the genomes were divided into windows using ‘makewindows’ from BEDTools^102^ (https://bedtools.readthedocs.io/en/latest/index.html), then depth was averaged over the windows using ‘bedcov’ from Samtools^103^ (https://www.htslib.org/) with parameters: -g SUPPLEMENTARY with or without -q 60.

For read level analysis, subtelomeric elements (Spacer, Sultan, rDNA and Suber) were searched in reads using blastn^104^ with parameters: -max_target_seqs 10000 -evalue 0.001 -outfmt “6 qaccver qlen saccver slen pident length mismatch gapopen qstart qend sstart send evalue bitscore”. Multiple hits at close positions were filtered using a custom bash script: hits were sorted by decreasing order of bitscore (*sort* -rgk 14) and overlapping matches were eliminated. Reads containing subtelomeric elements or telomeres (detected by TeloReader) were further blasted against a library of repeated elements^105^ and against reference genome, both of which were then filtered as above to exclude multiple matches. Annotations from all blast hits were then plotted on reads using R. Plots were created using an R script based on dplyr, tidyr and ggplot2 libraries. All lists of filtered blast hits lists were merged with ‘bind_rows’, reads of interest were identified (e.g. match to a given Spacer or Sultan) with ‘filter’ and all hits on this read subset were extracted with ‘semi_join’. To order these reads, first, an anchor was selected, most often a specific Spacer sequence, allowing to set an origin and the orientation of each read. From these two pieces of information, new coordinates were re-calculated for each hit along the reads and for read extremities. Hits were then plotted with distinct reads along the y-axis and coordinates along the x-axis (e.g. Fig. 3A, 3D, 4C-E). Due to the error rate of Nanopore sequencing at the raw read level together with rare artefactual chimeric reads^106^, we only describe rearrangements structurally supported by at least two reads.

Computations were mostly run on the cluster of the French Institute of Bioinformatics (https://www.france-bioinformatique.fr/en/home/).

## Acknowledgement

We thank Olivier Vallon for his critical reading of the manuscript. Research in GF’s laboratory was supported by the French National Research Agency (“ANR” grants: ANR-16-CE 12-0019 and ANR-18-CE12-0004). Research in ZX’s laboratory was supported by ANR grant “AlgaTelo” (ANR-17-CE20-0002-01), by Ville de Paris (Programme Émergence(s)) and by the French National Cancer Institute (grant: INCa_15192). FC is currently supported by a Marie Skłodowska-Curie Actions Postdoctoral Fellowship (101064365 – Cocco-Next).

## Author contributions

Investigation: FC, NA, CG, SE; Conceptualization: FC, GF, SE, ZX; Methodology: FC, NA, CG; Software: FC, CG; Formal analysis: FC, NA, CG, SE; Supervision: GF, ZX; Writing – Original Draft: FC, ZX; Writing – Review & Editing: all authors.

