## Supplementary Data for "Telomerase-independent survival leads to a mosaic of complex subtelomere rearrangements in *Chlamydomonas reinhardtii*"

**Title: Telomerase-independent survival leads to diverse and complex subtelomere rearrangements in *Chlamydomonas reinhardtii***

Frédéric Chaux[1,3], Nicolas Agier[1,*], Clotilde Garrido[1,*], Gilles Fischer[1], Stephan Eberhard[2] and Zhou Xu[1]

**Affiliations:**

[1]Sorbonne Université, CNRS, UMR7238, Institut de Biologie Paris-Seine, Laboratory of Computational and Quantitative Biology, 75005 Paris, France

[2]Sorbonne Université, CNRS, UMR7141, Institut de Biologie Physico-Chimique, Laboratory of Chloroplast Biology and Light-Sensing in Microalgae, 75005 Paris, France

[3]present address: Laboratory for Marine and Atmospheric Biogeochemistry, Ruđer Bošković Institute, 10000 Zagreb, Croatia

*These authors made equal contributions to this work.

| Strain | Read count | Total Gb | Depth | Mean read length (kb) | N50 (kb) |
|---|---|---|---|---|---|
| **CC-4533** | 541 694 | 5.42 | 35X | 10.0 | 19.9 |
| ***tel-m1-0*** | 308 009 | 3.23 | 25X | 10.5 | 24.1 |
| ***tel-m1-1*** | 917 886 | 4.78 | 35X | 5.2 | 14.3 |
| ***tel-m2-1*** | 381 156 | 4.18 | 35X | 11.0 | 19.9 |

**Supp. Table ST1. CC-4533 and telomerase mutants Nanopore sequencing statistics.** Shown are the "passed" datasets (read quality score >7). "N50" is the median read length weighted by base count.

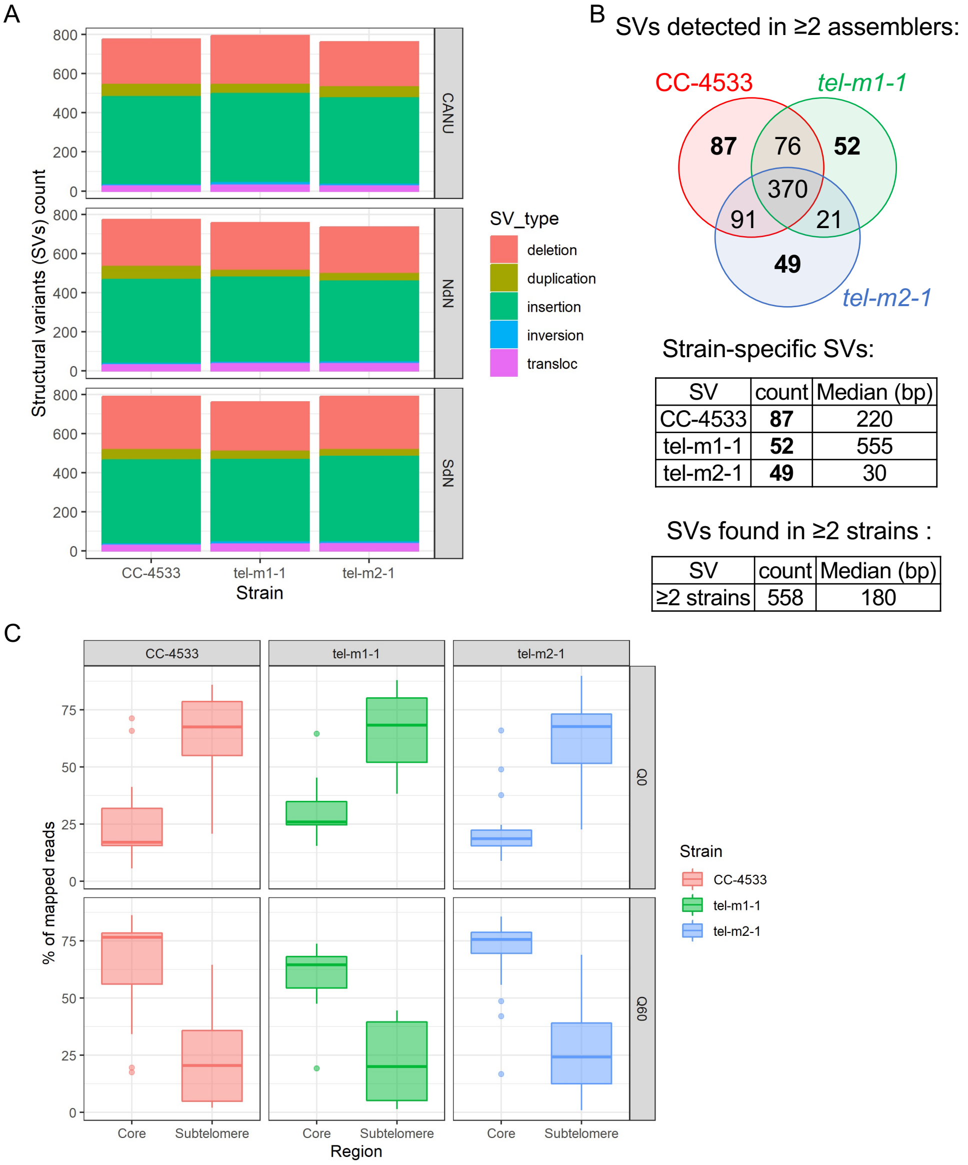


**Supplementary Figure 1. Analysis of SVs in the genome assemblies.** (A) Count of SVs of the indicated types detected by MUM&Co in the assemblies of CC-4533, *tel-m1-1* and *tel-m2-1*, obtained from 3 assemblers (Canu, NextDenovo and SMARTdenovo), and compared to CC-1690. (B) Venn diagram representation of unique and common SVs detected in the 3 strains, for SVs found in the genomes built by at least 2 assemblers. The tables give the median size of the SVs. (C) Boxplots of the percentage of reads of the lowest mapping quality (Q0) and of the highest (Q60), for reads mapping to subtelomeres and the rest of the genome.

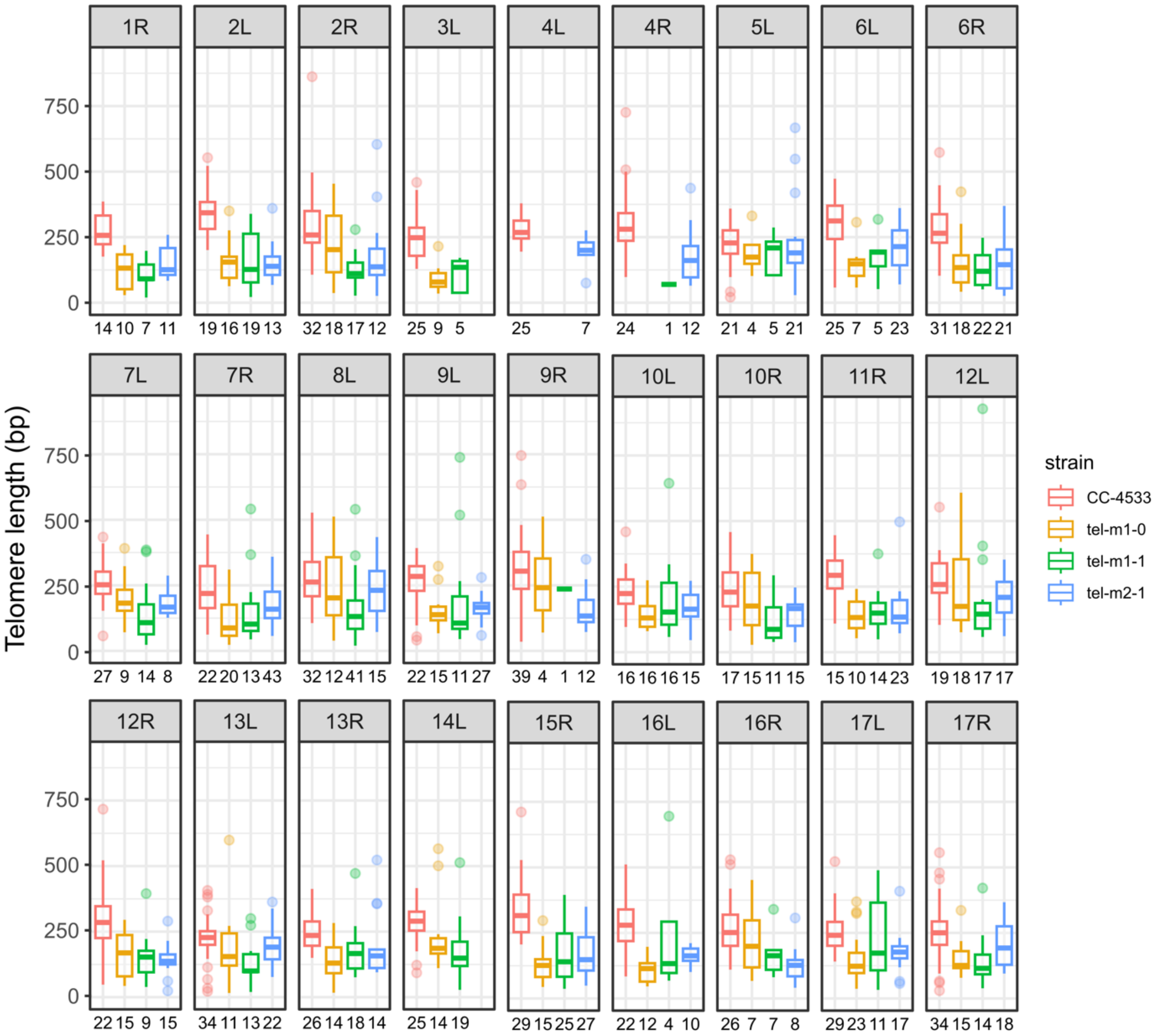


**Supplementary Figure 2. Telomere length distribution for each Sultan subtelomere.** The telomere length distribution of each uniquely identifiable class A and class subtelomere is represented as a boxplot for each indicated strain. The number of corresponding reads is indicated under the boxes.

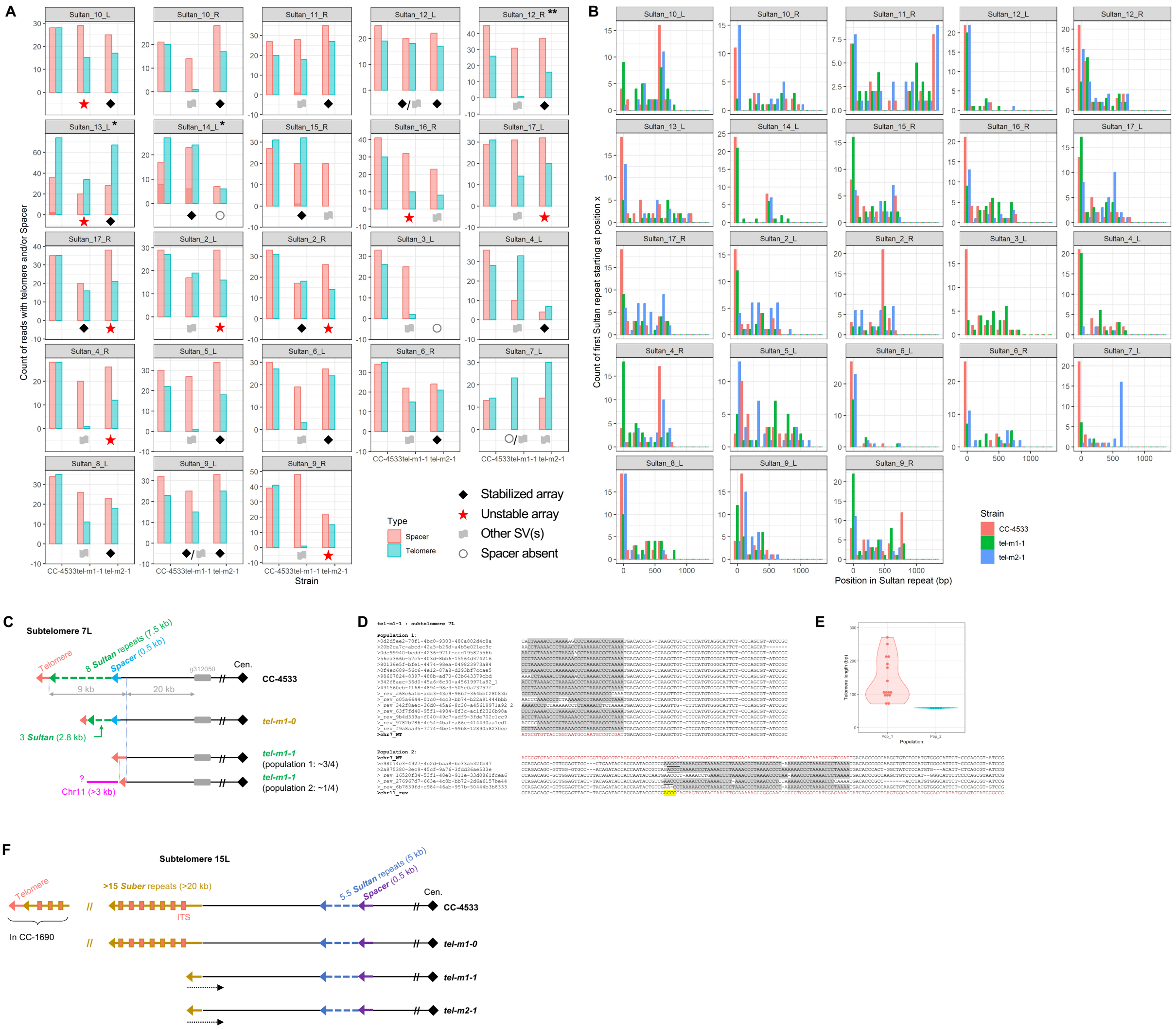


**Supplementary Figure 3. Some Sultan and Suber arrays shorten progressively until complete degradation.** (A) For the indicated class A and B Sultan arrays, the number of reads containing both a telomere sequence and Sultan elements is indicated as a blue bar. The number of reads containing both the Spacer element and Sultan elements is shown as a red bar. In CC-4533, the near equality between these two numbers for most subtelomeres strongly suggests that each of these subtelomeres, starts with a Spacer and ends with a telomere sequence at the individual DNA molecule level. It is mostly the case also for stabilized Sultan arrays in the mutants, marked by a black diamond. In contrast, unstable Sultan arrays (red star) nearly always show more Spacer-Sultan reads than telomere-Sultan reads, suggesting that the chromosome extremity ends with a Sultan element. Subtelomeres with additional SVs (gray flag) cannot be subjected to this analysis. Circles indicate the partial loss or complete absence of Spacer sequence. *For 13L and 14L, the Sultan element was found associated with 2 distinct Spacer elements (light and dark red parts of the histogram); besides, both subtelomeres are from class B, thus their Sultan elements contain insertions that might map elsewhere in the genome close to ITS, hence the high number of telomeres associated with the Sultan. **Subtelomere 12R contains two identical Spacer sequences, thus increasing the count of Sultan reads associated with a Spacer (see also Supp. Dataset 1). (B) Distribution of the position of Sultan-telomere transition in the last Sultan for stabilized Sultan arrays or of the last Sultan element in unstable arrays, in CC-4533, *tel-m1-1* and *tel-m2-1*. (C) Schematic representation of subtelomere 7L in CC-4533, *tel-m1-0* and *tel-m1-1*, showing a decrease in the number of Sultans in *tel-m1-0* compared to CC-4533 and a total loss of all Sultans and the Spacer sequence in *tel-m1-1*, with the formation of a new telomere and even the additional translocation of a sequence from chromosome 11 in a subpopulation. (D) Chromosome 7 to telomere junction in subpopulation 1 of reads and chromosome 7 to ITS to chromosome 11 junctions in subpopulation 2 of reads, as defined in (C). The last junction involves a microhomology of 4 bp (highlighted in yellow). (E) Telomere length distribution of the terminal telomere in subpopulation 1 and of the ITS in subpopulation 2, as defined in (C). (F) Schematic representation of class C subtelomere 15L displaying a structure containing Suber elements in *tel-m1-0* similar to CC-4533, which presumably ends with telomeres as shown for CC-1690[24], but with only one partial Suber remaining in *tel-m1-1* and *tel-m2-1*, suggesting a progressive loss of Suber elements.

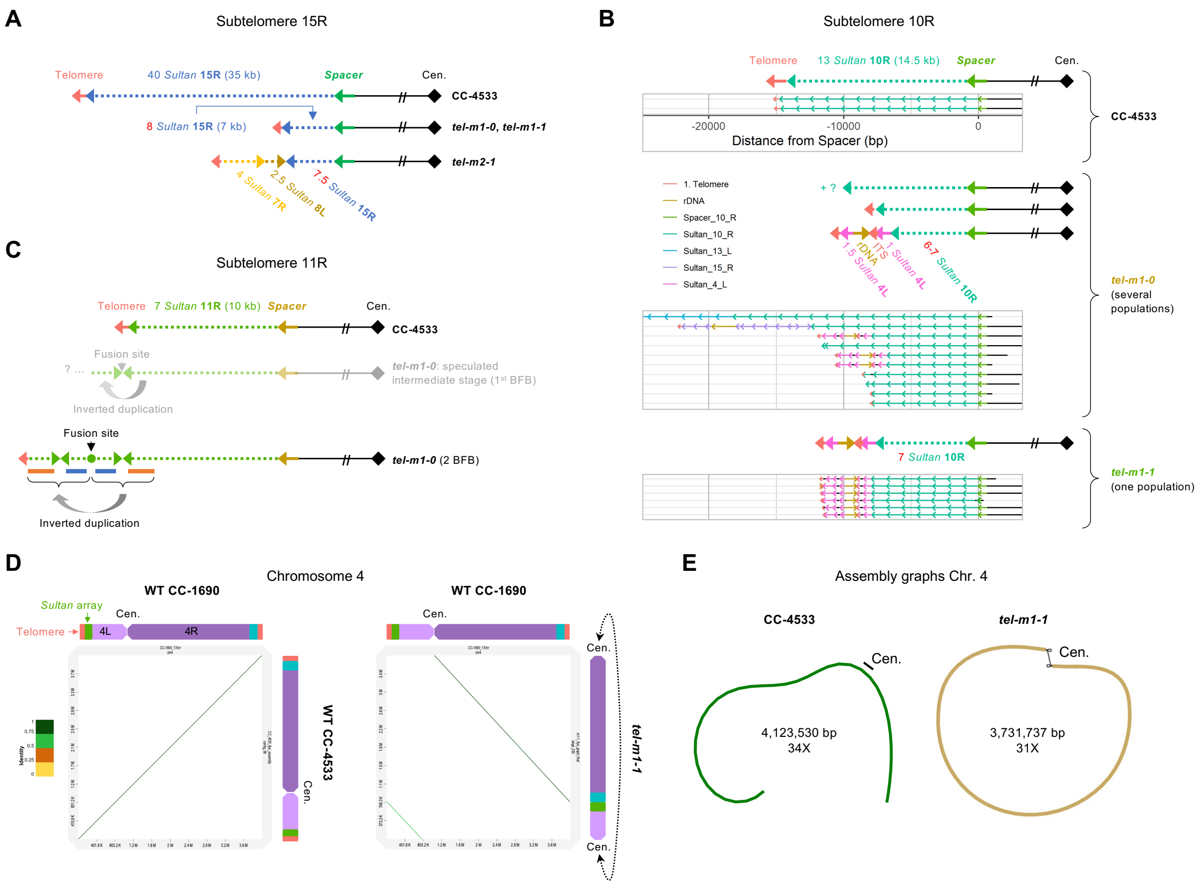


**Supplementary Figure 4. Complex multistep rearrangements at chromosome extremities.** (A) Schematic representation of subtelomere 15R in CC-4533, *tel-m1-0*, *tel-m1-1* and *tel-m2-1*. (B) Schematic representation of subtelomere 10R and reads supporting the depicted structures. Several subpopulations are present in *tel-m1-0* and after purifying selection, *tel-m1-1* is only comprised of one subpopulation. (C) Schematic representation of subtelomere 11R in tel-m1-0 compared to CC-4533. Signature of BFB with multiple inverted repeats suggests at least 2 cycles of BFB. The speculated intermediate stage is shown in shaded colors. (D) Dot plot representation of the alignment of chromosome 4 between CC-1690 and CC-4533 (left), and CC-1690 and *tel-m1-1* (right). The extremities of chromosome 4 of CC-4533 are no longer present in *tel-m1-1*. (E) Graph representation of chromosome 4 of CC-4533 (linear) and *tel-m1-1* (circular topology).

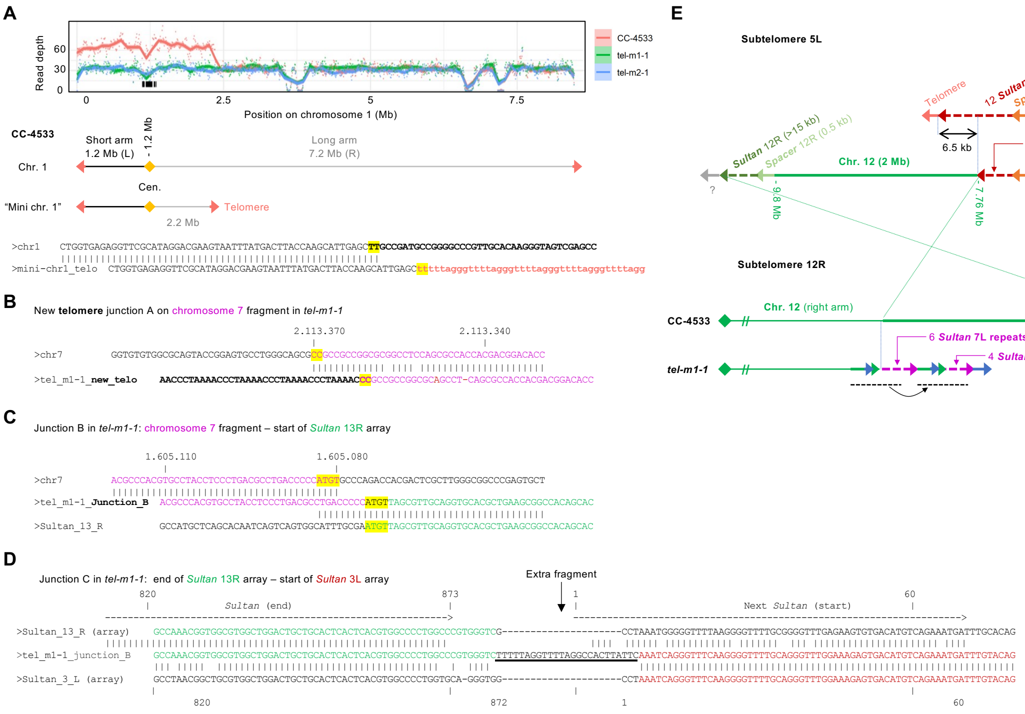


**Supplementary Figure 5. New telomere-capped extremities and junction sequences.** (A) Schematic representation of the mini-chromosome 1 in CC-4533 (middle part), corresponding to a centromere-containing duplicated region of chromosome 1 as shown by the sequencing depth graph (upper part). The junction sequence from mini-chromosome 1 to the new telomere is displayed, with a 2-bp microhomology highlighted in yellow. (B) Sequence of junction A as marked in Fig. 5B, showing a 2-bp microhomology highlighted in yellow. (C) Sequence of junction B as marked in Fig. 5B, with a 4-bp microhomology. (D) Sequence of junction C as marked in Fig. 5B, with the insertion of a short extra fragment between Sultan 13R array and Sultan 3L array, the 2 border Sultan elements being full length.

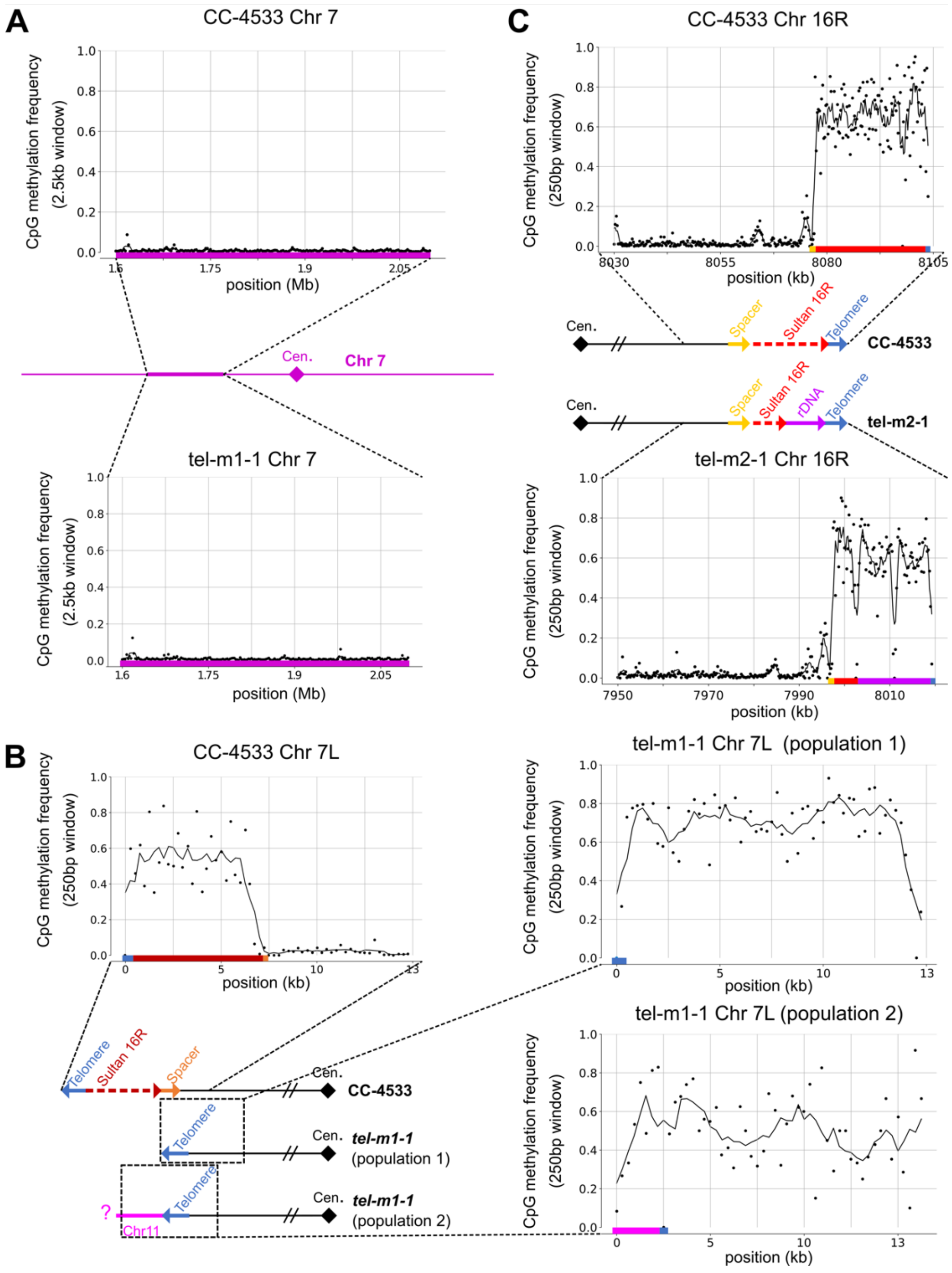


**Supplementary Figure 6. Methylation frequency of other chromosome regions and extremities.** (A-C) Analysis of 5mC frequency at CpG sites using reads that unambiguously spanned the indicated regions, in different illustrative cases. (A) At the original region of chromosome 7 that was duplicated in *tel-m1-1*, shown here for CC-4533 and *tel-m1-1*. (B) At subtelomere 16R where in *tel-m2-1*, the structure shows a shorter hypermethylated Sultan array and a hypermethylated rDNA sequence capped by a telomere, compared to CC-4533 where only the hypermethylated Sultan array is present.

(D) At the new 7L extremity of *tel-m1-1* where methylation spreads further towards the interior of the genome compared to CC-4533 where the hypermethylated domain stops at the Spacer sequence.

**Supp. Dataset 1. Spacer-based analysis of subtelomeres.** All Spacer-containing reads anchored at the Spacer element are depicted for each subtelomere for CC-4533, *tel-m1-0*, *tel-m1-1* and *tel-m2-1*.
