## Supplementary Dataset 1 for "Telomerase-independent survival leads to a mosaic of complex subtelomere rearrangements in *Chlamydomonas reinhardtii*"

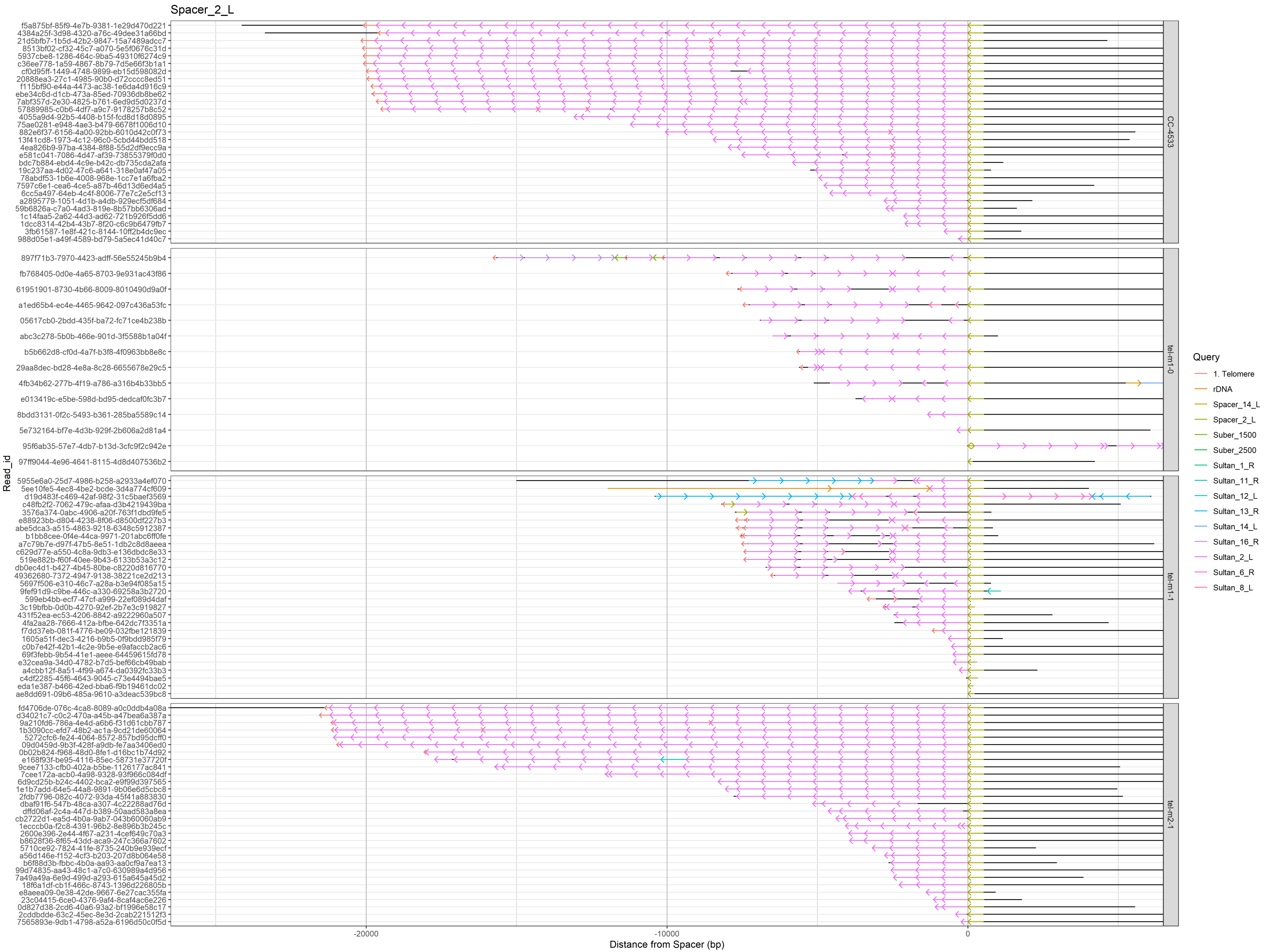

Spacer_2_L
CC-4533
tel-m1-0
tel-m1-1
tel-m2-1
Read_id
Distance from Spacer (bp)
-20000
-10000
0
Query
1. Telomere
rDNA
Spacer_14_L
Spacer_2_L
Suber_1500
Suber_2500
Sultan_1_R
Sultan_11_R
Sultan_12_L
Sultan_13_R
Sultan_14_L
Sultan_16_R
Sultan_2_L
Sultan_6_R
Sultan_8_L

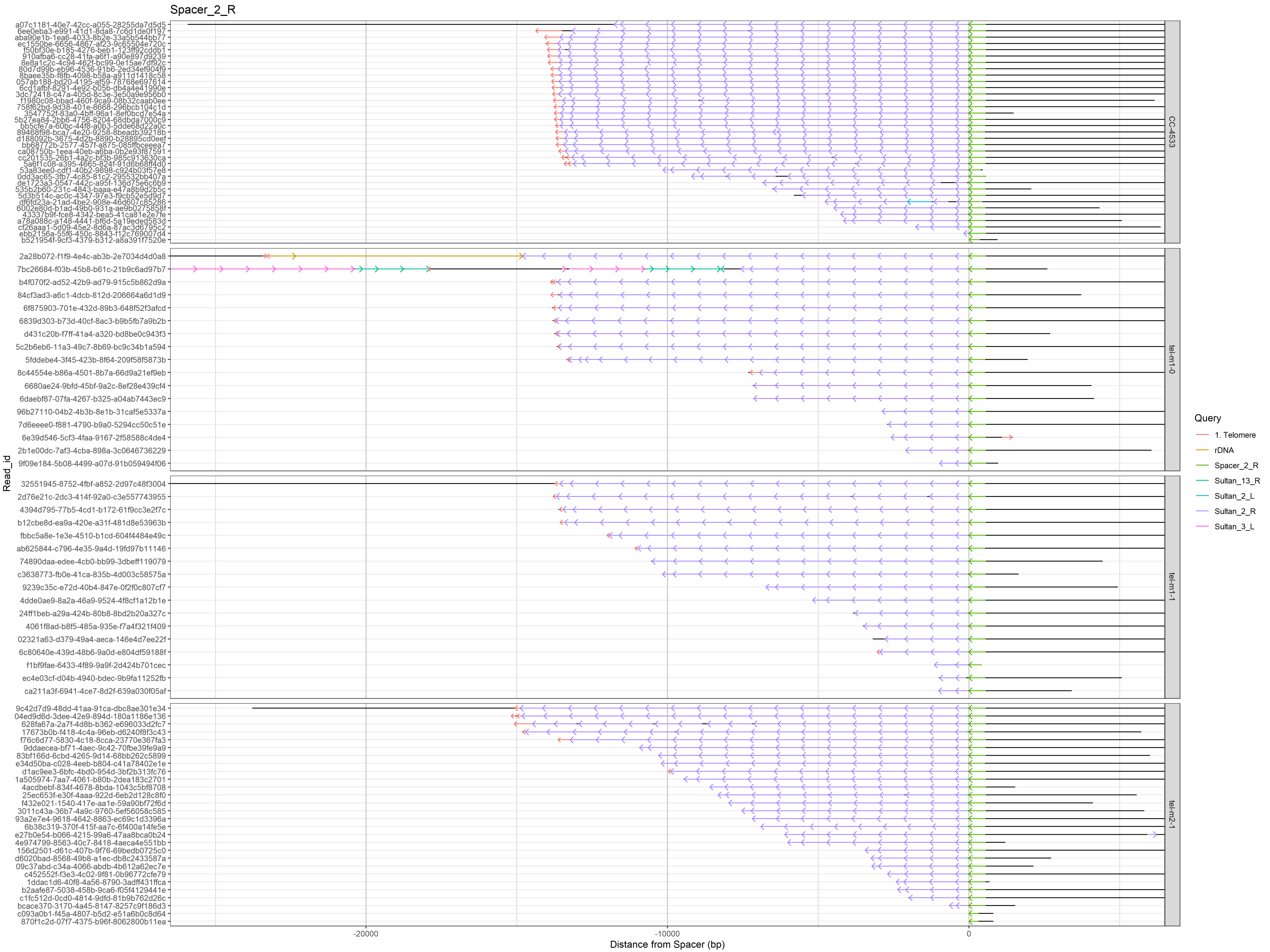

Spacer_2_R
Read_id
Distance from Spacer (bp)
-20000
-10000
0
CC-4533
tel-m1-0
tel-m1-1
tel-m2-1
Query
1. Telomere
rDNA
Spacer_2_R
Sultan_13_R
Sultan_2_L
Sultan_2_R
Sultan_3_L

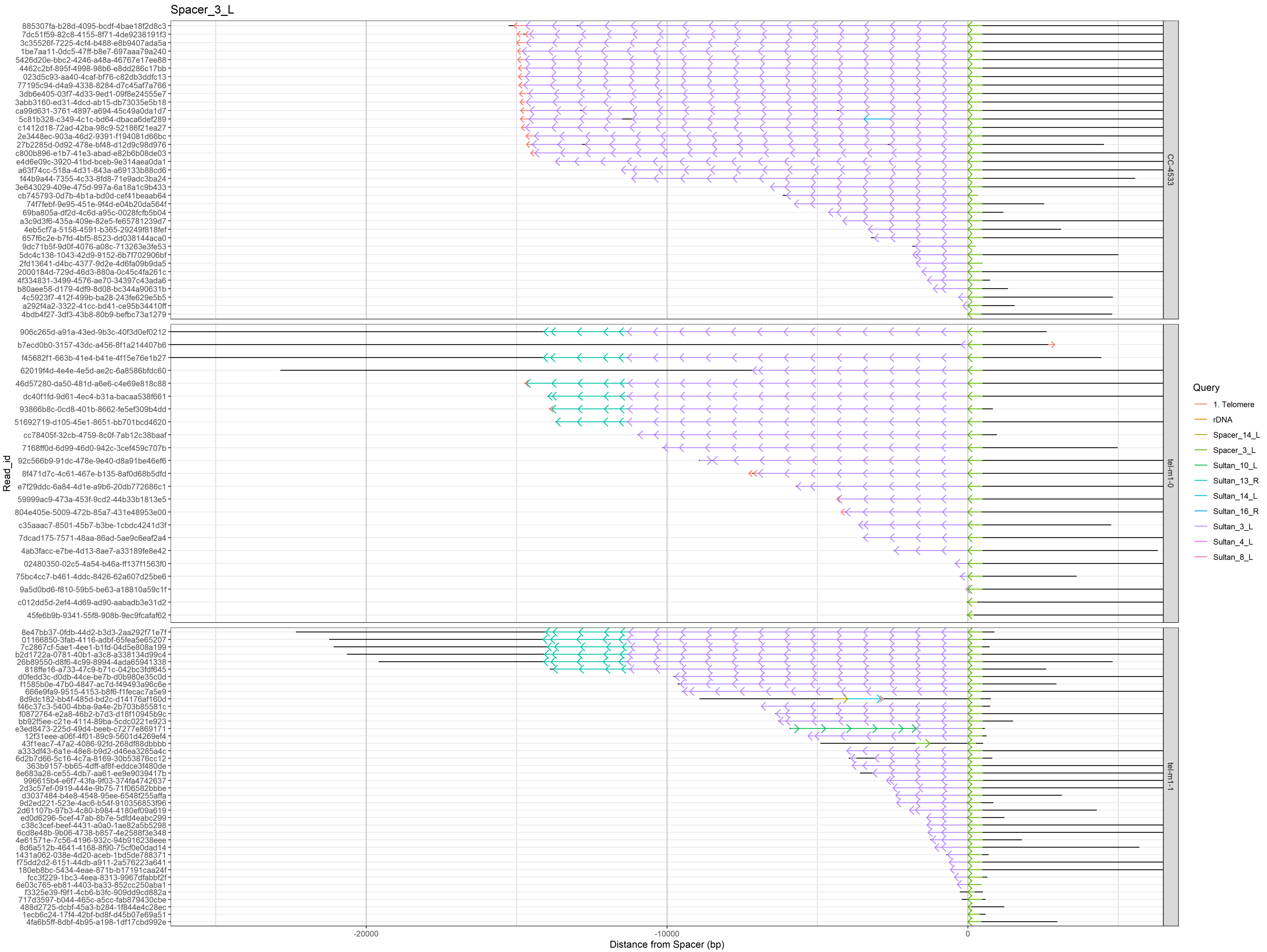
Spacer_3_L
Read_id
Distance from Spacer (bp)
-20000
-10000
0
CC-4533
tel-m1-0
tel-m1-1
Query
1. Telomere
rDNA
Spacer_14_L
Spacer_3_L
Sultan_10_L
Sultan_13_R
Sultan_14_L
Sultan_16_R
Sultan_3_L
Sultan_4_L
Sultan_8_L

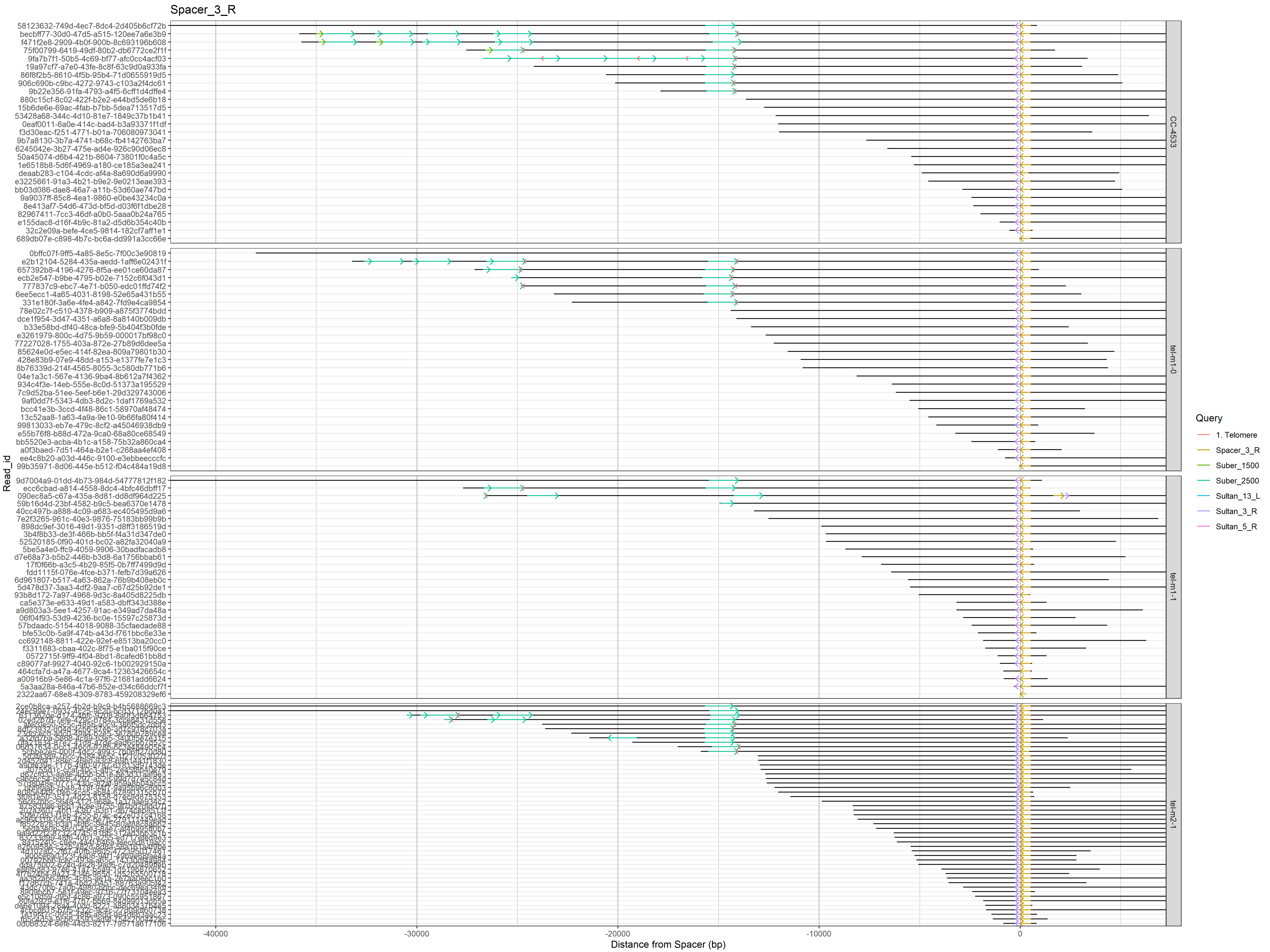

Spacer_3_R
Read_id
Distance from Spacer (bp)
-40000
-30000
-20000
-10000
0
CC-4533
tel-m1-0
tel-m1-1
tel-m2-1
Query
1. Telomere
Spacer_3_R
Suber_1500
Suber_2500
Sultan_13_L
Sultan_3_R
Sultan_5_R

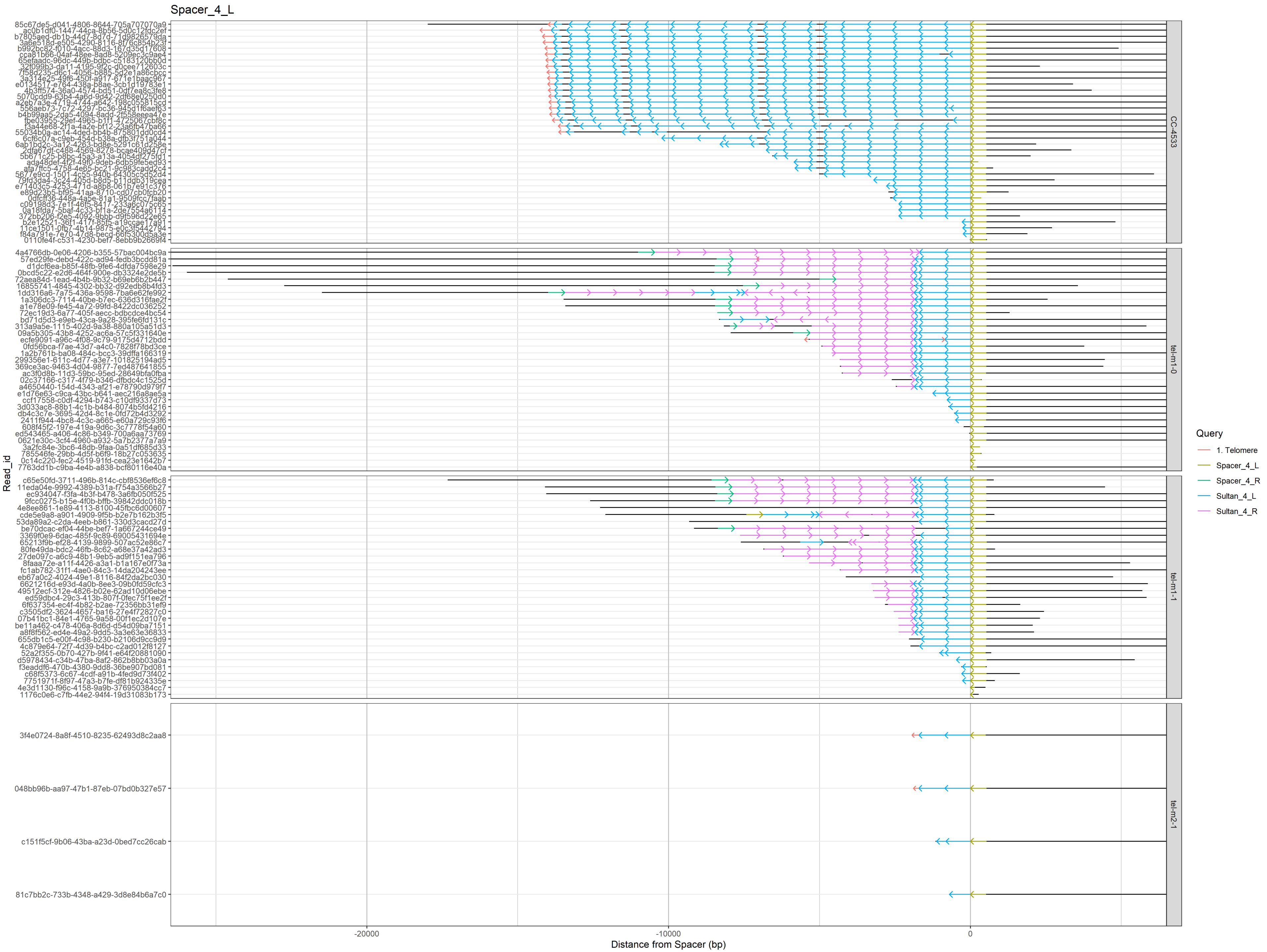

Spacer_4_L
Read_id
Distance from Spacer (bp)
-20000
-10000
0
Query
1. Telomere
Spacer_4_L
Spacer_4_R
Sultan_4_L
Sultan_4_R
CC-4533
tel-m1-0
tel-m1-1
tel-m2-1
3f4e0724-8a8f-4510-8235-62493d8c2aa8
048bb96b-aa97-47b1-87eb-07bd0b327e57
c151f5cf-9b06-43ba-a23d-0bed7cc26cab
81c7bb2c-733b-4348-a429-3d8e84b6a7c0

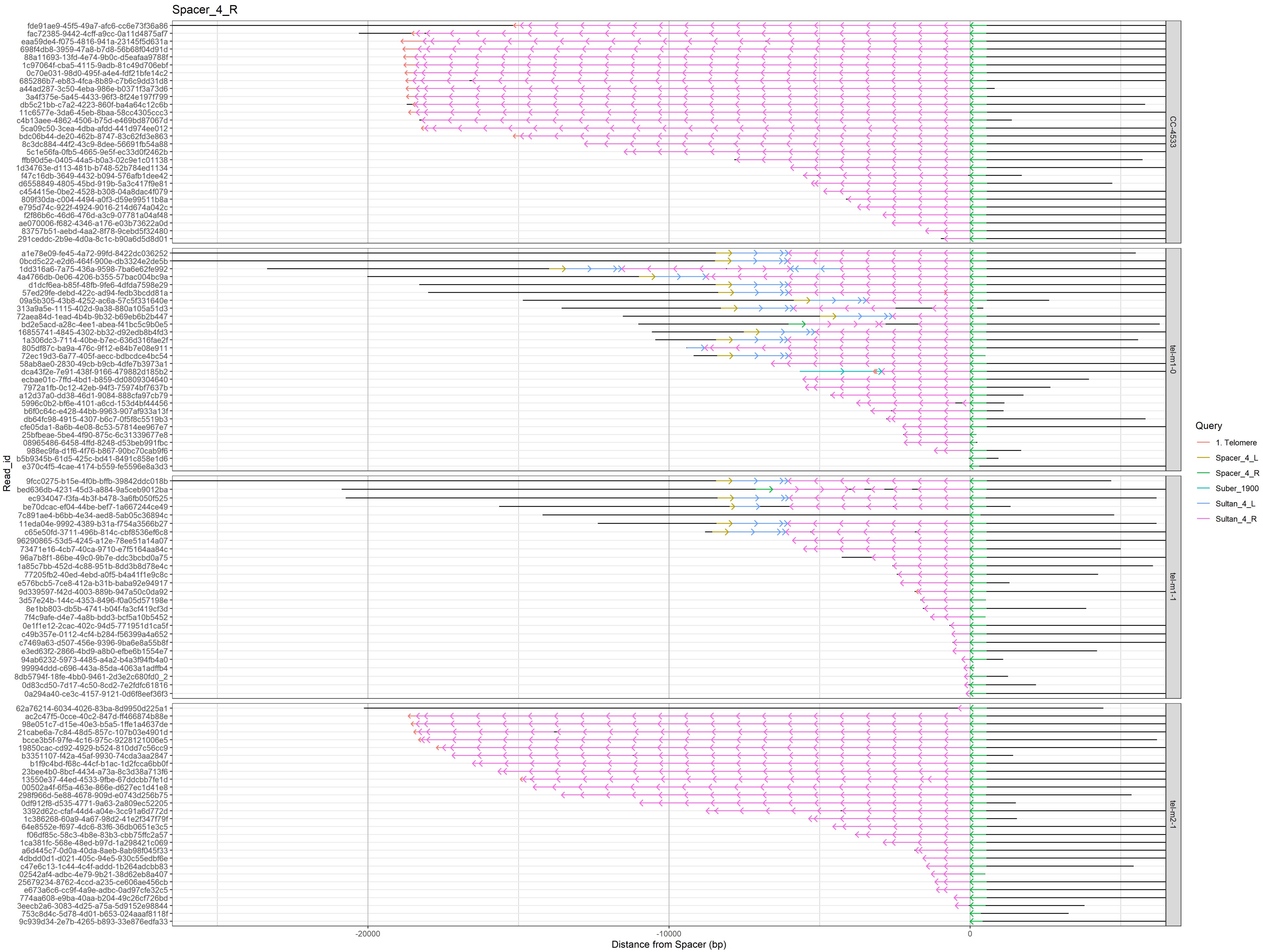

Spacer_4_R
Read_id
Distance from Spacer (bp)
-20000
-10000
0
CC-4533
tel-m1-0
tel-m1-1
tel-m2-1
Query
1. Telomere
Spacer_4_L
Spacer_4_R
Suber_1900
Sultan_4_L
Sultan_4_R

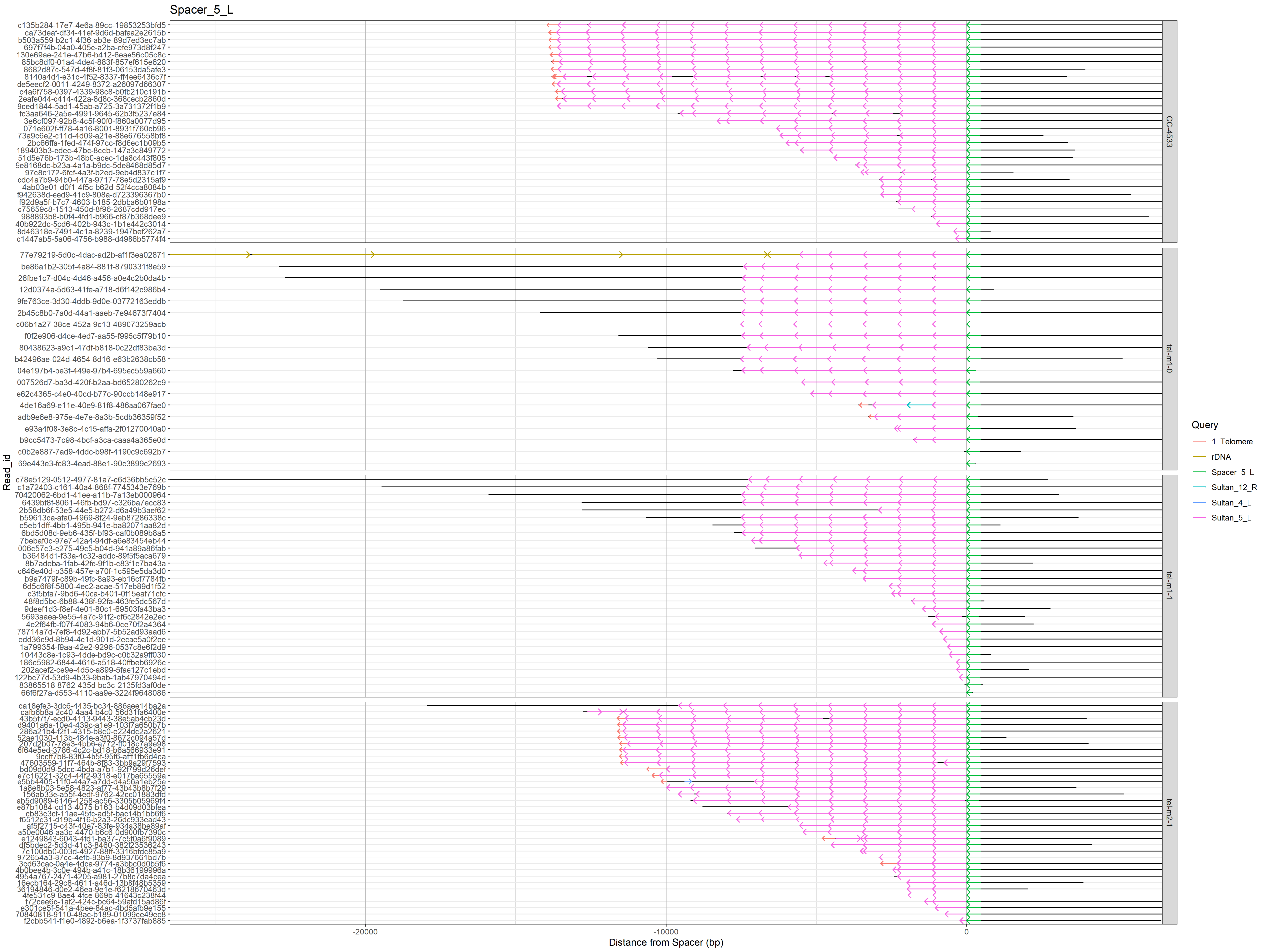

Spacer_5_L
Read_id
Distance from Spacer (bp)
-20000
-10000
0
CC-4533
tel-m1-0
tel-m1-1
tel-m2-1
Query
1. Telomere
rDNA
Spacer_5_L
Sultan_12_R
Sultan_4_L
Sultan_5_L

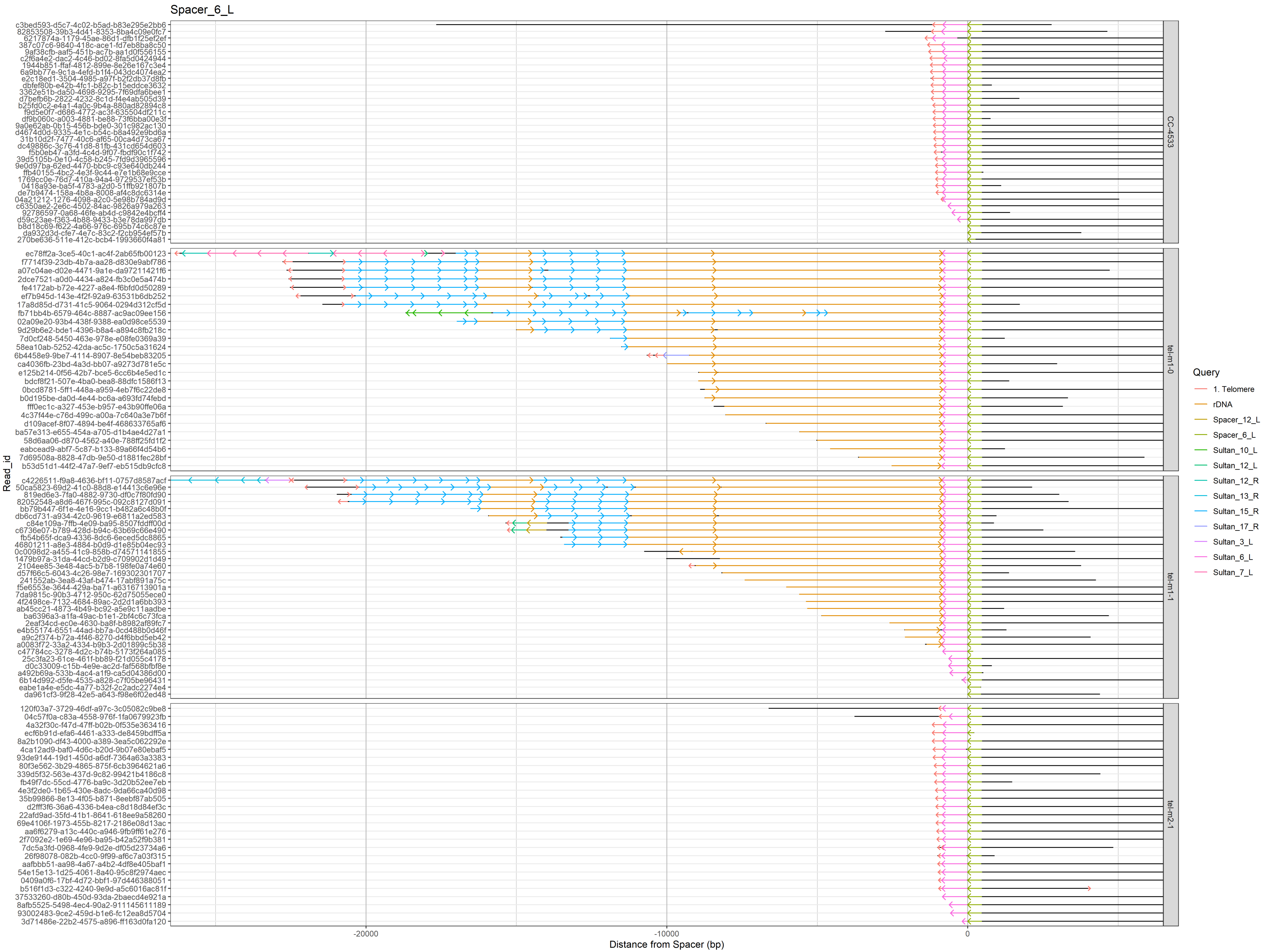

Spacer_6_L
Read_id
Distance from Spacer (bp)
-20000
-10000
0
CC-4533
tel-m1-0
tel-m1-1
tel-m2-1
Query
1. Telomere
rDNA
Spacer_12_L
Spacer_6_L
Sultan_10_L
Sultan_12_L
Sultan_12_R
Sultan_13_R
Sultan_15_R
Sultan_17_R
Sultan_3_L
Sultan_6_L
Sultan_7_L

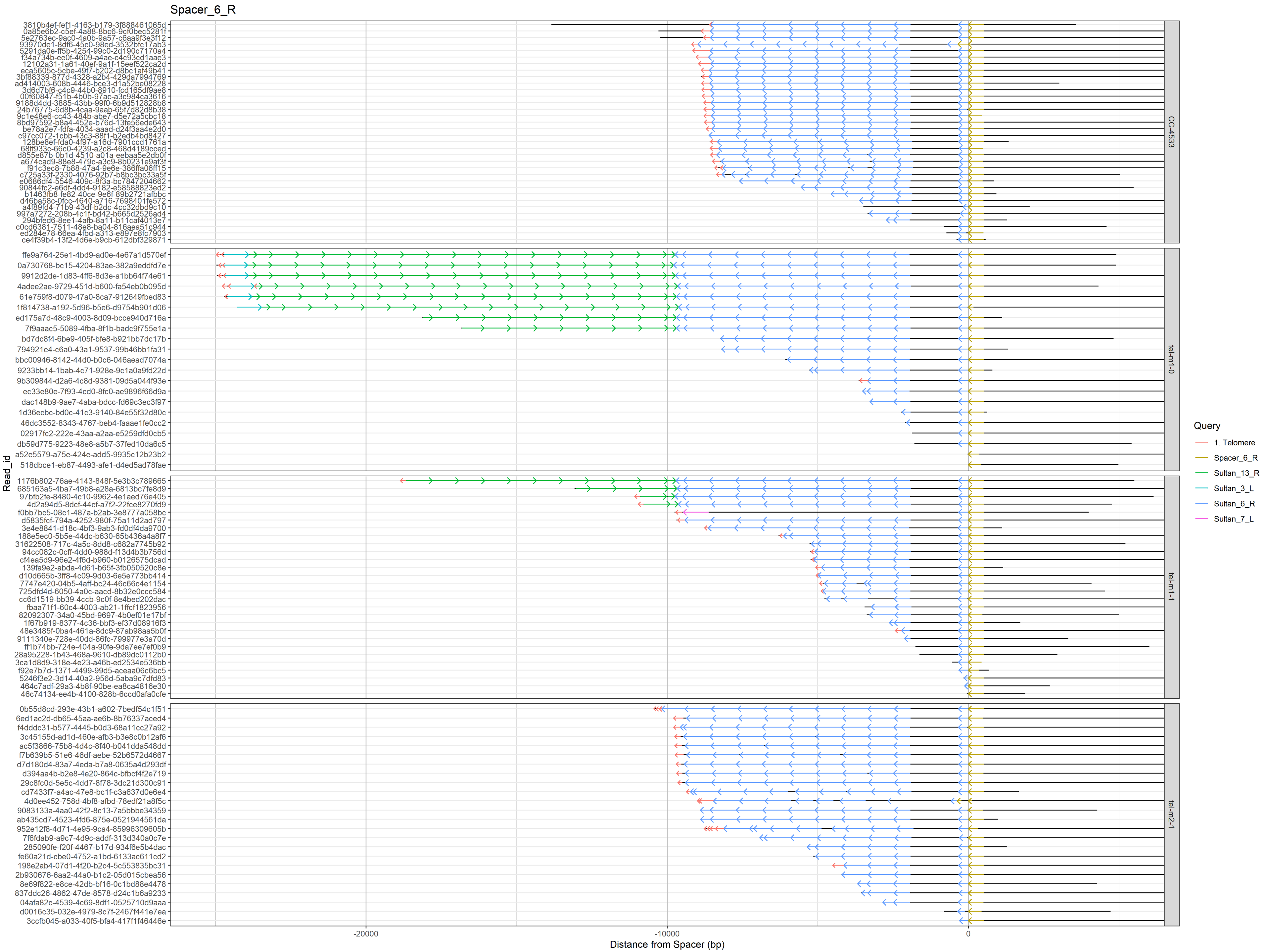

Spacer_6_R
Read_id
Distance from Spacer (bp)
-20000
-10000
0
Query
1. Telomere
Spacer_6_R
Sultan_13_R
Sultan_3_L
Sultan_6_R
Sultan_7_L
CC-4533
tel-m1-0
tel-m1-1
tel-m2-1

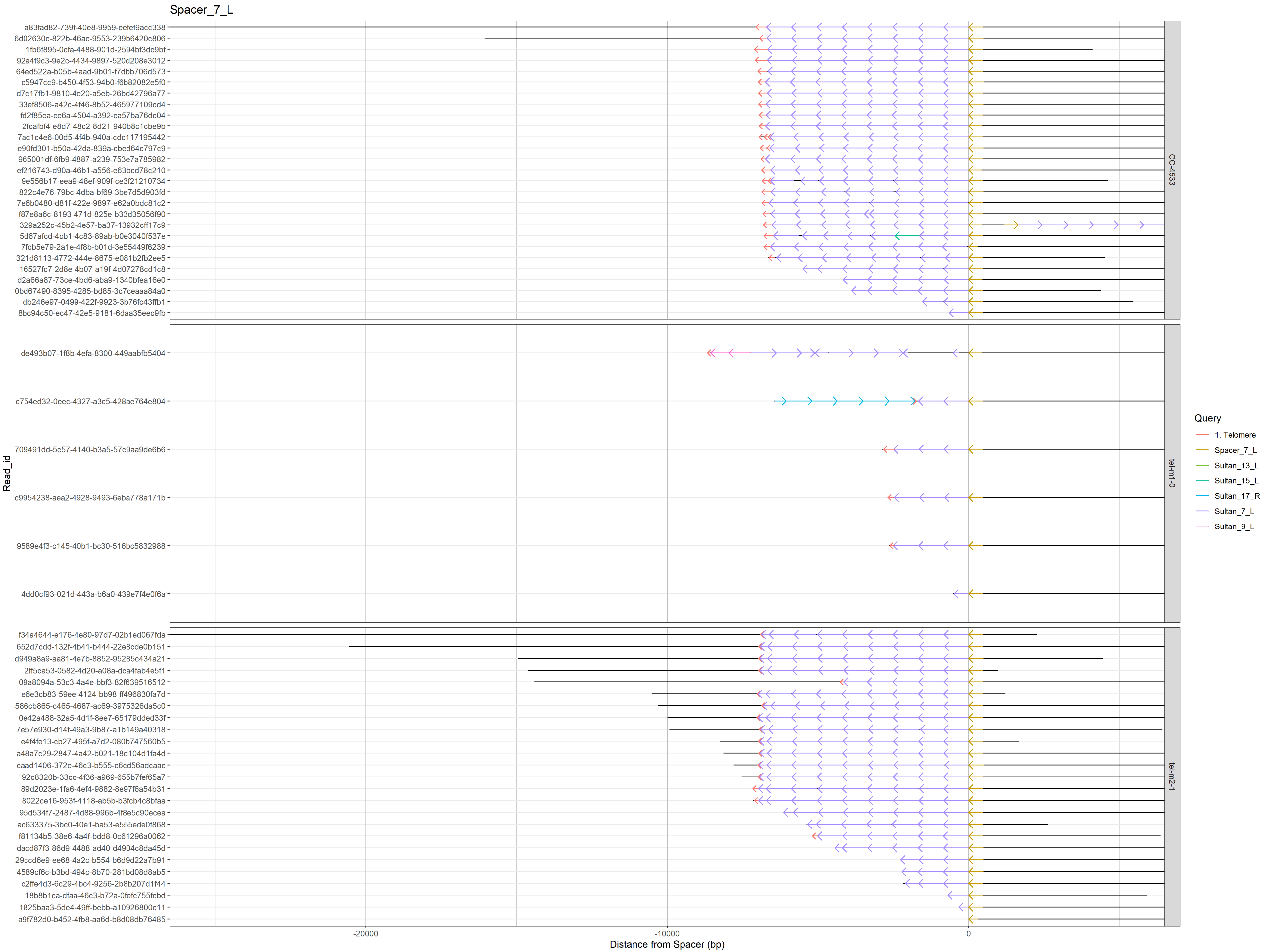

Spacer_7_L
a83fad82-739f-40e8-9959-eefef9acc338
6d02630c-822b-46ac-9553-239b6420c806
1fb6f895-0cfa-4488-901d-2594bf3dc9bf
92a4f9c3-9e2c-4434-9897-520d208e3012
64ed522a-b05b-4aad-9b01-f7dbb706d573
c5947cc9-b450-4f53-94b0-f6b82082e5f0
d7c17fb1-9810-4e20-a5eb-26bd42796a77
33ef8506-a42c-4f46-8b52-465977109cd4
fd2f85ea-ce6a-4504-a392-ca57ba76dc04
2fcafbf4-e8d7-48c2-8d21-940b8c1cbe9b
7ac1c4e6-00d5-4f4b-940a-cdc117195442
e90fd301-b50a-42da-839a-cbed64c797c9
965001df-6fb9-4887-a239-753e7a785982
ef216743-d90a-46b1-a556-e63bcd78c210
9e556b17-eea9-48ef-909f-ce3f21210734
822c4e76-79bc-4dba-bf69-3be7d5d903fd
7e6b0480-d81f-422e-9897-e62a0bdc81c2
f87e8a6c-8193-471d-825e-b33d35056f90
329a252c-45b2-4e57-ba37-13932cff17c9
5d67afcd-4cb1-4c83-89ab-b0e3040f537e
7fcb5e79-2a1e-478b-b01d-3e55449f6239
321d8113-4772-444e-8675-e081b2fb2ee5
16527fc7-2d8e-4b07-a19f-4d07278cd1c8
d2a66a87-73ce-4bd6-aba9-1340bfea16e0
0bd67490-8395-4285-bd85-3c7ceaaa84a0
db246e97-0499-422f-9923-3b76fc43ffb1
8bc94c50-ec47-42e5-9181-6daa35eec9fb
CC-4533
de493b07-1f8b-4efa-8300-449aabfb5404
c754ed32-0eec-4327-a3c5-428ae764e804
709491dd-5c57-4140-b3a5-57c9aa9de6b6
c9954238-aea2-4928-9493-6eba778a171b
9589e4f3-c145-40b1-bc30-516bc5832988
4dd0cf93-021d-443a-b6a0-439e7f4e0f6a
tel-m1-0
f34a4644-e176-4e80-97d7-02b1ed067fda
652d7cdd-132f-4b41-b444-22e8cde0b151
d949a8a9-aa81-4e7b-8852-95285c434a21
2ff5ca53-0582-4d20-a08a-dca4fab4e5f1
09a8094a-53c3-4a4e-bbf3-82f639516512
e6e3cb83-59ee-4174-bb98-ff496830fa7d
586cb865-c465-4687-ac69-3975326da5c0
0e42a488-32a5-4d1f-8ee7-65179dded33f
7e57e930-d14f-49a3-9b87-a1b149a40318
e4f4fe13-cb27-495f-a7d2-080b747560b5
a48a7c29-2847-4a42-b021-18d104d1fa4d
caad1406-372e-46c3-b555-c6cd56adcaac
92c8320b-33cc-4f36-a969-655b7fef65a7
89d2023e-1fa6-4ef4-9882-8e97f6a54b31
8022ce16-953f-4118-ab5b-b3fcb4c8bfaa
95d534f7-2487-4d88-996b-4f8e5c90ecea
ac633375-3bc0-40e1-ba53-e555ede0f868
f81134b5-38e6-4a4f-bdd8-0c61296a0062
dacd87f3-86d9-4488-ad40-d4904c8da45d
29ccd6e9-ee68-4a2c-b554-b6d9d22a7b91
4589cf6c-b3bd-494c-8b70-281bd08d8ab5
c2ffe4d3-6c29-4bc4-9256-2b8b207d1f44
18b8b1ca-dfaa-46c3-b72a-0fefc755fcbd
1825baa3-5de4-49ff-bebb-a10926800c11
a9f782d0-b452-4fb8-aa6d-b8d08db76485
tel-m2-1
Read_id
-20000
-10000
0
Distance from Spacer (bp)
Query
1. Telomere
Spacer_7_L
Sultan_13_L
Sultan_15_L
Sultan_17_R
Sultan_7_L
Sultan_9_L

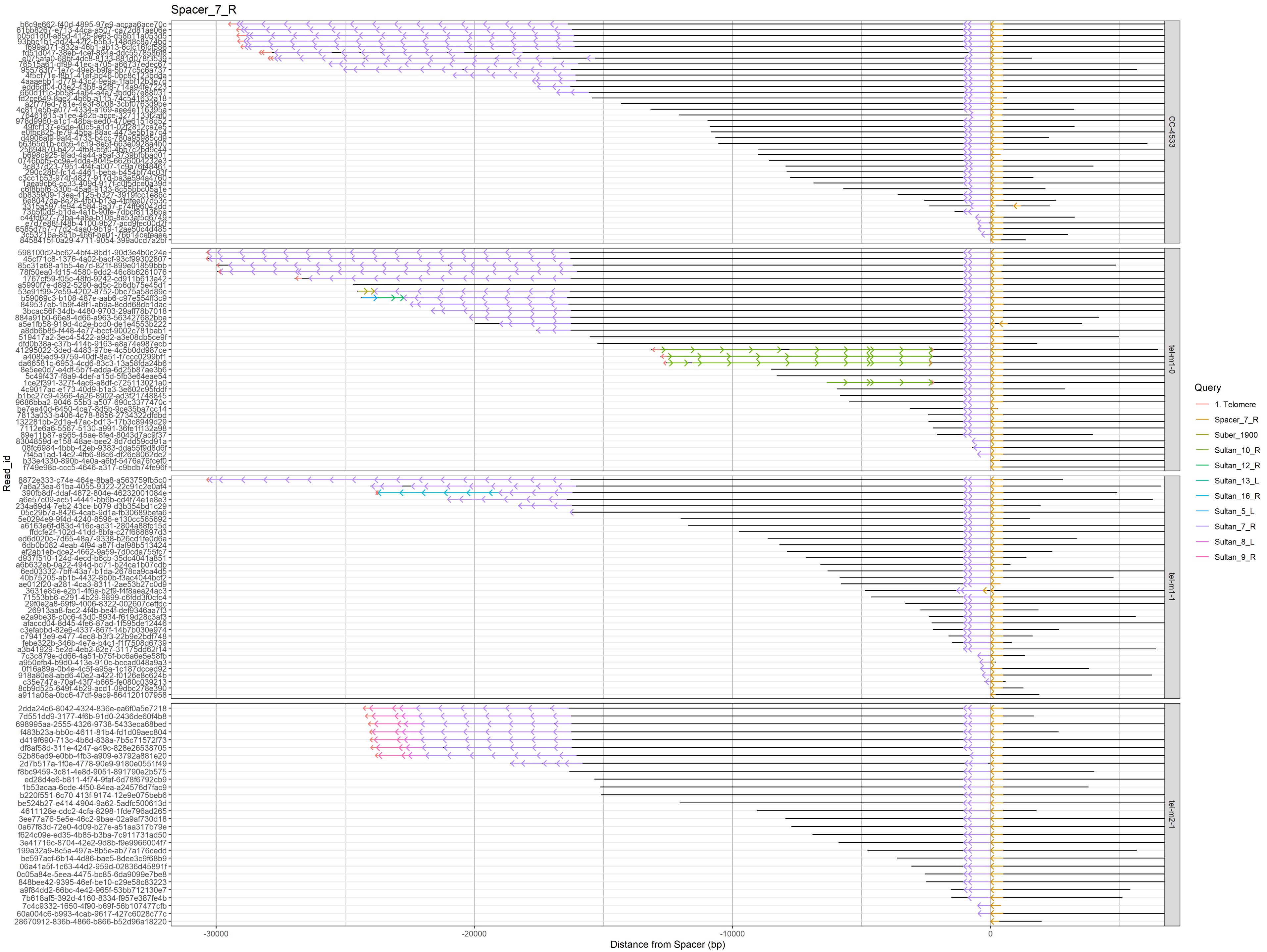

Spacer_7_R
Read_id
Distance from Spacer (bp)
-30000
-20000
-10000
0
CC-4533
tel-m1-0
tel-m1-1
tel-m2-1
Query
1. Telomere
Spacer_7_R
Suber_1900
Sultan_10_R
Sultan_12_R
Sultan_13_L
Sultan_16_R
Sultan_5_L
Sultan_7_R
Sultan_8_L
Sultan_9_R

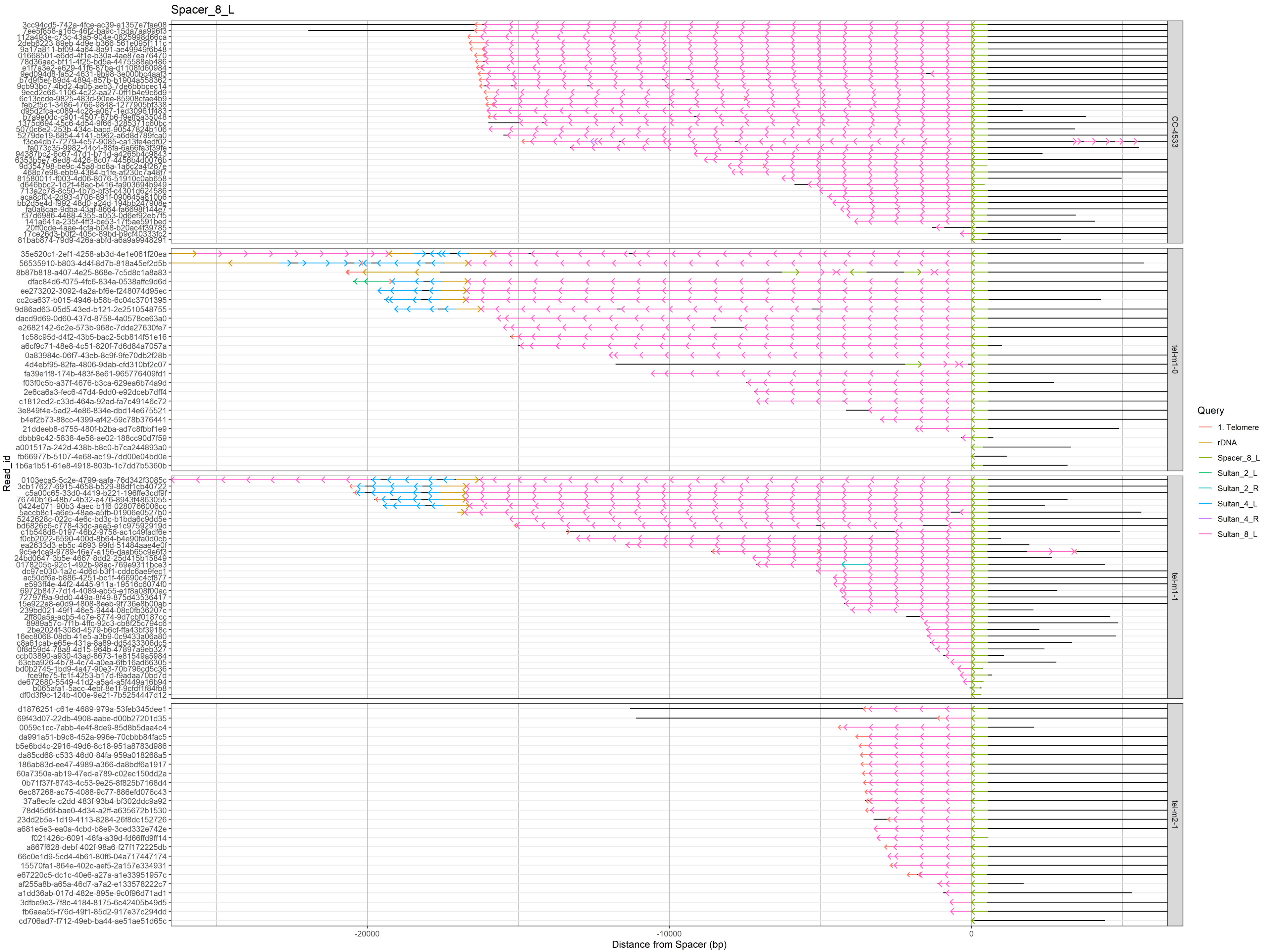
Spacer_8_L
Read_id
Distance from Spacer (bp)
-20000
-10000
0
CC-4533
tel-m1-0
tel-m1-1
tel-m2-1
Query
1. Telomere
rDNA
Spacer_8_L
Sultan_2_L
Sultan_2_R
Sultan_4_L
Sultan_4_R
Sultan_8_L

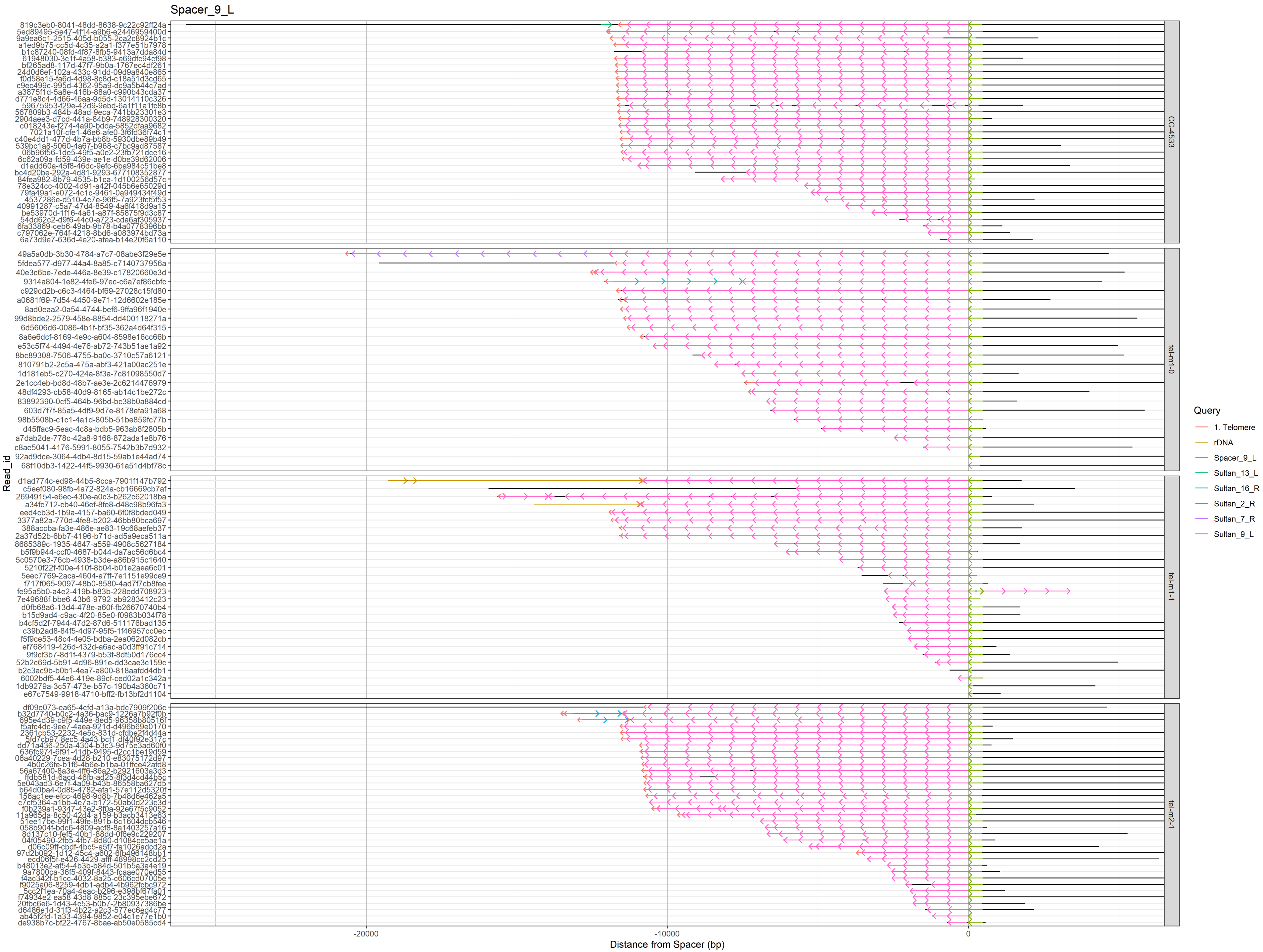

Spacer_9_L
Read_id
Distance from Spacer (bp)
-20000
-10000
0
CC-4533
tel-m1-0
tel-m1-1
tel-m2-1
Query
1. Telomere
rDNA
Spacer_9_L
Sultan_13_L
Sultan_16_R
Sultan_2_R
Sultan_7_R
Sultan_9_L

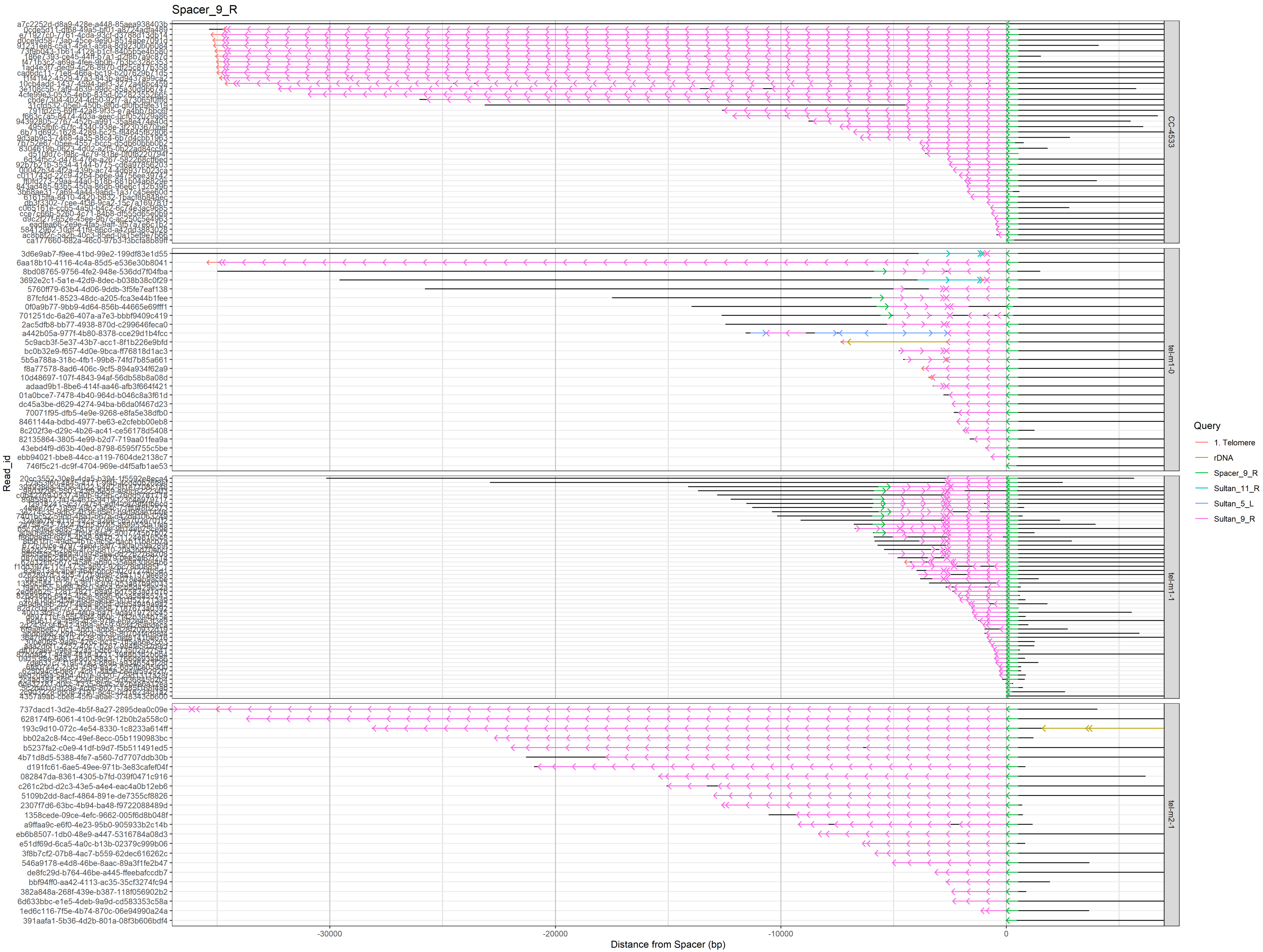

Spacer_9_R
Read_id
Distance from Spacer (bp)
-30000
-20000
-10000
0
CC-4533
tel-m1-0
tel-m1-1
tel-m2-1
Query
1. Telomere
rDNA
Spacer_9_R
Sultan_11_R
Sultan_5_L
Sultan_9_R
3d6e9ab7-f9ee-41bd-99e2-199df83e1d55
6aa18b10-4116-4c4a-85d5-e536e30b8041
8bd08765-9756-4fe2-948e-536dd7f04fba
3692e2c1-5a1e-42d9-8dec-b038b38c0f29
5760ff79-63b4-4d06-9ddb-3f5fe7eaf138
87fcfd41-8523-48dc-a205-fca3e44b1fee
0f0a9b77-9bb9-4d64-856b-44665e69fff1
701251dc-6a26-407a-a7e3-bbbf9409c419
2ac5dfb8-bb77-4938-870d-c299646feca0
a442b05a-977f-4b80-8378-cce29d1b4fcc
5c9acb3f-5e37-43b7-acc1-8f1b226e9bfd
bc0b32e9-f657-4d0e-9bca-ff76818d1ac3
5b5a788a-318c-4fb1-99b8-74fd7b85a661
f8a77578-8ad6-406c-9cf5-894a934f62a9
10d48697-107f-4843-94af-56db58b8a08d
adaad9b1-8be6-414f-aa46-afb3f664f421
01a0bce7-7478-4b40-964d-b046c8a3f61d
dc45a3be-d629-4274-94ba-b6da0f467d23
70071f95-dfb5-4e9e-9268-e8fa5e38dfb0
8461144a-bdbd-4977-be63-e2cfebb00eb8
8c202f3e-d29c-4b26-ac41-ce56178d5408
82135864-3805-4e99-b2d7-719aa01fea9a
43ebd4f9-d63b-40ed-8798-6595f755c5be
ebb94021-bbe8-44cc-a119-7604de2138c7
746f5c21-dc9f-4704-969e-d4f5afb1ae53
737dacd1-3d2e-4b5f-8a27-2895dea0c09e
628174f9-6061-410d-9c9f-12b0b2a558c0
193c9d10-072c-4e54-8330-1c8233a614ff
bb02a2c8-f4cc-49ef-8ecc-05b1190983bc
b5237fa2-c0e9-41df-b9d7-f5b511491ed5
4b71d8d5-5388-4fe7-a560-7d7707ddb30b
d191fc61-6ae5-49ee-971b-3e83cafef04f
082847da-8361-4305-b7fd-039f0471c916
c261c2bd-d2c3-43e5-a4e4-eac4a0b12eb6
5109b2dd-8acf-4864-891e-de7355cf8826
2307f7d6-63bc-4b94-ba48-f9722088489d
1358cede-09ce-4efc-9662-005f6d8b048f
a9ffaa9c-e6f0-4e23-95b0-905933b2c14b
eb6b8507-1db0-48e9-a447-5316784a08d3
e51df69d-6ca5-4a0c-b13b-02379c999b06
3f8b7cf2-07b8-4ac7-b559-62dec616262c
546a9178-e4d8-46be-8aac-89a3f1fe2b47
de8fc29d-b764-46be-a445-ffeebafccdb7
bbf94ff0-aa42-4113-ac35-35cf3274fc94
382a848a-268f-439e-b387-118f056902b2
6d633bbc-e1e5-4deb-9a9d-cd583353c58a
1ed6c116-7f5e-4b74-870c-06e94990a24a
391aafa1-5b36-4d2b-801a-08f3b606bdf4

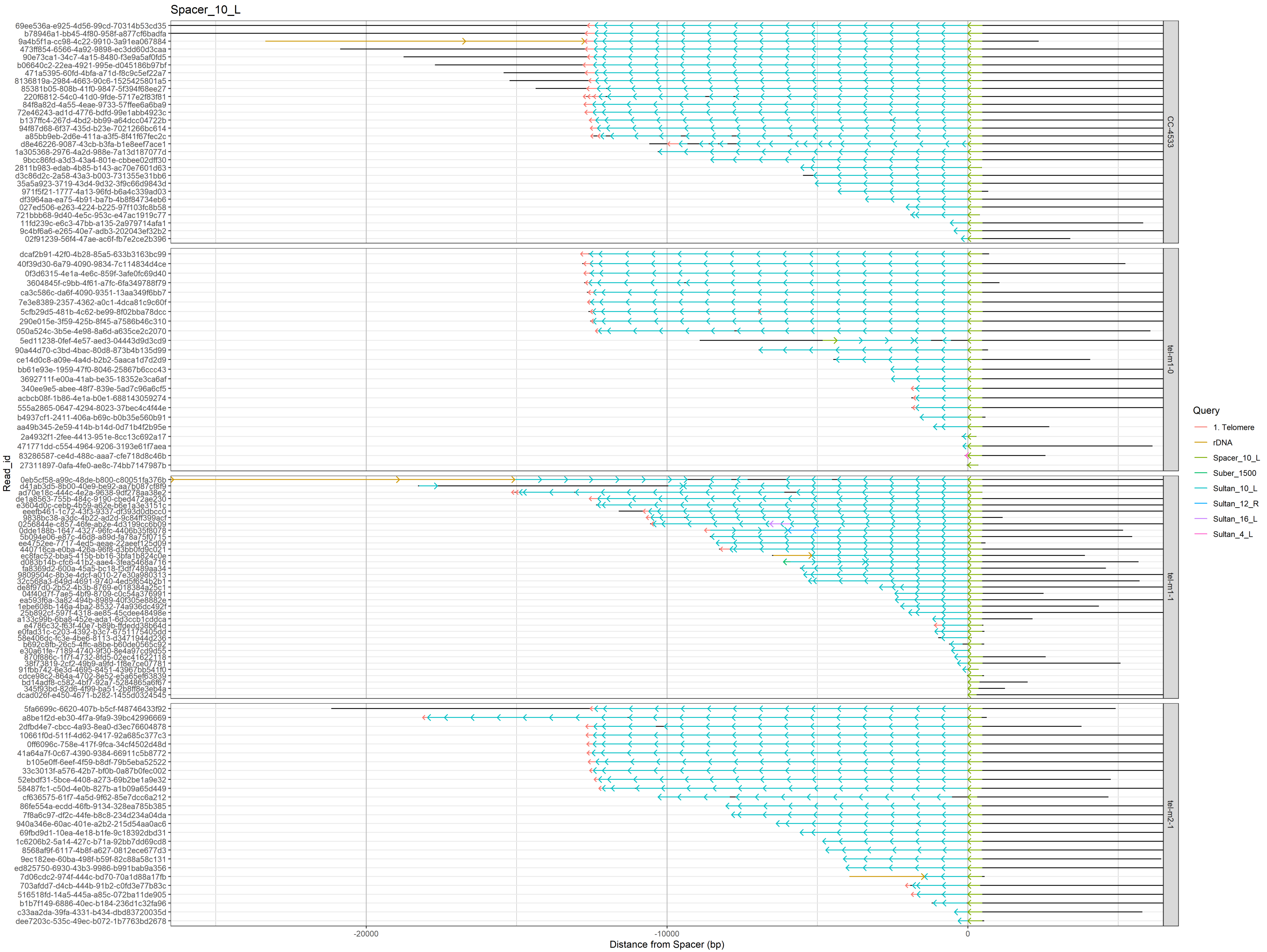

Spacer_10_L
Read_id
Distance from Spacer (bp)
-20000
-10000
0
CC-4533
tel-m1-0
tel-m1-1
tel-m2-1
Query
1. Telomere
rDNA
Spacer_10_L
Suber_1500
Sultan_10_L
Sultan_12_R
Sultan_16_L
Sultan_4_L

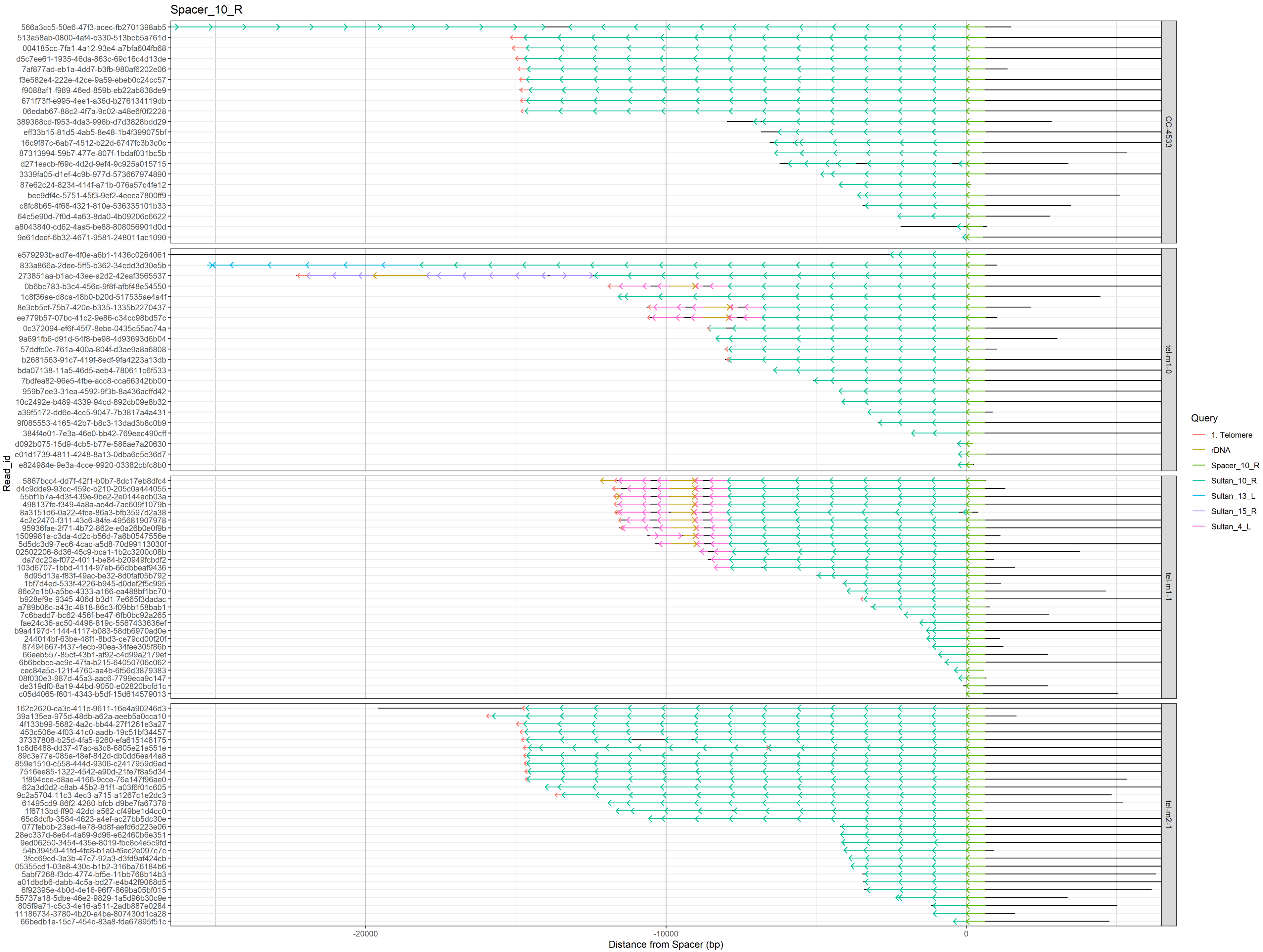

Spacer_10_R
566a3cc5-50e6-47f3-acec-fb2701398ab5
513a58ab-0800-4af4-b330-513bcb5a761d
004185cc-7fa1-4a12-93e4-a7bfa604fb68
d5c7ee61-1935-46da-863c-69c16c4d13de
7af877ad-eb1a-4dd7-b3fb-980af6202e06
f3e582e4-222e-42ce-9a59-ebeb0c24cc57
f9088af1-f989-46ed-859b-eb22ab838de9
671f73ff-e995-4ee1-a36d-b276134119db
06edab67-88c2-4f7a-9c02-a48e6f0f2228
389368cd-f953-4da3-996b-d7d3828bdd29
eff33b15-81d5-4ab5-8e48-1b4f399075bf
16c9f87c-6ab7-4512-b22d-6747fc3b3c0c
87313994-59b7-477e-807f-1bdaf031bc5b
d271eacb-f69c-4d2d-9ef4-9c925a015715
3339fa05-d1ef-4c9b-977d-573667974890
87e62c24-8234-414f-a71b-076a57c4fe12
bec9df4c-5751-45f3-9ef2-4eeca7800ff9
c8fc8b65-4f68-4321-810e-536335101b33
64c5e90d-7f0d-4a63-8da0-4b09206c6622
a8043840-cd62-4aa5-be88-808056901d0d
9e61deef-6b32-4671-9581-248011ac1090
CC-4533
e579293b-ad7e-4f0e-a6b1-1436c0264061
833a866a-2dee-5ff5-b362-34cdd3d30e5b
273851aa-b1ac-43ee-a2d2-42eaf3565537
0b6bc783-b3c4-456e-9f8f-afbf48e54550
1c8f36ae-d8ca-48b0-b20d-517535ae4a4f
8e3cb5cf-75b7-420e-b335-1335b2270437
ee779b57-07bc-41c2-9e86-c34cc98bd57c
0c372094-ef6f-45f7-8ebe-0435c55ac74a
9a691fb6-d91d-54f8-be98-4d93693d6b04
57ddfc0c-761a-400a-804f-d3ae9a8a6808
b2681563-91c7-419f-8edf-9fa4223a13db
bda07138-11a5-46d5-aeb4-780611c6f533
7bdfea82-96e5-4fbe-acc8-cca66342bb00
959b7ee3-31ea-4592-9f3b-8a436acffd42
10c2492e-b489-4339-94cd-892cb09e8b32
a39f5172-dd6e-4cc5-9047-7b3817a4a431
9f085553-4165-42b7-b8c3-13dad3b8c0b9
384f4e01-7e3a-46e0-bb42-769eec490cff
d092b075-15d9-4cb5-b77e-586ae7a20630
e01d1739-4811-4248-8a13-0dba6e5e36d7
e824984e-9e3a-4cce-9920-03382cbfc8b0
tel-m1-0
5867bcc4-dd7f-42f1-b0b7-8dc17eb8dfc4
d4c9dde9-93cc-459c-b210-205c0a444055
55bf1b7a-4d3f-439e-9be2-2e0144acb03a
498137fe-f349-4a8a-ac4d-7ac609f1079b
8a3151d6-0a22-4fca-86a3-bfb3597d2a38
4c2c2470-f311-43c6-84fe-495681907978
95936fae-2f71-4b72-862e-e0a26b0e0f9b
1509981a-c3da-4d2c-b56d-7a8b0547556e
5d5dc3d9-7ec6-4cac-a5d8-70d99113030f
02502206-8d36-45c9-bca1-1b2c3200c08b
da7dc20a-f072-4011-be84-b20949fcbdf2
103d6707-1bbd-4114-97eb-66dbbeaf9436
8d95d13a-f83f-49ac-be32-8d0faf05b792
1bf7d4ed-533f-4226-b945-d0def2f5c995
86e2e1b0-a5be-4333-a166-ea488bf1bc70
b928ef9e-9345-406d-b3d1-7e665f3dadac
a789b06c-a43c-4818-86c3-f09bb158bab1
7c6badd7-bc62-456f-be47-6fb0bc92a265
fae24c36-ac50-4496-819c-5567433636ef
b9a4197d-1144-4117-b083-58db6970ad0e
244014bf-63be-48f1-8bd3-ce79cd00f20f
87494667-f437-4ecb-90ea-34fee305f86b
66eeb557-85cf-43b1-af92-c4d99a2179ef
6b6bcbcc-ac9c-47fa-b215-64050706c062
cec84a5c-121f-4760-aa4b-6f56d3879383
08f030e3-987d-45a3-aac6-7799eca9c147
de319df0-8a19-44bd-9050-e02820bcfd1c
c05d4065-f601-4343-b5df-15d614579013
tel-m1-1
162c2620-ca3c-411c-9611-16e4a90246d3
39a135ea-975d-48db-a62a-aeeb5a0cca10
4f133b99-5682-4a2c-bb44-27f1261e3a27
453c506e-4f03-41c0-aadb-19c51bf34457
37337808-b25d-4fa5-9260-efa615148175
1c8d6488-dd37-47ac-a3c8-6805e21a551e
89c3e77a-085a-48ef-842d-db0dd6ea44a8
859e1510-c558-444d-9306-c2417959d6ad
7516ee85-1322-4542-a90d-21fe7f8a5d34
1f894cce-d8ae-4166-9cce-76a147f96ae0
62a3d0d2-c8ab-45b2-81f1-a03f6f01c605
9c2a5704-11c3-4ec3-a715-a1267c1e2dc3
61495cd9-86f2-4280-bfcb-d9be7fa67378
1f6713bd-ff90-42dd-a562-cf49be1d4cc0
65c8dcfb-3584-4623-a4ef-ac27bb5dc30e
077febbb-23ad-4e78-9d8f-aefd6d223e06
28ec337d-8e64-4a69-9d96-e62460b6e351
9ed06250-3454-435e-8019-fbc8c4e5c9fd
54b39459-41fd-4fe8-b1a0-f6ec2e097c7c
3fcc69cd-3a3b-47c7-92a3-d3fd9af424cb
05355cd1-03e8-430c-b1b2-316ba76184b6
5abf7268-f3dc-4774-bf5e-11bb768b14b3
a01dbdb6-dabb-4c5a-bd27-e4b42f9068d5
6f92395e-4b0d-4e16-96f7-869ba05bf015
55737a18-5dbe-46e2-9829-1a5d96b30c9e
805f9a71-c5c3-4e16-a511-2adb887e0284
11186734-3780-4b20-a4ba-8d7430d1ca28
66bedb1a-15c7-454c-83a8-fda67895f51c
tel-m2-1
Read_id
-20000
-10000
0
Distance from Spacer (bp)
Query
1. Telomere
rDNA
Spacer_10_R
Sultan_10_R
Sultan_13_L
Sultan_15_R
Sultan_4_L

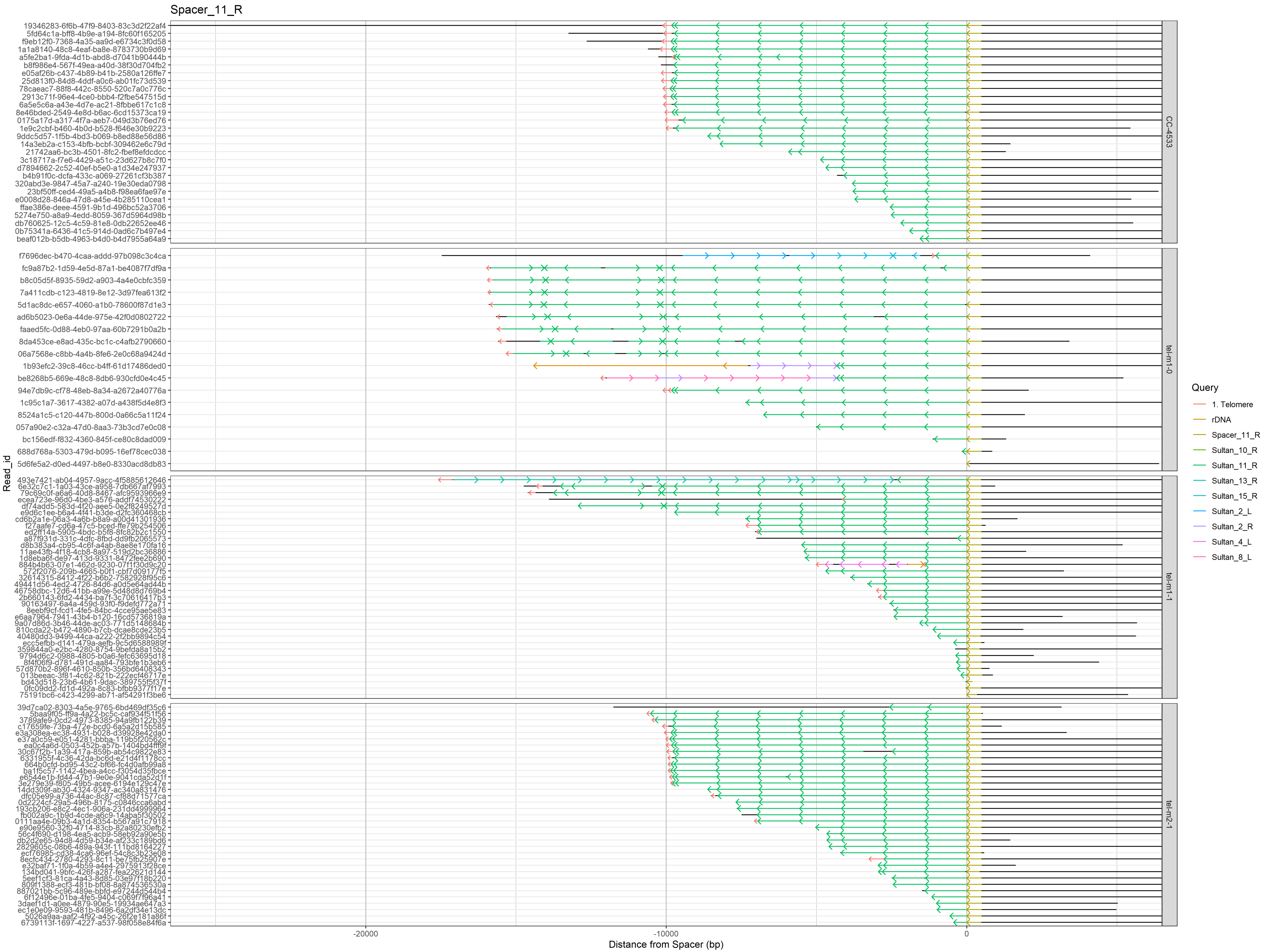

Spacer_11_R
Read_id
Distance from Spacer (bp)
-20000
-10000
0
CC-4533
tel-m1-0
tel-m1-1
tel-m2-1
Query
1. Telomere
rDNA
Spacer_11_R
Sultan_10_R
Sultan_11_R
Sultan_13_R
Sultan_15_R
Sultan_2_L
Sultan_2_R
Sultan_4_L
Sultan_8_L

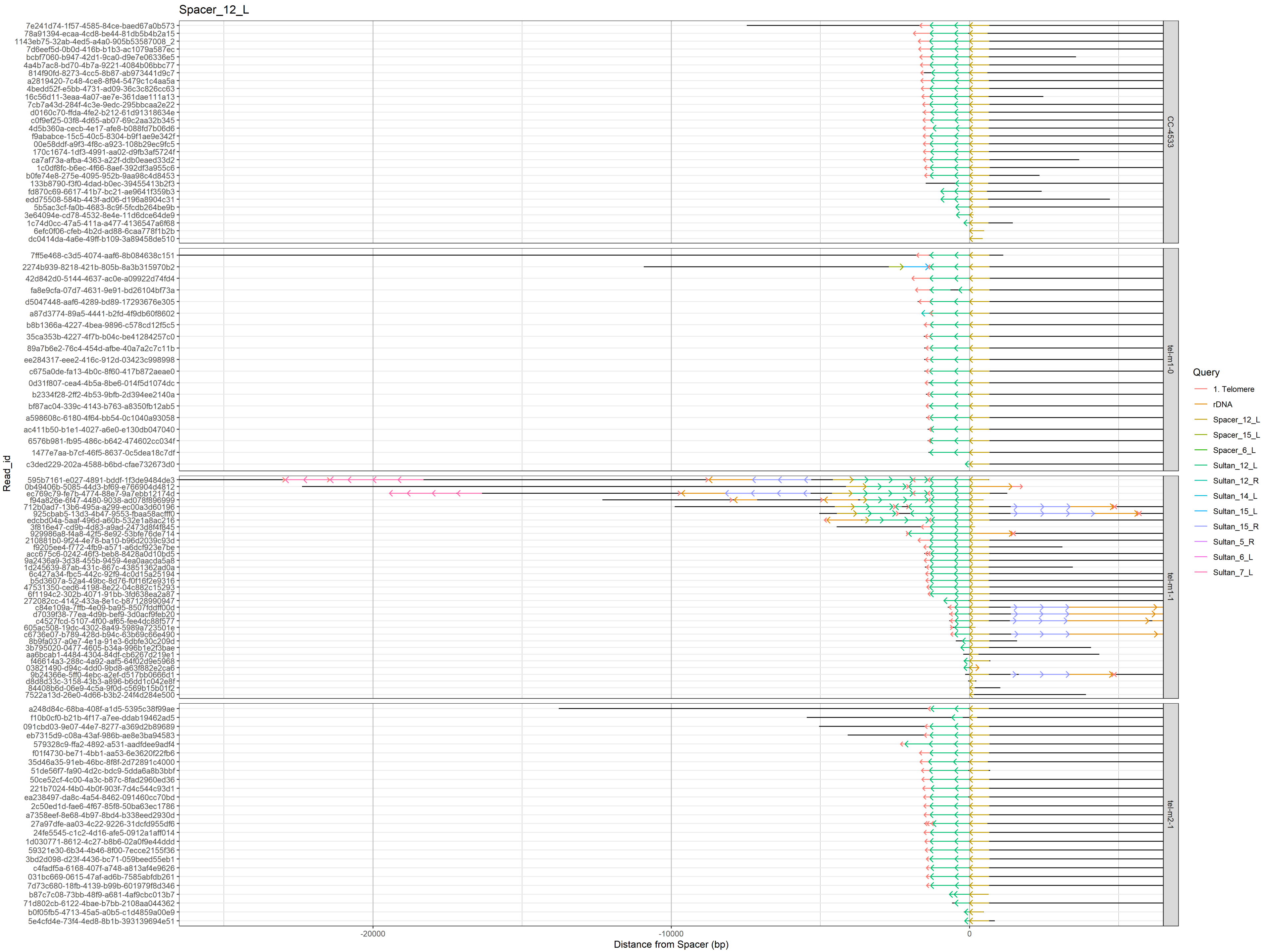

Spacer_12_L
Read_id
Distance from Spacer (bp)
-20000
-10000
0
CC-4533
tel-m1-0
tel-m1-1
tel-m2-1
Query
1. Telomere
rDNA
Spacer_12_L
Spacer_15_L
Spacer_6_L
Sultan_12_L
Sultan_12_R
Sultan_14_L
Sultan_15_L
Sultan_15_R
Sultan_5_R
Sultan_6_L
Sultan_7_L

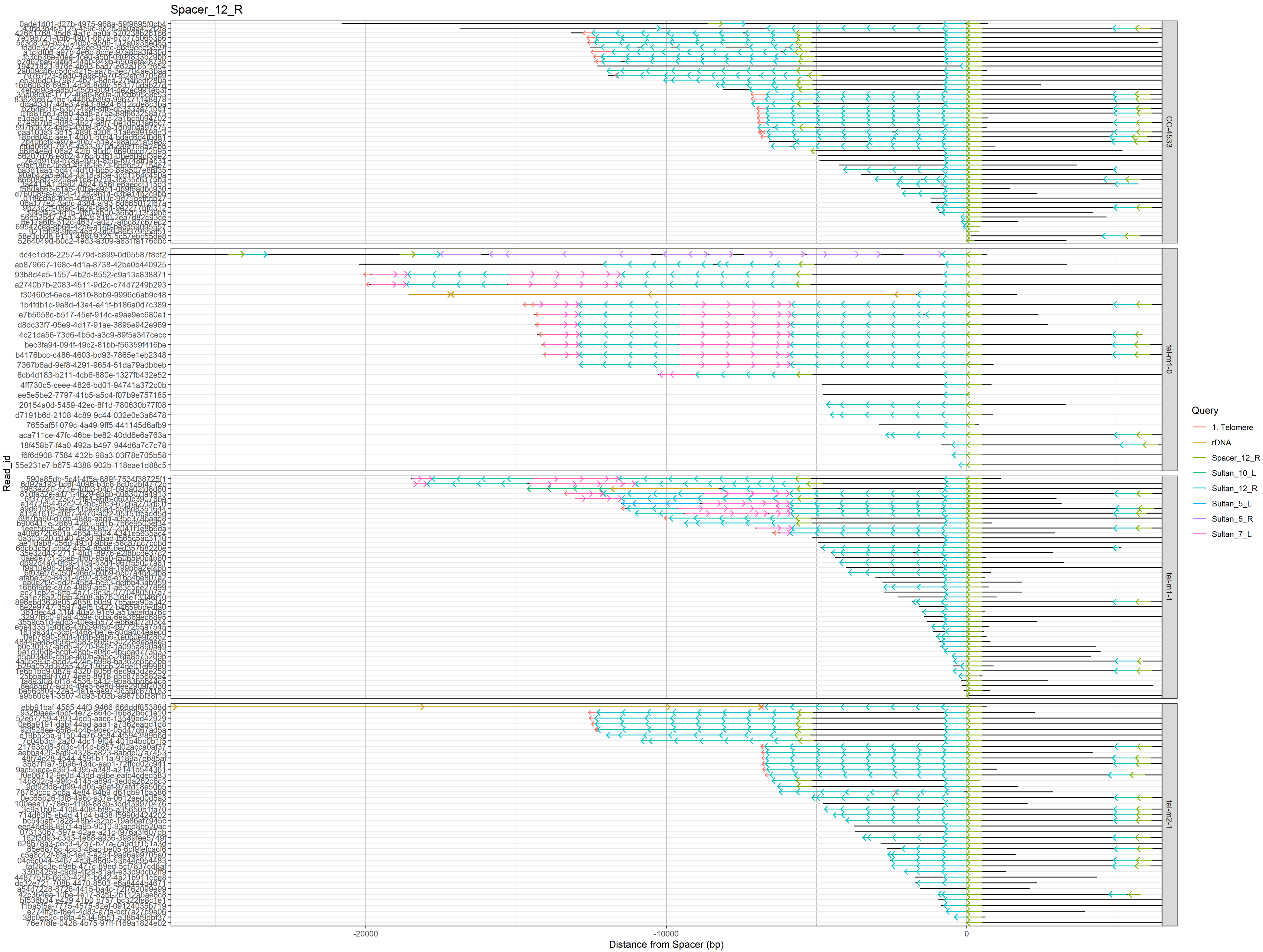

Spacer_12_R
Read_id
Distance from Spacer (bp)
-20000
-10000
0
CC-4533
tel-m1-0
tel-m1-1
tel-m2-1
Query
1. Telomere
rDNA
Spacer_12_R
Sultan_10_L
Sultan_12_R
Sultan_5_L
Sultan_5_R
Sultan_7_L
dc4c1dd8-2257-479d-b899-0d65587f8df2
ab879667-168c-4d1a-8738-42be0b440925
93b8d4e5-1557-4b2d-8552-c9a13e838871
a2740b7b-2083-4511-9d2c-c74d7249b293
f30460cf-6eca-4810-8bb9-9996c6ab9c48
1b4fdb1d-9a8d-43a4-a41f-b186a0d7c389
e7b5658c-b517-45ef-914c-a9ae9ec680a1
d8dc33f7-05e9-4d17-91ae-3895e942e969
4c21da56-73d6-4b5d-a3c9-89f5a347cecc
bec3fa94-094f-49c2-81bb-f56359f416be
b4176bcc-c486-4603-bd93-7865e1eb2348
7367b6ad-9ef8-4291-9654-51da79adbbeb
8cb4d183-b211-4cb6-880e-1327fb432e52
4ff730c5-ceee-4826-bd01-94741a372c0b
ee5e5be2-7797-41b5-a5c4-f07b9e757185
20154a0d-5459-42ec-8f1d-780630b77f08
d7191b6d-2108-4c89-9c44-032e0e3a6478
7655af5f-079c-4a49-9ff5-441145d6afb9
aca711ce-47fc-46be-be82-40dd6e6a763a
18f458b7-f4a0-492a-b497-944d6a7c7c78
f6f6d908-7584-432b-98a3-03f78e705b58
55e231e7-b675-4388-902b-118eae1d88c5

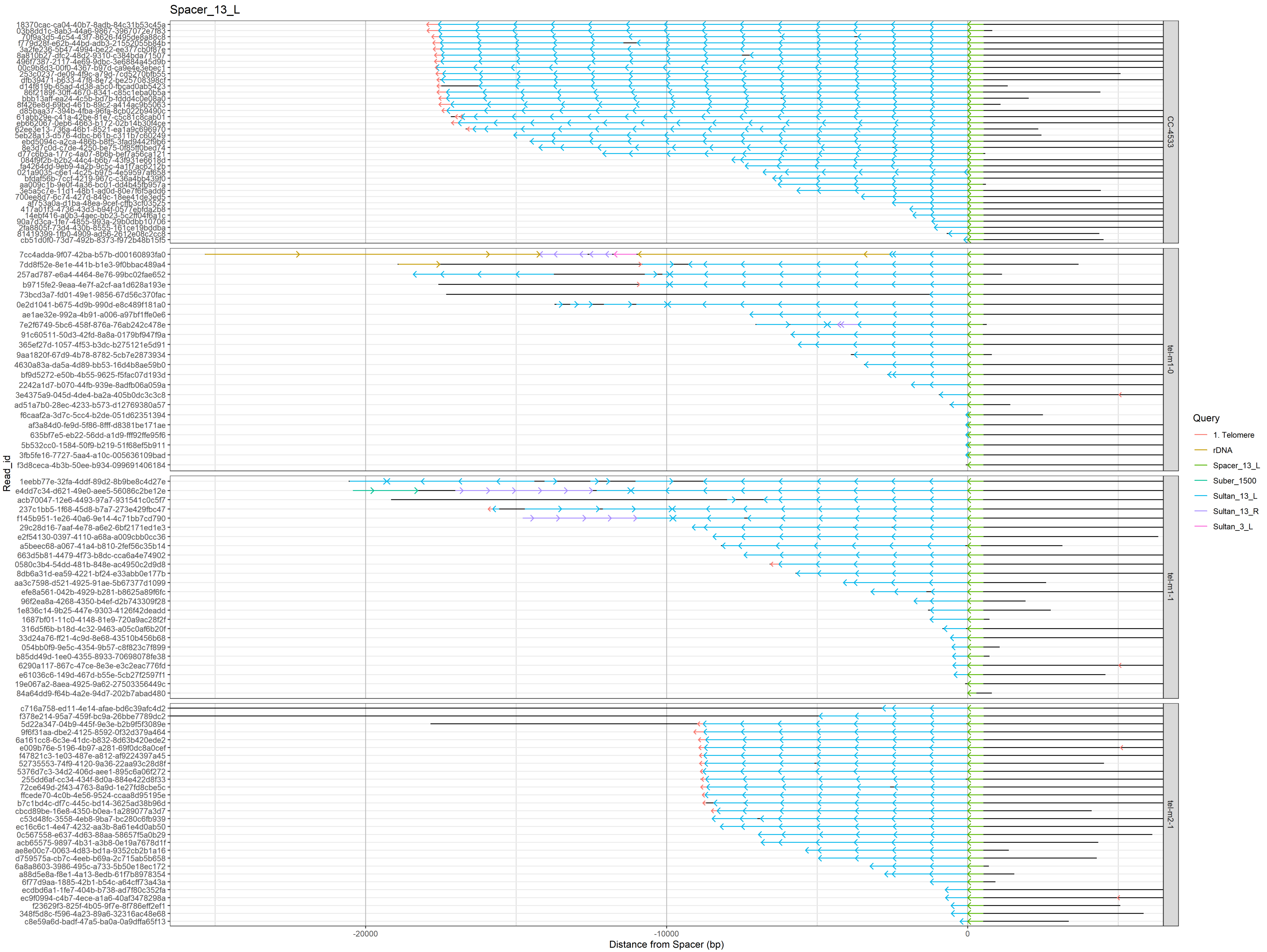

Spacer_13_L
CC-4533
tel-m1-0
tel-m1-1
tel-m2-1
Read_id
-20000
-10000
0
Distance from Spacer (bp)
Query
1. Telomere
rDNA
Spacer_13_L
Suber_1500
Sultan_13_L
Sultan_13_R
Sultan_3_L

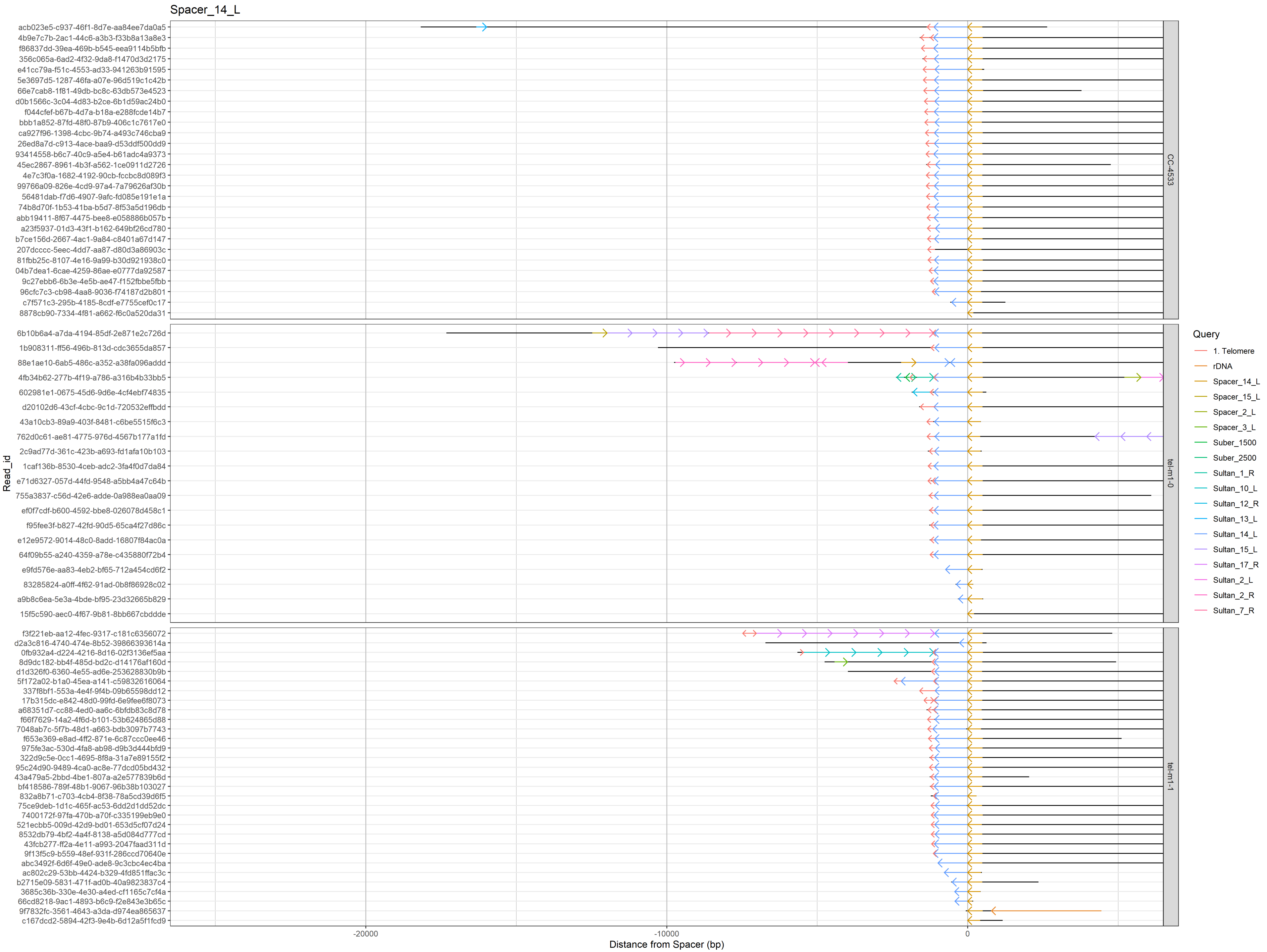

Spacer_14_L
acb023e5-c937-46f1-8d7e-aa84ee7da0a5
4b9e7c7b-2ac1-44c6-a3b3-f33b8a13a8e3
f86837dd-39ea-496b-b545-eea9114b5bfb
356c065a-6ad2-4f32-9da8-f1470d3d2175
e41cc79a-f51c-4553-ad33-941263b91595
5e3697d5-1287-46fa-a07e-96d519c1c42b
66e7cab8-1f81-49db-bc8c-63db573e4523
d0b1566c-3c04-4d83-b2ce-6b1d59ac24b0
f044cfef-b67b-4d7a-b18a-e288fcde14b7
bbb1a852-87fd-48f0-87b9-406c1c7617e0
ca927f96-1398-4cbc-9b74-a493c746cba9
26ed8a7d-c913-4ace-baa9-d53ddf500dd9
93414558-b6c7-40c9-a5e4-b61adc4a9373
45ec2867-8961-4b3f-a562-1ce0911d2726
4e7c3f0a-1682-4192-90cb-fccbc8d089f3
99766a09-826e-4cd9-97a4-7a79626af30b
56481dab-f7d6-4907-9afc-fd085e191e1a
74b8d70f-1b53-41ba-b5d7-8f53a5d196db
abb19411-8f67-4475-bee8-e058886b057b
a23f5937-01d3-43f1-b162-649bf26cd780
b7ce156d-2667-4ac1-9a84-c8401a67d147
207dcccc-5eec-4dd7-aa87-d80d3a86903c
81fbb25c-8107-4e16-9a99-b30d921938c0
04b7dea1-6cae-4259-86ae-e0777da92587
9c27ebb6-6b3e-4e5b-ae47-f152fbbe5fbb
96cfc7c3-cb98-4aa8-9036-f74187d2b801
c7f571c3-295b-4185-8cdf-e7755cef0c17
8878cb90-7334-4f81-a662-f6c0a520da31
CC-4533
6b10b6a4-a7da-4194-85df-2e871e2c726d
1b908311-ff56-496b-813d-cdc3655da857
88e1ae10-6ab5-486c-a352-a38fa096addd
4fb34b62-277b-4f19-a786-a316b4b33bb5
602981e1-0675-45d6-9d6e-4cf4ebf74835
d20102d6-43cf-4cbc-9c1d-720532effbdd
43a10cb3-89a9-403f-8481-c6be5515f6c3
762d0c61-ae81-4775-976d-4567b177a1fd
2c9ad77d-361c-423b-a693-fd1afa10b103
1caf136b-8530-4ceb-adc2-3fa4f0d7da84
e71d6327-057d-44fd-9548-a5bb4a47c64b
755a3837-c56d-42e6-adde-0a988ea0aa09
ef0f7cdf-b600-4592-bbe8-026078d458c1
f95fee3f-b827-42fd-90d5-65ca4f27d86c
e12e9572-9014-48c0-8add-16807f84ac0a
64f09b55-a240-4359-a78e-c435880f72b4
e9fd576e-aa83-4eb2-bf65-712a454cd6f2
83285824-a0ff-4f62-91ad-0b8f86928c02
a9b8c6ea-5e3a-4bde-bf95-23d32665b829
15f5c590-aec0-4f67-9b81-8bb667cbddde
tel-m1-0
f3f221eb-aa12-4fec-9317-c181c6356072
d2a3c816-4740-474e-8b52-39866393614a
0fb932a4-d224-4216-8d16-02f3136ef5aa
8d9dc182-bb4f-485d-bd2c-d14176af160d
d1d326f0-6360-4e55-ad6e-253628830b9b
5f172a02-b1a0-45ea-a141-c59832616064
337f8bf1-553a-4e4f-9f4b-09b65598d12
17b315dc-e842-48d0-99fd-6e9fee6f8073
a68351d7-cc88-4ed0-aa6c-6bfdb83c8d78
f66f7629-14a2-4f6d-b101-53b624865d88
7048ab7c-5f7b-48d1-a663-bdb3097b7743
f653e369-e8ad-4ff2-871e-6c87ccc0ee46
975fe3ac-530d-4fa8-ab98-d9b3d444bfd9
322d9c5e-0cc1-4695-8f8a-31a7e89155f2
95c24d90-9489-4ca0-ac8e-77dcd05bd432
43a479a5-2bbd-4be1-807a-a2e577839b6d
bf418586-789f-48b1-9067-96b38b103027
832a8b71-c703-4cb4-8f38-78a5cd39d6f5
75ce9deb-1d1c-465f-ac53-6dd2d1dd52dc
7400172f-97fa-470b-a70f-c335199eb9e0
521ecbb5-009d-42d9-bd01-653d5cf07d24
8532db79-4bf2-4a4f-8138-a5d084d777cd
43fcb277-ff2a-4e11-a993-2047faad311d
9f13f5c9-b559-48ef-931f-286ccd70640e
abc3492f-6d6f-49e0-ade8-9c3cbc4ec4ba
ac802c29-53bb-4424-b329-4fd851ffac3c
b2715e09-5831-471f-ad0b-40a9823837c4
3685c36b-330e-4e30-a4ed-cf1165c7cf4a
66cd8218-9ac1-4893-b6c9-f2e843e3b65c
9f7832fc-3561-4643-a3da-d974ea865637
c167dcd2-5894-42f3-9e4b-6d12a5f1fcd9
tel-m1-1
Read_id
-20000
-10000
0
Distance from Spacer (bp)
Query
1. Telomere
rDNA
Spacer_14_L
Spacer_15_L
Spacer_2_L
Spacer_3_L
Suber_1500
Suber_2500
Sultan_1_R
Sultan_10_L
Sultan_12_R
Sultan_13_L
Sultan_14_L
Sultan_15_L
Sultan_17_R
Sultan_2_L
Sultan_2_R
Sultan_7_R

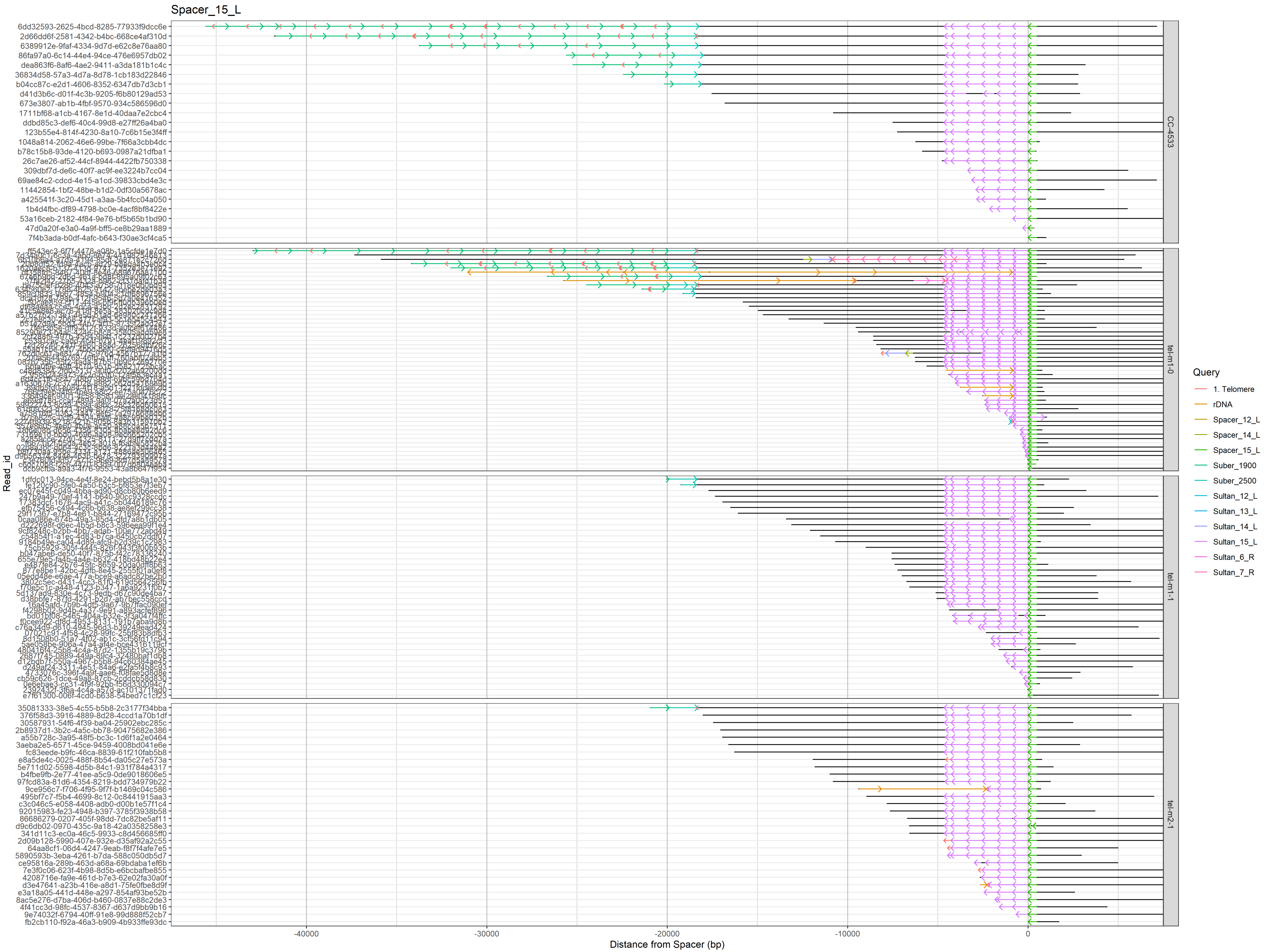

Spacer_15_L
Read_id
Distance from Spacer (bp)
-40000
-30000
-20000
-10000
0
CC-4533
tel-m1-0
tel-m1-1
tel-m2-1
Query
1. Telomere
rDNA
Spacer_12_L
Spacer_14_L
Spacer_15_L
Suber_1900
Suber_2500
Sultan_12_L
Sultan_13_L
Sultan_14_L
Sultan_15_L
Sultan_6_R
Sultan_7_R

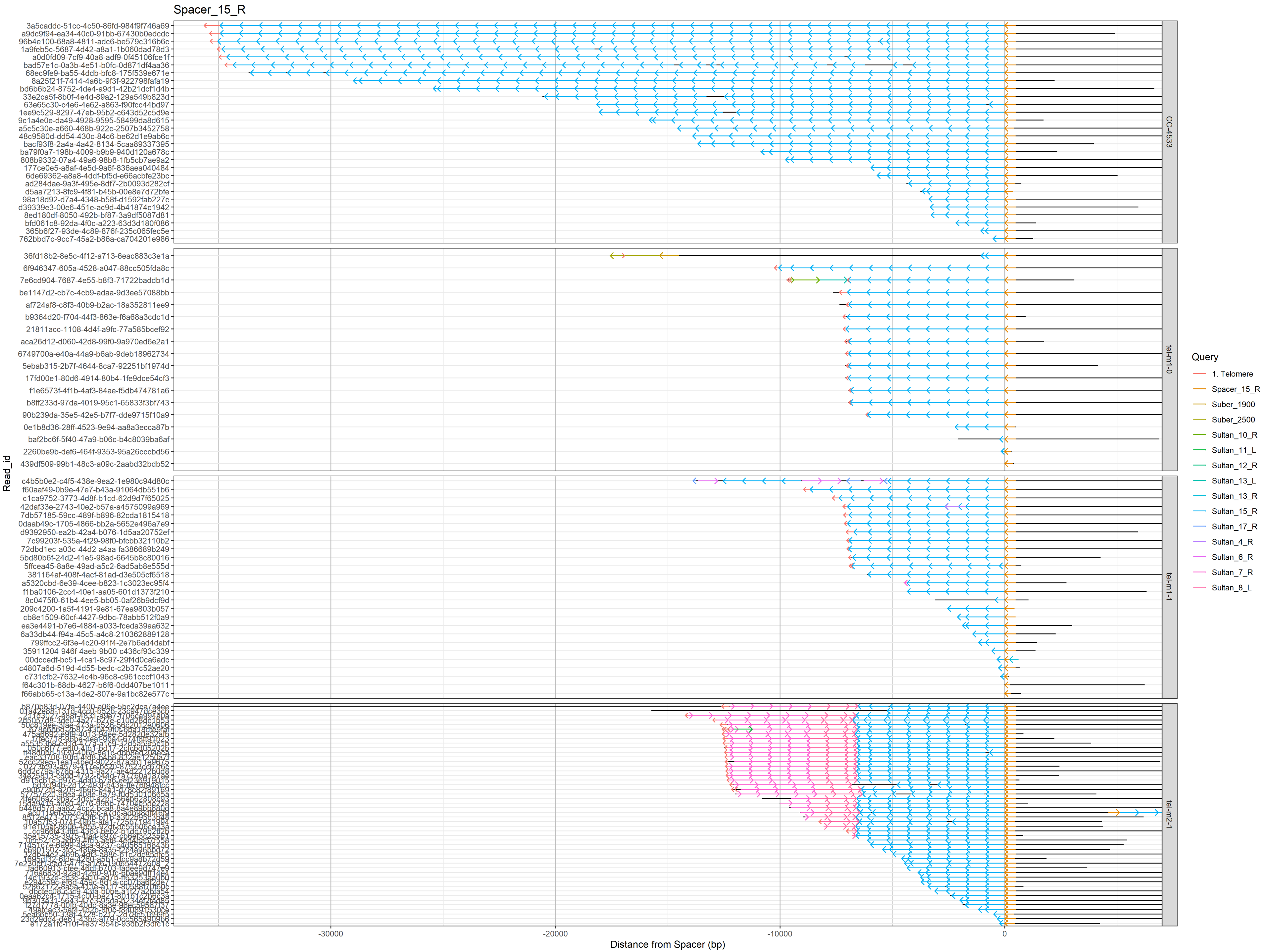
Spacer_15_R
Read_id
Distance from Spacer (bp)
-30000
-20000
-10000
0
CC-4533
tel-m1-0
tel-m1-1
tel-m2-1
Query
1. Telomere
Spacer_15_R
Suber_1900
Suber_2500
Sultan_10_R
Sultan_11_L
Sultan_12_R
Sultan_13_L
Sultan_13_R
Sultan_15_R
Sultan_17_R
Sultan_4_R
Sultan_6_R
Sultan_7_R
Sultan_8_L

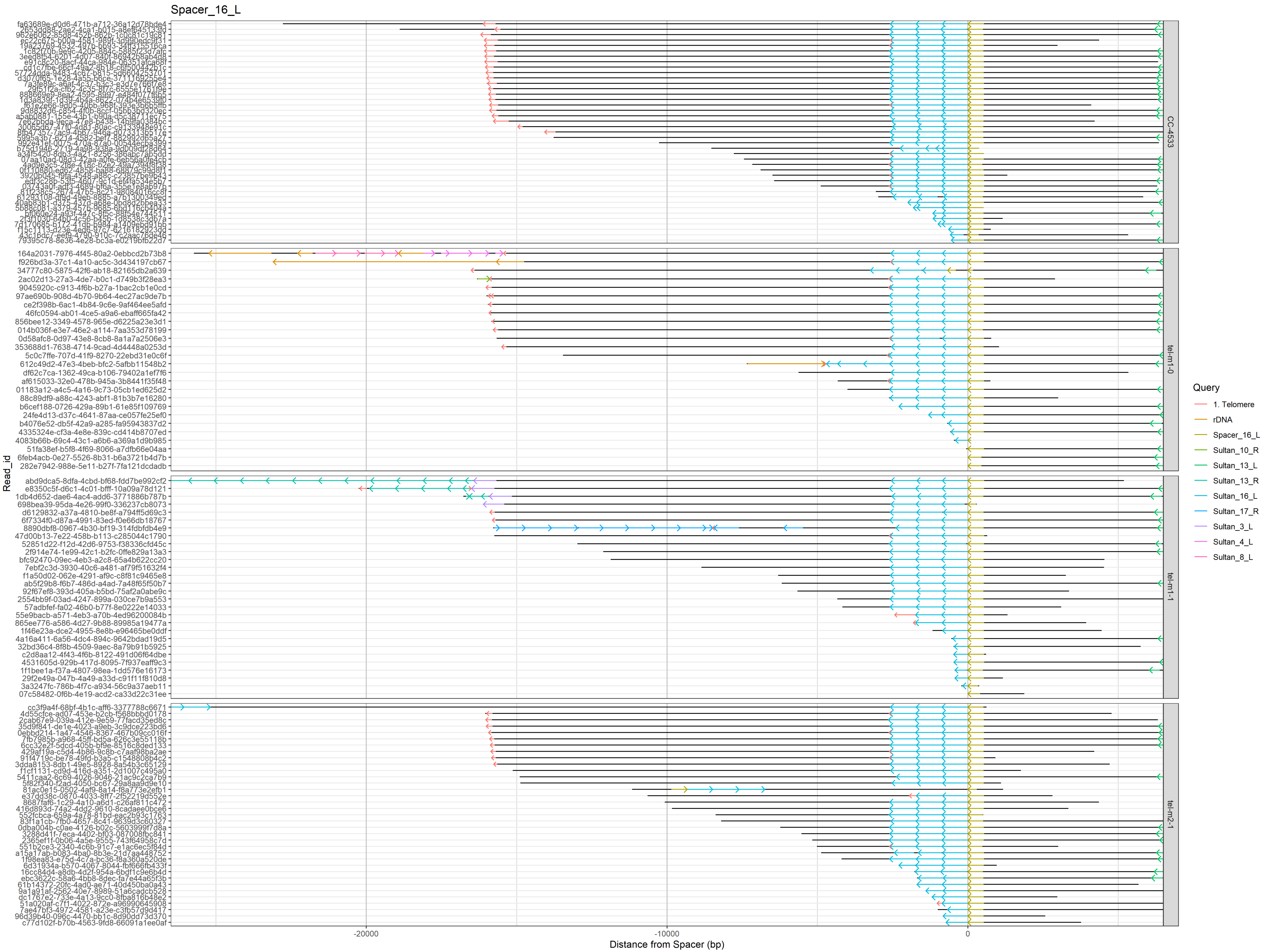

Spacer_16_L
Read_id
Distance from Spacer (bp)
-20000
-10000
0
CC-4533
tel-m1-0
tel-m1-1
tel-m2-1
Query
1. Telomere
rDNA
Spacer_16_L
Sultan_10_R
Sultan_13_L
Sultan_13_R
Sultan_16_L
Sultan_17_R
Sultan_3_L
Sultan_4_L
Sultan_8_L

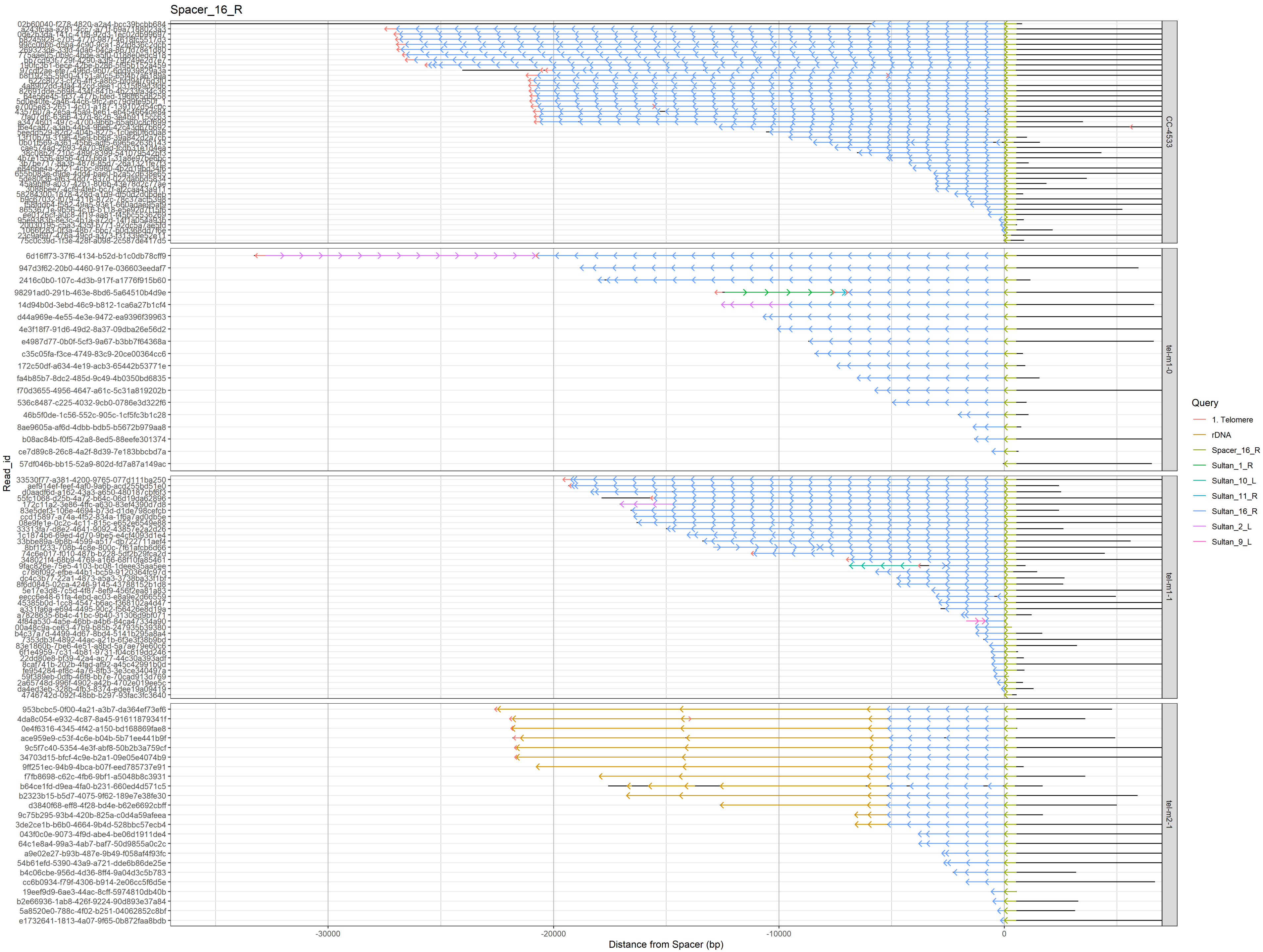

Spacer_16_R
Read_id
Distance from Spacer (bp)
-30000
-20000
-10000
0
CC-4533
tel-m1-0
tel-m1-1
tel-m2-1
Query
1. Telomere
rDNA
Spacer_16_R
Sultan_1_R
Sultan_10_L
Sultan_11_R
Sultan_16_R
Sultan_2_L
Sultan_9_L
6d16ff73-37f6-4134-b52d-b1c0db78cff9
947d3f62-20b0-4460-917e-036603eedaf7
2416c0b0-107c-4d3b-917f-a1776f915b60
98291ad0-291b-463e-8bd6-5a64510b4d9e
14d94b0d-3ebd-46c9-b812-1ca6a27b1cf4
d44a969e-4e55-4e3e-9472-ea9396f39963
4e3f18f7-91d6-49d2-8a37-09dba26e56d2
e4987d77-0b0f-5cf3-9a67-b3bb7f64368a
c35c05fa-f3ce-4749-83c9-20ce00364cc6
172c50df-a634-4e19-acb3-65442b53771e
fa4b85b7-8dc2-485d-9c49-4b0350bd6835
f70d3655-4956-4647-a61c-5c31a819202b
536c8487-c225-4032-9cb0-0786e3d322f6
46b5f0de-1c56-552c-905c-1cf5fc3b1c28
8ae9605a-af6d-4dbb-bdb5-b5672b979aa8
b08ac84b-f0f5-42a8-8ed5-88eefe301374
ce7d89c8-26c8-4a2f-8d39-7e183bbcbd7a
57df046b-bb15-52a9-802d-fd7a87a149ac
953bcbc5-0f00-4a21-a3b7-da364ef73ef6
4da8c054-e932-4c87-8a45-91611879341f
0e4f6316-4345-4f42-a150-bd168869fae8
ace959e9-c53f-4c6e-b04b-5b71ee441b9f
9c5f7c40-5354-4e3f-abf8-50b2b3a759cf
34703d15-bfcf-4c9e-b2a1-09e05e4074b9
9ff251ec-94b9-4bca-b07f-eed785737e91
f7fb8698-c62c-4fb6-9bf1-a5048b8c3931
b64ce1fd-d9ea-4fa0-b231-660ed4d571c5
b2323b15-b5d7-4075-9f62-189e7e38fe30
d3840f68-eff8-4f28-bd4e-b62e6692cbff
9c75b295-93b4-420b-825a-c0d4a59afeea
3de2ce1b-b6b0-4664-9b4d-528bbc57ecb4
043f0c0e-9073-4f9d-abe4-be06d1911de4
64c1e8a4-99a3-4ab7-baf7-50d9855a0c2c
a9e02e27-b93b-487e-9b49-f058af4f93fc
54b61efd-5390-43a9-a721-dde6b86de25e
b4c06cbe-956d-4d36-8ff4-9a04d3c5b783
cc6b0934-f79f-4306-b914-2e06cc5f6d5e
19eef9d9-6ae3-44ac-8cff-5974810db40b
b2e66936-1ab8-426f-9224-90d893e37a84
5a8520e0-788c-4f02-b251-04062852c8bf
e1732641-1813-4a07-9f65-0b872faa8bdb

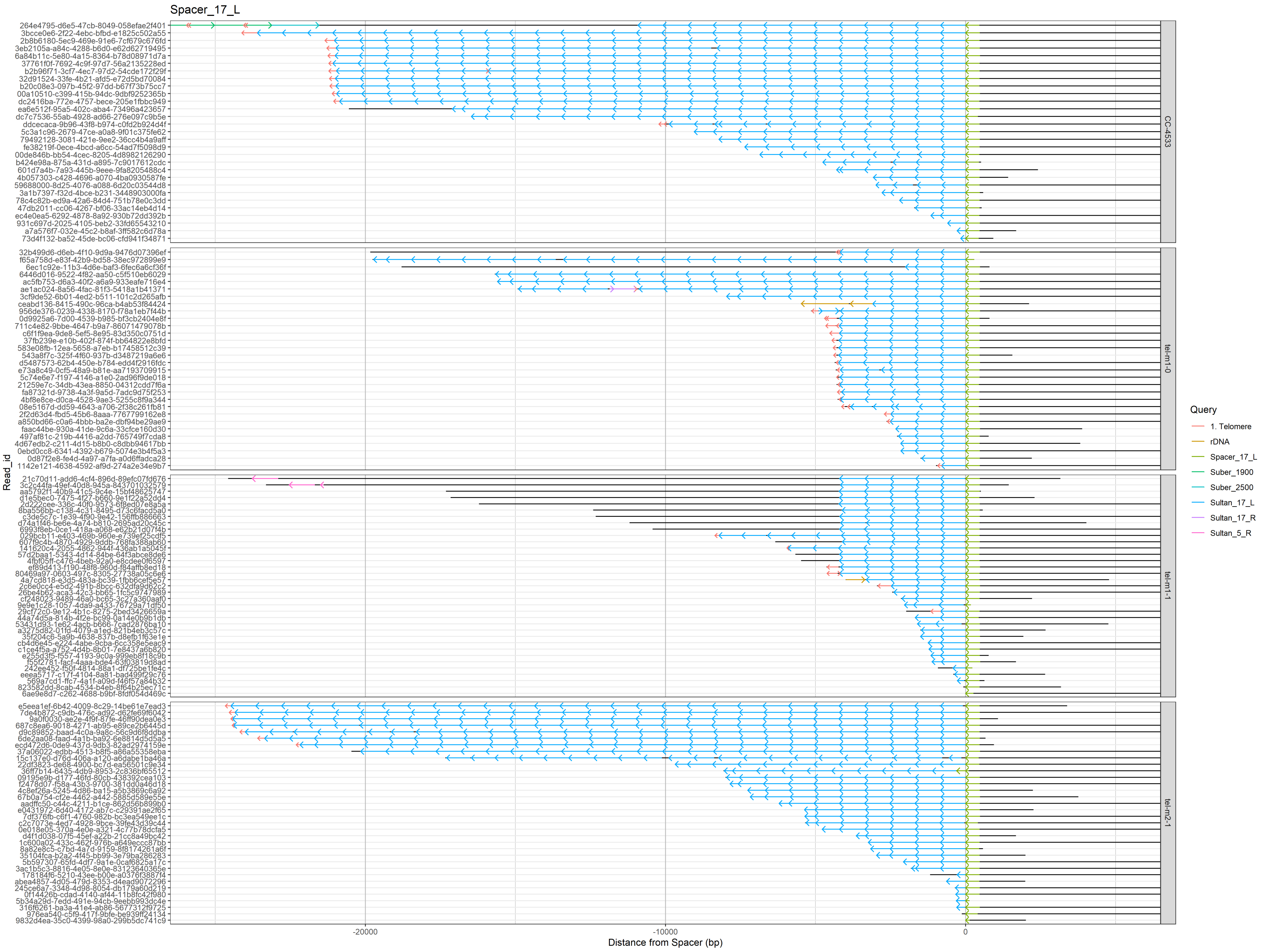
Spacer_17_L
Read_id
Distance from Spacer (bp)
-20000
-10000
0
CC-4533
tel-m1-0
tel-m1-1
tel-m2-1
Query
1. Telomere
rDNA
Spacer_17_L
Suber_1900
Suber_2500
Sultan_17_L
Sultan_17_R
Sultan_5_R

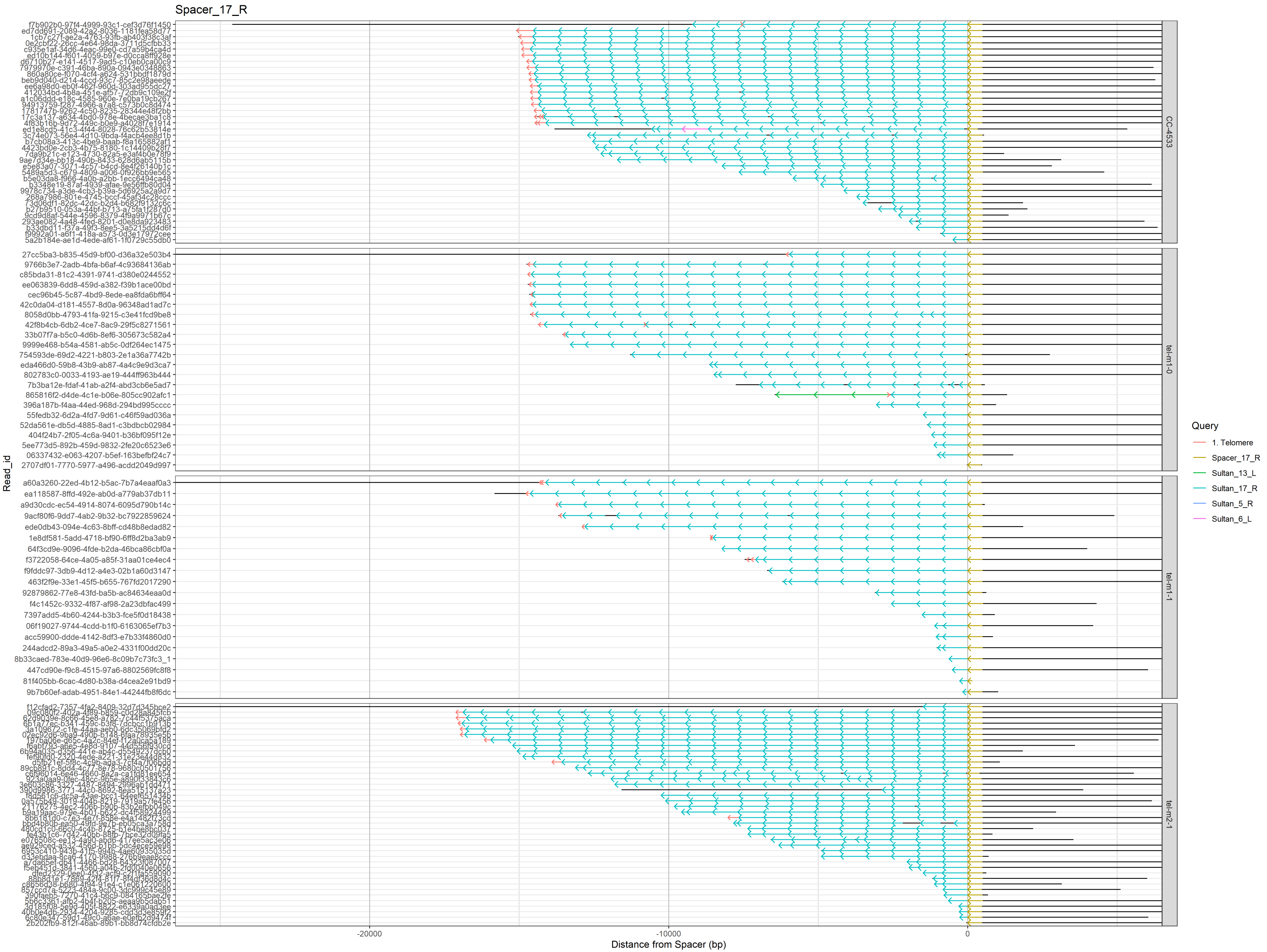

Spacer_17_R
Read_id
Distance from Spacer (bp)
-20000
-10000
0
CC-4533
tel-m1-0
tel-m1-1
tel-m2-1
Query
1. Telomere
Spacer_17_R
Sultan_13_L
Sultan_17_R
Sultan_5_R
Sultan_6_L
